# Chronic chemogenetic slow-wave-sleep enhancement in mice

**DOI:** 10.1101/2025.01.23.634538

**Authors:** Heinrich S. Gompf, Loris L. Ferrari, Greta Pilkauskaite, Christelle Anaclet

## Abstract

While epidemiological associations and brief studies of sleep effects in human disease have been conducted, rigorous long-term studies of sleep manipulations in animal models are needed to establish causation and to understand mechanisms. We have previously developed a mouse model of acute slow-wave-sleep (SWS) enhancement using chemogenetic activation of parafacial zone GABAergic neurons (PZ^GABA^) in the parvicellular reticular formation of the pontine brainstem. However, it was unknown if SWS could be enhanced chronically in this model.

In the present study, mice expressing the chemogenetic receptor hM3Dq in PZ^GABA^ were administered once daily with, sequentially, the chemogenetic ligands, clozapine N-oxide (CNO), deschloroclozapine (DCZ) and compound 21 (C21), and sleep-wake phenotypes were analyzed using electroencephalogram (EEG) and electromyogram (EMG).

We found that SWS time is increased for three hours following the administration of the chemogenetic ligand, and at the same magnitude for at least six months. This phenotype is associated with longer SWS episode duration and an increase of slow wave activity (SWA) of similar magnitude throughout the six-month dosing period. Interestingly, following the end of the six-month dosing period, SWA remains increased for at least a week. In control mice, six-month long daily administration of the chemogenetic ligands did not affect SWS quantity or quality.

This study validates a mouse model of chronic SWS enhancement that will allow mechanistic investigations into how SWS promotes physiological function and prevents diseases. The approach of a rotating schedule of three chemogenetic ligands may be broadly applicable in chemogenetic studies that require chronic administration.

## Introduction

A growing number of studies show that long-term sleep disruption affects a wide array of physiological functions and is associated with numerous chronic conditions such as Alzheimer’s disease (Minakawa et al., 2019; Shen et al., 2016). From this it is inferred that longer-term sleep promotion may deliver disease-ameliorating advantages particularly prior to disease onset. While there is little doubt that improved sleep is an appealing recommendation for healthy aging, there is currently very little causal mechanistic evidence to support this. Animal models of disease and disease resilience or avoidance are crucial in filling these gaps. Before advancing to the translational and clinical stages of investigating this treatment avenue, we need to consider whether sleep enhancement in general, and specific enhancement of particular stages of sleep such as slow-wave-sleep (SWS), will provide beneficial effects over the long term. We have previously developed a mouse model of SWS enhancement which increases and enhances SWS by acute chemogenetic activation of parafacial zone GABAergic neurons (PZ^GABA^) (Anaclet et al., 2014). We have also previously shown that the oral route of chemogenetic agonist administration is as effective as the traditional intraperitoneal injections (Ferrari et al., 2022). Here we present a daily long-term SWS enhancement.

In mice, two major sleep stages are conventionally distinguished, non-rapid eye movement sleep (NREMS) also called SWS and rapid eye movement sleep (REMS). In humans, NREMS is further subdivided into three stages. Stage 1 is drowsiness, stage 2 is light sleep and stage 3 is deep sleep, also called SWS. SWS is characterized by high amplitude slow waves at the delta (0.5-4 Hz) frequency band, also called slow wave activity (SWA). Of NREMS stages, SWS has particular importance. Specifically, SWS has been shown to be involved in clearing the metabolites accumulated during wakefulness and intense neuronal activity (Grubb & Lauritzen, 2019), and is important for the processing and consolidation of learning (Malkani & Zee, 2022). As relates to metabolites, a significant number of animal studies provide evidence that the pathological hallmarks of Alzheimer’s disease, amyloid β and tau protein, accumulate in the interstitial fluid during wakefulness and are cleared acutely during SWS (Holth et al., 2019). Moreover, it is during SWS, synaptic remodeling—downscaling (Tononi & Cirelli, 2020), upscaling (Kopp et al., 2006) and synaptic fidelity tuning (Angelakos et al., 2017)—also occurs. This suggests that long-term sleep disruption will both accelerate the accumulation of the amyloid β and tau protein in the brain, as well as compromise memory ensemble integrity, thereby both increasing the risk of developing Alzheimer’s disease and decreasing the resilience against its neurodegenerative consequences. Extrapolating from acute sleep deprivation studies and epidemiological evidence suggesting chronic sleep deprivation during mid-life increases the risk of developing Alzheimer’s, it has been hypothesized that chronic sleep enhancement can prevent neurodegeneration and promote cognitive function. Similarly, chronic deficits of sleep have been involved in other disorders, such as epilepsy (Garg et al., 2022), cardiovascular disorders (Chasens et al., 2021), cognitive decline (Dzierzewski et al., 2022) and others (Shen et al., 2016), and sleep promotion has been suggested as a potential interventional strategy. To test this hypothesis and study the underlying mechanisms, it was first necessary to develop an animal model of chronic sleep enhancement.

The effect of chronic dosing with chemogenetic ligands was never previously investigated in sleep-wake studies. In addition, a number of chemogenetic studies have found either antagonistic or null effects of repeated and/or chronic dosage of chemogenetic agonists, possibly due to receptor desensitization or neuroadaptive changes (Claes et al., 2022). Finally, a study showed abnormal gut microbiota composition following seven-day chronic dosing of clozapine N-oxide (CNO) or compound 21 (C21) in drinking water (Guo et al., 2021). Most of these studies used one chemogenetic ligand and did not control for accumulation of the metabolites over time. Therefore, in this study, to limit metabolite accumulation, we used three previously validated chemogenetic ligands: CNO, deschloroclozapine (DCZ) and C21 (Ferrari et al., 2022), administrated acutely and sequentially (day 1: CNO, day 2: DCZ and day 3: C21, day 4: CNO, etc…). This was to limit the accumulation of any metabolite specific to either CNO, DCZ or C21. CNO is converted to clozapine which is further converted in N-desmethylclozapine (NDMC). CNO and clozapine are undetecteble in the CSF 24-hr following administration of CNO (3 mg/kg) in monkey (Raper et al., 2017). The same study showed NDMC remains detectable in the plasma for up to 48-hr following CNO (10 mg/kg) administration though plasma levels decrease rapidly earlier than that. DCZ metabolites are C21 and DCZ-N-oxide. These three componds are undetecteble in the mouse brain 24-hr following DCZ (0.1 mg/kg) (Nagai et al., 2020). C21 metabolites are unknown and C21 clearance from the brain apprears to be longer than the other chemogenetic ligands and their metabolites (Jendryka et al., 2019). Therefore, we decided on an experimental paradigm of a three-day wash-out period between each chemogenetic ligand to limit accumulation of metabolic products. In summary, we show that chemogenetic activation of PZ^GABA^ results in SWS enhancement of the same magnitude, daily, over at least 6 months and has no sleep-wake effects in control mice.

## Methods

### Animals

A total of 12 pathogen-free mice, on the C57BL/6J genetic background, were used in this study. The mice resulted from the cross of the following mouse lines: *Vgat*-IRES-Cre (S*lc32a1tm2(cre)Lowl*/J, Jackson stock #028862), *EGFP-L10A* (Jackson stock #024750) and B6.CgTg (APPswe,PSEN1dE9)85Dbo/Mmjax (Jackson stock #5864). The study includes a SWS enhanced group (PZ^Vgat-hM3Dq^ mice) (Anaclet et al., 2014) and a control group. The SWS enhanced group includes 3 males and 3 females, 3 Vgat-Cre+/GFP/APP,PS1+ mice and 3 Vgat-Cre+/GFP/APP,PS1- mice. The control group includes 5 males and 1 female, 1 Vgat-Cre+/GFP/APP,PS1+ mouse and 5 Vgat-Cre+/GFP/APP,PS1- mice. Mice were bred at our animal facility and underwent genotyping both before and after experiments, as previously described (Jankowsky et al., 2005; Vong et al., 2011). Care of these animals met the National Institutes of Health standards, as set forth in the *Guide for the Care and Use of Laboratory Animals* and all protocols were approved by the University of Massachusetts Chan Medical School and the University of California Davis Institutional Animal Care and Use Committees.

### Surgery

PZ^Vgat-hM3Dq^ group: At the age of 8 weeks, naïve mice were subjected to two independent surgeries separated by four weeks. Mice were anesthetized with ketamine/xylazine [100 and 10 mg/kg, respectively, intraperitoneally (IP)] and then placed in a stereotaxic apparatus. During the first surgery mice received bilateral injections of an adeno-associated viral (AAV; serotype 2) vector expressing the hM3Dq receptor and mCherry (reporter gene) in a cre-dependent configuration (hSyn-DIO-hM3Dq-mCherry-AAV, UMASS Vector Core, titer: 3.0E+12 Viral particles/mL) into the PZ, as previously described (Anaclet *et al*., 2014). Coordinates from Bregma were −5.6 mm antero-posterior, ± 1.0 mm lateral, −4.2 mm dorso-ventral, as per the mouse atlas of Paxinos and Franklin (Paxinos & Franklin, 2001). The AAV (200 nl) was injected into the PZ of mice using a 10 µl Hamilton syringe (Hamilton Co., Reno, NV) at a rate of 3 nl/sec driven by an UMP2 microinfusion pump with a SMARTouch Controller (World Precision Instruments, Inc., Sarasota, Florida). During the second surgery, mice were implanted with four EEG screw electrodes (2 frontal [1 mm frontal, 1 mm lateral from bregma] and 2 parietal [mid-distance between bregma and lambda and 1 mm lateral from the mid-line] electrodes (Pinnacle Technology Inc., United States, Catalog #8403) and two flexible electromyogram (EMG) wire electrodes in the neck muscles (Protech International, United States, catalog #E363/76/SPC), previously soldered to a 6-pin connector (Heilind Electronics, United States, catalog #853-43-006-10- 001000) and the assembly was secured to the skull with dental cement. After completing the surgery, mice were kept in a warm environment until resuming normal activity as previously described (Anaclet et al., 2014). The mice in the control group were subjected to same surgery protocol for EEG/EMG implantation.

### Sleep-Wake Recording

Following the second surgery, the mice were housed individually in transparent barrels in an insulated sound-proofed recording chamber maintained at an ambient temperature of 22 ± 1 ◦C and on a 12 h light/dark cycle (lights-on at 07:00, Zeitgeber time: ZT0) with food and water available *ad libitum*. Following 10 days for post-surgical recovery, mice were connected to flexible recording cables and habituated to the recording conditions for 5 days before starting polygraphic recording. One cortical EEG (bipolar, fronto-parietal, ipsilateral) and the EMG signals were amplified (A-M System 3500, United States) and digitalized with a resolution of 256 Hz using Vital Recorder (Biopac Systems Inc, United States).

### Drug Administration

During the habituation period, mice were trained to the voluntary oral administration (Jelly) (Ferrari et al., 2022; Mahoney et al., 2019). Vehicle Jelly (control) was prepared as follows: gelatine (7 g, original unflavored gelatin, Knox) and Splenda (10 g) were mixed in 49 ml ddH2O + 1 ml natural food flavor (strawberry flavor, Frontier Co-op, United States) and stirred at 50°C until the mix was clear. The stock solution was then placed in 4°C for 1-2 months of storage. To prepare the jellies, the stock solution was stirred at 50°C until completely melted, permitting dilution of the drugs. Then the appropriate volume of stock solution (control-Jelly) or of stock solution containing clozapine N-oxide (CNO, 0.3 mg/kg, CNO-Jelly), deschloroclozapine (DCZ, 0.5 mg/kg, DCZ-Jelly) or compound 21 (C21, 3 mg/kg, C21-Jelly), was pipetted into the cap of an Eppendorf tube (0.05 ml / 10 g of mouse) and placed at 4°C until it forms a jelly again. Before giving to the mice, the Jelly was removed from the cap of the Eppendorf tube and placed in the cap of a 15 ml falcon tube which was placed in the mouse cage. In order to train the mice to eat the Jelly, mice were food deprived overnight (12-14 hr) before the first control-Jelly presentation. The following morning, mice were given control-Jelly, placed in a 15 ml falcon tube cap and on the floor of the cage. Mice were left without food until they ate the Jelly (5-10 min). Once the mice had eaten the jelly, they were given regular chow ad libitum. Mice were then given control-Jelly daily without food deprivation. After a 5-day habituation period, the mice were eating the jelly within < 1 min following presentation. CNO, DCZ and C21 were obtained from the NIMH Chemical Synthesis and Drug Supply Program.

### Experimental timeline

Following the 24-hr baseline recording period, the mice were given a control-Jelly at 10:00 [Zeitgeber time 3 (ZT3)]. Then the mice were given CNO-Jelly (0.3 mg/kg), DCZ-Jelly (0.5 mg/kg) or C21-Jelly (3 mg/kg), daily (ZT3) for a 6-month period. At the end of the drug treatment, mice were recorded for an additional 7-day washout period (**Fig. 1**). The mice remained connected to the recording cable and were recorded for 2 weeks per month and were left free from the cable for the rest of the month to relieve them from the cable constraints. Cable connection/disconnection as well as the weekly cage change were performed between ZT11-12, when the mice show an increase in wake time in anticipation of the active period, to minimize the sleep-wake disruptions.

**Figure 1:**
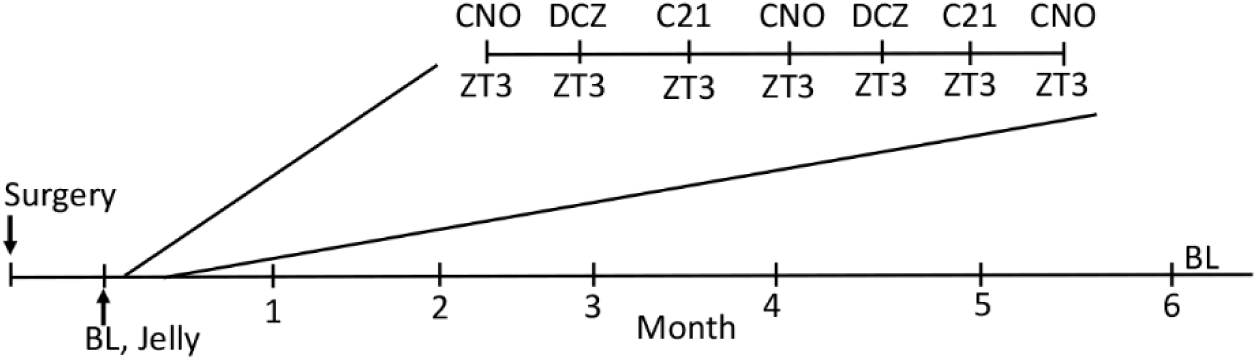
Experimental timeline. Two weeks after surgery, the mice were recorded in the baseline condition (BL, 48-hr) followed by control administration of Jelly at ZT3. Then the mice received daily administration of CNO, DCZ or C21, in Jelly, at ZT3 for 6 months. At the end of the administration period, the mice were recorded for 7-day wash-out period.

### Sleep Scoring and Analysis

The 24-hr baseline recording, the 24-hr post control-Jelly and the 24-hr post CNO-Jelly, DCZ-Jelly or C21-Jelly from the 2 first administrations of each month and the post 6-month dosing washout period (7 days) were analyzed.

Using SleepSign for Animal (Biopac Systems Inc, United States) assisted by spectral analysis using fast Fourier transform (FFT), polygraphic records were visually scored in 10 s epochs for wakefulness, SWS, and REMS. Wakefulness is characterized by low amplitude fast frequency EEG associated with EMG activity. SWS is characterized by high amplitude, low frequency EEG, and low EMG activity. REMS is characterized by an EEG dominated by hippocampal theta rhythm and no EMG activity (Anaclet et al., 2015). The percentage of time spent in wakefulness, SWS, and REMS were summarized for each group and each condition. The SWS and REMS latencies are defined as the time between the placement of the Jelly on the floor of the cage and the onset of the first SWS episode, lasting >20 s, and the onset of the first REMS episode, lasting >10 s, respectively.

Recordings were scored again in 4 s epochs to allow for performance of the cortical EEG power spectral analysis. Based on visual and spectral analysis, epochs containing artifacts occurring during active wakefulness (with large movements) or containing two vigilance states were visually identified and omitted from the spectral analysis. Re-scoring with a shorter epoch length allows us to provide a greater time resolution and thus minimizes the proportion of total time omitted from the analysis. Recordings containing artifacts during more than 20% of the recorded time were removed from the spectral analysis. Cortical EEG power spectra were computed for consecutive 4 s epochs within the frequency range of 0.5 – 60 Hz using a fast Fourier transform (FFT) routine. The data were collapsed into 0.5 Hz bins. The data were standardized by expressing each frequency bin as a percentage relative to the total power of the same epochs [for example, (bin power * 100)/0.5–60 Hz total power]. To analyze the EEG frequency bands, standardized power bins were summed in delta (δ, 0.5 – 4.5 Hz), theta (θ, 4.5 – 10 Hz), sigma (α, 10 – 15 Hz), beta (β, 15 – 30 Hz), and gamma (γ, 30 – 60 Hz) bands.

### Statistical analysis

Statistical analysis was performed using Prism v10 (GraphPad Software, San Diego, CA, United States). PZ^Vgat-hM3Dq^ group: Following confirmation that the data met the assumptions of the ANOVA model, 1) two-way ANOVA for repeated measures followed by a post hoc Bonferroni test was used to compare the effect of the length of drug dosing on sleep-wake amount and power bands; and 2) one-way ANOVA for repeated measures followed by a post hoc Bonferroni test was used to compare the effect of the length of drug dosing on 24-hr sleep-wake amounts as well as on the SWS and REMS latencies. To compare the power band distribution between the baseline day before the chronic treatment and the 7^th^ day of the washout period following the chronic treatment, a mixed-effect analysis for repeated measures followed by the two-stage linear step-up procedure of Benjamini, Krieger and Yekutieli was used, due to missing individual data points. Control group: one mouse died 1 month before the end of the study and one mouse EEG became contaminated during the 6^th^ month recording, resulting in N = 5 in the 6^th^ month sleep-wake amount data and N = 4 in the 6^th^ month power spectral analysis of the EEG. Therefore, a mixed-effect analysis for repeated measures followed by Bonferroni test was used.

## Results

### Daily sleep-wake proportions

To examine whether repeated daily oral administration of chemogenetic agonists produces antagonistic or null effects on sleep measures over time, sleep/wake stages were quantified for the first two 24-hour periods of each month in PZ^Vgat-hM3Dq^ mice. The daily percentage of wakefulness following administration was not significantly different compared with control administration, with the exception of a significant decrease at the 1st month DCZ administration (**Fig. 2A1-C1**). However, the one-way ANOVA shows a significant effect of the 6-month daily drug dosing of CNO [F (7, 35) = 3.844, P = 0.0034, **Fig. 2A1**] and DCZ [F (7, 35) = 2.698, P = 0.0240, **Fig. 2B1**], but not C21 [F (7, 35) = 2.242, P = 0.0540, **Fig. 2C1**]. The daily 6-month drug dosing also had a significant effect on the percentage of daily SWS following each chemogenetic agonist administration: CNO [F (7, 35) = 4.948, P = 0.0006, **Fig. 2A2**], DCZ [F (7, 35) = 3.041, P = 0.0132, **Fig. 2B2**] and C21 [F (7, 35) = 10.30, P < 0.0001, **Fig. 2C2**]. The daily percentage of SWS was significantly increased following administration of CNO in the 1^st^, the 5^th^ and the 6^th^ month, and following administration of DCZ and C21 in the 1^st^ month compared with control administration (**Fig. 2A2-C2**). The 6-month daily treatment significantly affected the daily percentage of REMS: CNO [F (7, 35) = 9.902, P < 0.0001, **Fig. 2A3**], DCZ [F (7, 35) = 3.324, P = 0.0081, **Fig. 2B3**] and C21 [F (7, 35) = 11.58, P < 0.0001, **Fig. 2C3**]. The daily REMS percentages were significantly reduced compared with control administration except following the 5^th^ month administration of CNO and the 1^st^ two administrations, the 1^st^ month, the 5^th^ month and the 6^th^ month administration of DCZ (**Fig. 2A3, B3, C3**).

**Figure 2:**
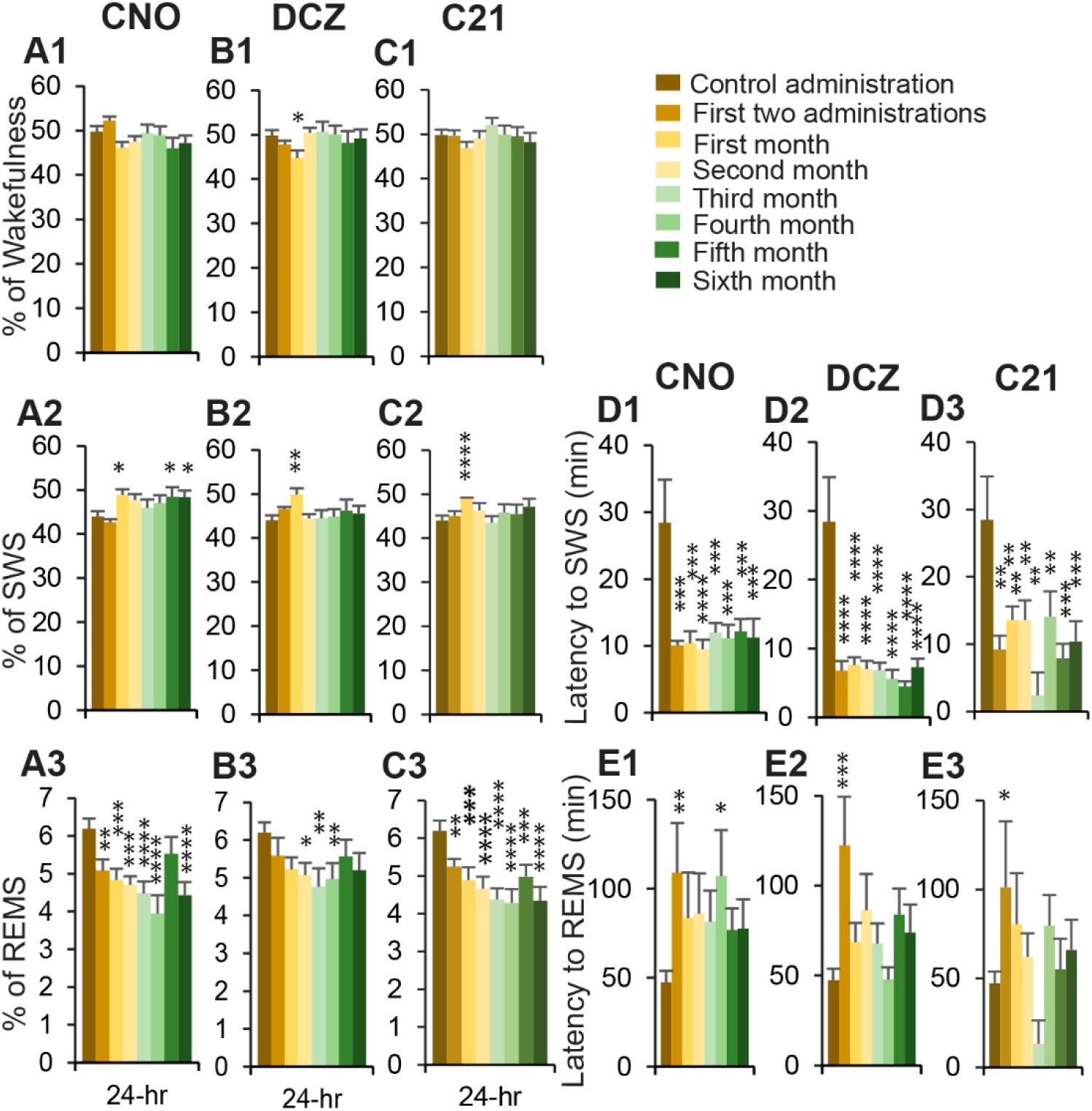
24-hr sleep-wake percentages and sleep latencies in PZ^Vgat-hM3Dq^ mice. (**A-C**) Daily percentage (± SEM) of wakefulness (**A1,B1,C1**), SWS (**A2,B2,C2**) and REMS (**A3,B3,C3**) following administration of CNO (**A**), DCZ (**B**) or C21 (**C**) and from the two first administration of each month. (**D-E**) Latency to SWS (**D**) and to REMS (**E**) following placement of the CNO-Jelly (**D1,E1**), DCZ-Jelly (**D2,E2**) or C21-Jelly (**D3,E3**) in the cage. N = 6 mice. Significant differences compared with control administration, * p < 0.05, ** p < 0.01, *** p < 0.001, **** p < 0.0001, one-way ANOVA followed by Bonferroni test.

**Figure 3:**
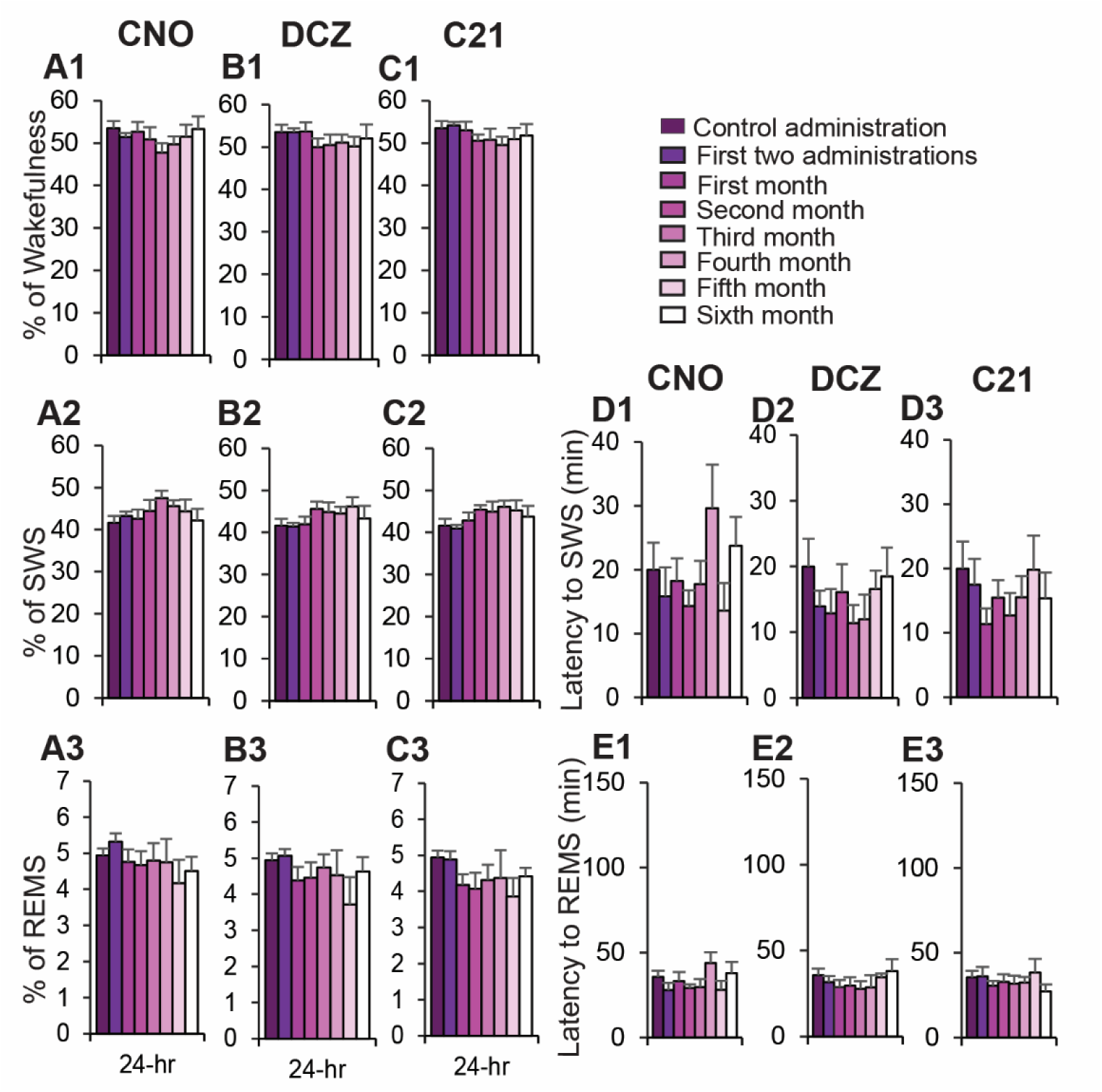
24-hr sleep-wake percentages and sleep latencies in control mice. (**A-C**) Daily percentage (± SEM) of wakefulness (**A1,B1,C1**), SWS (**A2,B2,C2**) and REMS (**A3,B3,C3**) following administration of CNO (**A**), DCZ (**B**) or C21 (**C**) and from the two first administration of each month. (**D-E**) Latency to SWS (**D**) and to REMS (**E**) following placement of the CNO-Jelly (**D1,E1**), DCZ-Jelly (**D2,E2**) or C21-Jelly (**D3,E3**) in the cage. BL to 5^th^ month: N = 6 mice; 6^th^ month: N = 5 mice. No significant differences compared with control administration, mixed-effect analysis for multiple comparison followed by Bonferroni test.

In control mice, the 6-month administration period had no significant effect on the percentage of daily wakefulness following each chemogenetic agonist administration: CNO [F (2.532, 11.94) = 1.236, P = 0.3344, **Fig. 3A1**], DCZ [F (2.318, 11.26) = 1.278, P = 0.3215, **Fig. 3B1**] and C21 [F (2.923, 14.20) = 0.8614, P = 0.4812, **Fig. 3C1**]. Similarly, the daily percentage of SWS was not affected by the 6-month daily drug dosing: CNO [F (2.561, 12.07) = 1.464, P = 2725, **Fig. 3A2**], DCZ [F (2.268, 11.02) = 2.102, P = 0.1657, **Fig. 3B2**] and C21 [F (2.834, 13.77) = 1.487, P = 0.2620, **Fig. 3C2**]. Finally, a similar lack of effect was seen on daily REMS percentages: CNO [F (2.254, 10.62) = 1.524, P = 0.2633, **Fig. 3A3**], DCZ [F (2.613, 12.69) = 1.308, P = 3117, **Fig. 3B3**] and C21 [F (2.141, 10.40) = 1.133, P = 3337, **Fig. 2C3**]. Each month, the daily proportion of the three vigilance stages were not significantly deferent as compared with control administration. These data confirm that 6-month long daily administration of the chemogenetic ligands alone, in mice lacking the chemogenetic receptor, do not affect the daily proportion of the three vigilance stages in control mice.

### Sleep latencies

We examined sleep onset latency in PZ^Vgat-hM3Dq^ mice. The 6-month daily drug dosing had a significant effect on SWS latency following administration of CNO [F (7, 35) = 5.565, P = 0.0002, **Fig. 2D1**], DCZ [F (7, 35) = 10.44, P < 0.0001, **Fig. 2D2**] and C21 [F (7, 35) = 4.483, P = 0.0012, **Fig. 2D3**]. SWS latencies were consistently significantly decreased throughout the 6-month treatment as compared with control administration (**Fig. 2D1-3**). This provides support to assert that desensitization is not occurring in our paradigm. The 6-month daily drug dosing had a significant effect on REMS latency when CNO [F (7, 35) = 2.561, P = 0.0306, **Fig. 2E1**] and DCZ [F (7, 35) = 3.752, P = 0.0039, **Fig. 2E2**] were administrated but not C21 [F (7, 35) = 1.969, P = 0.0878, **Fig. 2E3**]. However, multiple comparisons rarely reached significance between individual month of chemogenetic agonist administration and control administration (**Fig. 2E1-3**), probably due to high variability.

These effects were not seen in the control mouse group (**Fig. 3D-E**). The 6-month daily drug dosing had no significant effect on SWS latency following administration of CNO [F (2.282, 11.08) = 2.610, P = 0.1134, **Fig. 3D1**], DCZ [F (2.057, 9.990) = 2.425, P = 0.1377, **Fig. 3D2**] and C21 [F (2.451, 11.90) = 1.778, P = 0.2088, **Fig. 3D3**]. Similarly, the 6-month daily drug dosing had no significant effect on REMS latency when CNO [F (2.050, 9.958) = 2.557, P = 0.1263, **Fig. 3E1**], DCZ [F (2.691, 13.07) = 1.382, P = 0.2908, **Fig. 3E2**] or C21 [F (2.460, 11.95) = 1.030, P = 0.4013, **Fig. 3E3**] were administrated. Multiple comparisons did not show significant different sleep latency values between individual months of chemogenetic agonist administration and control administration (**Fig. 3D1-3 & C1-3**). These results indicate that daily administration of the chemogenetic ligand does not induce habituation for the dosing or non-specific effects on the ability to fall asleep in control mice.

### Ultradian variations in sleep-wake proportions

Sleep-wake stage percentages were significantly affected at 3-hr time windows following drug administration in PZ^Vgat-hM3Dq^ mice compared to control administration (**Fig. 4**). The percentage of wakefulness was significantly decreased during the 3-hr (ZT3-6) period following CNO (**Fig. 4A1**) and C21 (**Fig. 4C1**) administration from the 2^nd^-6^th^ month, but only from the 5^th^-6^th^ month on days when DCZ (**Fig. 4B1**) was administrated. The percentage of SWS was significantly increased during the 3-hr (ZT3-6) following CNO (**Fig. 4A2**) and C21 (**Fig. 4C2**) administration from the 1^st^ two administrations of each drug to the 6^th^ month, but only from month 1-6 following the initial dose of DCZ (**Fig. 4B2**). The data show that not only is desensitization not occurring but rather exhibits a trend towards increasing effect sizes (greater statistical significance compared to control administration) over time. The percentage of REMS was significantly decreased during the entire light period following drug administration (ZT3-6 & ZT6-12; **Fig. 4A3, B3, C3**), from the 1^st^ two administrations of each drug to the 6^th^ month of administration. The REMS inhibition effect showed increased significance during the ZT6-12 period than during the ZT3-6 period (**Fig. 4A3, B3, C3**) but this might be due to the relatively small amount of REMS during the first 3-hr following control administration. No REMS rebound was seen during the following dark period (ZT12-0) or during the 3-hr light period (ZT0-3) preceding the next day administration, as was also the case with acute administration in our sleep enhancement paradigm (Anaclet et al., 2014).

**Figure 4:**
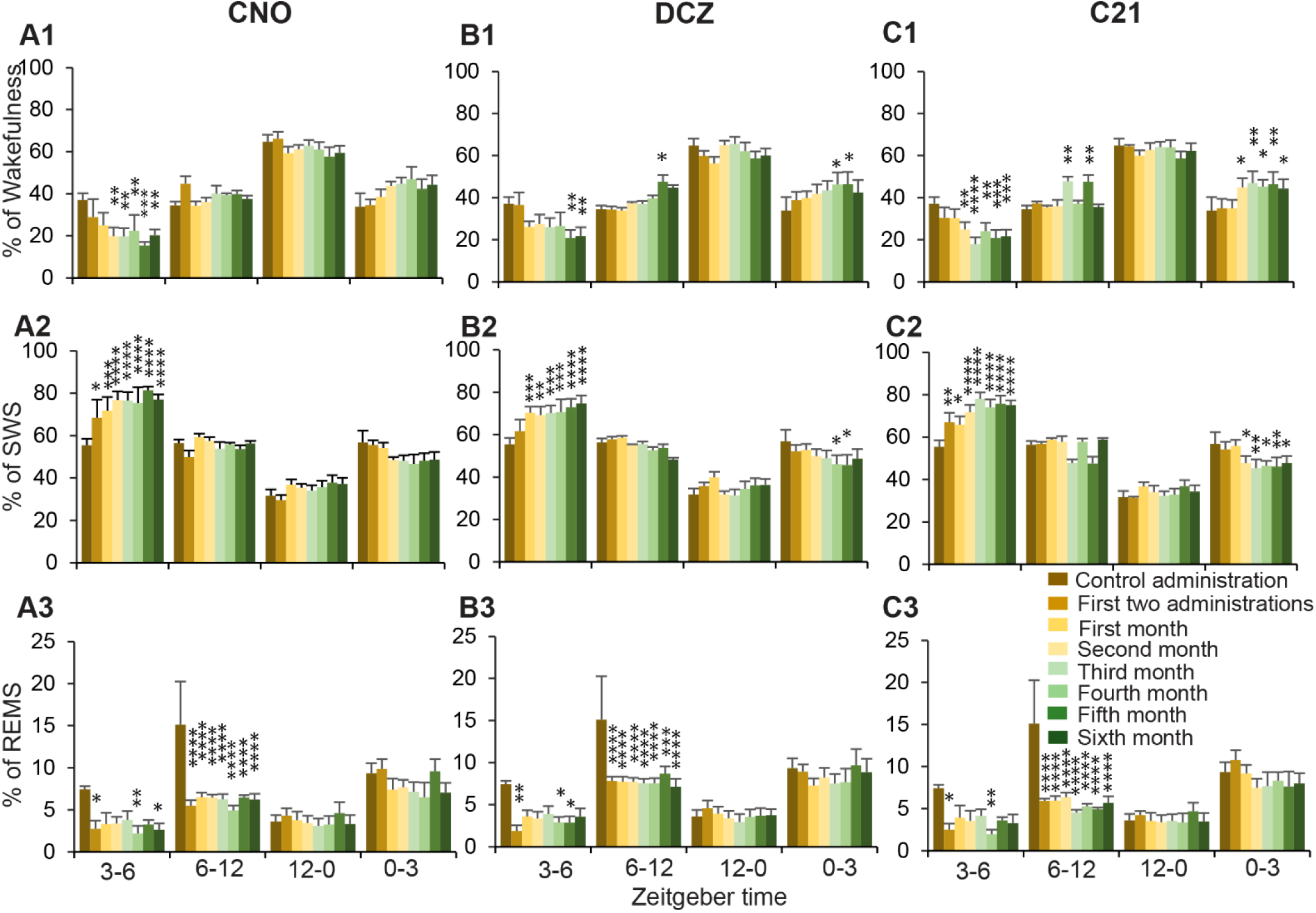
Percentage of sleep-wake states during the 6-month dosing in PZ^Vgat-hM3Dq^ mice. (**A-C**) Percentage (± SEM) of wakefulness (**A1,B1,C1**), SWS (**A2,B2,C2**) and REMS (**A3,B3,C3**) following administration of CNO (**A**), DCZ (**B**) or C21 (**C**) and from the two first administrations of each month, between ZT3-6, ZT6-12, ZT12-0 and ZT0-3. N = 6 mice. Significant differences compared with control administration, * p < 0.05, ** p < 0.01, *** p < 0.001, **** p < 0.0001, two-way ANOVA followed by Bonferroni test.

These effects were mostly not observed in the control group (**Fig. 5**), where the 6-month daily dosing of the chemogenetic agonists did not affect the sleep-wake distribution. The exception being, in agreement with a previous study (Traut et al., 2023), that the percentage of REMS is significantly decreased during the 3-hr (ZT3-6) following C21 administration, and this effect persisted throughout the entire 6-month dosing period (**Fig. 5C3**).

**Figure 5:**
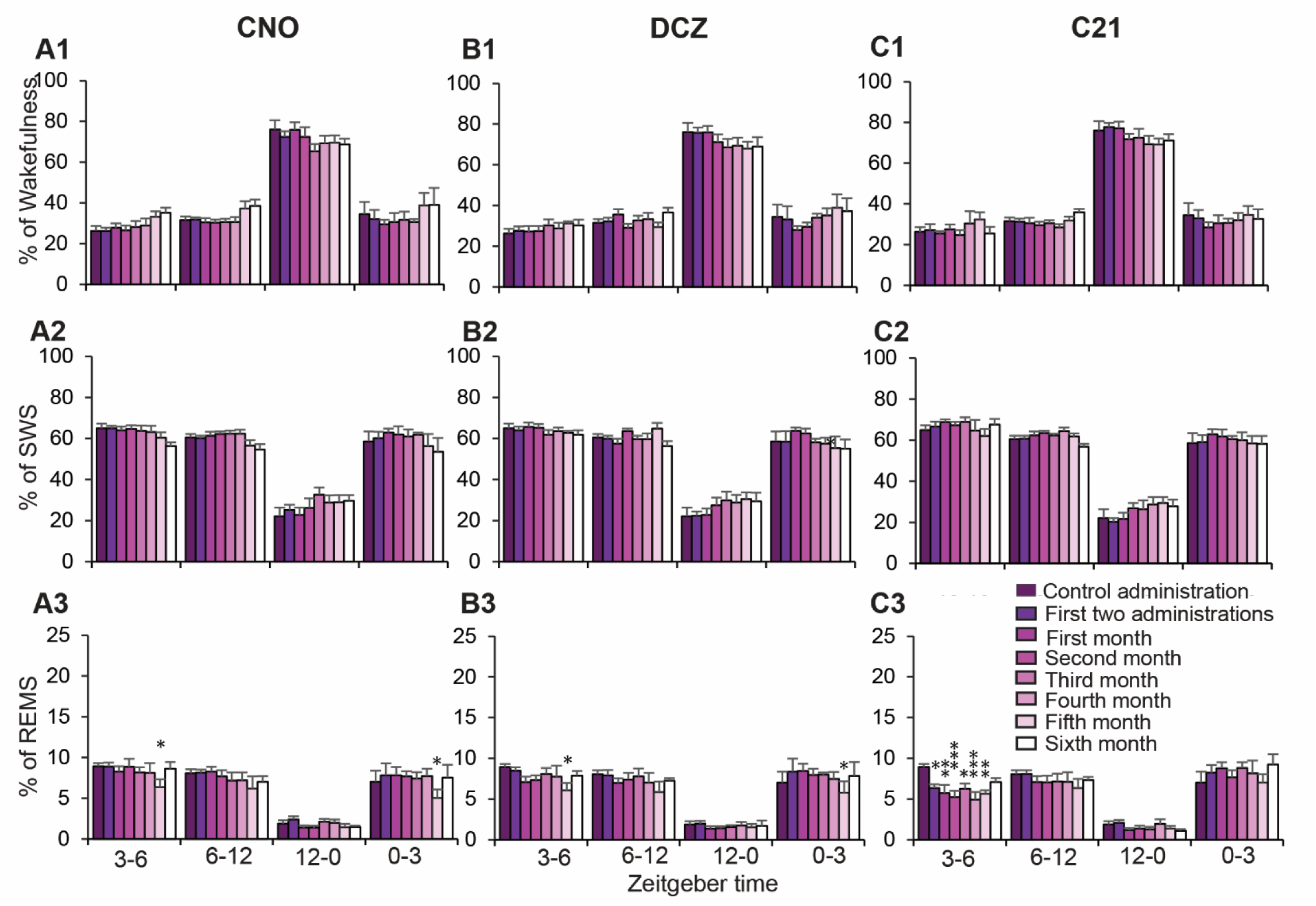
Percentage of sleep-wake states during the 6-month dosing in control mice. (**A-C**) Percentage (± SEM) of wakefulness (**A1,B1,C1**), SWS (**A2,B2,C2**) and REMS (**A3,B3,C3**) following administration of CNO (**A**), DCZ (**B**) or C21 (**C**) and from the two first administrations of each month, between ZT3-6, ZT6-12, ZT12-0 and ZT0-3. BL to 5^th^ month: N = 6 mice; 6^th^ month: N = 5 mice. Significant differences compared with control administration, * p < 0.05, ** p < 0.01, *** p < 0.001, mixed-effect analysis for multiple comparison followed by Bonferroni test.

Interestingly, in the PZ^Vgat-hM3Dq^ mouse group, but not in the control group, the percentage of wakefulness shows an increasing percentage of wakefulness in the 3 hours before drug administration (ZT0-3) over the six months of treatment (CNO not significant but DCZ and C21 at significance of at least P<0.05, **Fig. 4A1, B1, C1**). Since this occurs in SWS enhanced mice but not in control mice (**Fig. 5A1, B1, C1**), immediately prior to jelly presentation, it suggests more an anticipatory behavior to the drug effects on SWS than a potential rewarding effect from the jelly. Supporting this are anecdotal, observed but not quantified, visual observations that PZ^Vgat-hM3Dq^ mice were almost always awake at the time of the drug presentation (ZT3), and that this trend was not seen in the control mouse group which were also anecdotally observed to have been mostly asleep at the time of the drug presentation (ZT3).

### Sleep consolidation

In PZ^Vgat-hM3Dq^ mice, the 6-month treatment significantly affected the percentage of SWS in various SWS bout lengths [CNO: F(20.19, 115.3) = 2.259, P = 0.0037; DCZ: F(23.82, 136.1) = 2.548, P = 0.0004; C21: F(18.92, 108.1) = 2.7633, P = 0.0005] during the 3-hr (ZT3-6) following administration of the agonists. At the same time, the number of SWS episodes in various SWS bout lengths was significantly affected by C21 administration [C21: F(26.20, 149.7) = 1.669, P = 0.0304] but not by CNO or DCZ administrations [CNO: F(20.44, 116.8) = 1.502, P = 0.0918; DCZ: F(14.24, 81.38) = 1.047, P = 0.4171]. As previously reported (Anaclet et al., 2014), the SWS enhancement phenotype is characterized by the lengthening of SWS episodes during the 3-hr period (ZT3-6) following chemogenetic agonist administration. Following control administration, no SWS episodes longer than 20 mins were seen. However, following chemogenetic agonist administration, SWS episodes longer than 20 or even 40 mins are observed (**Fig. 6A1, B1, C1 & Suppl. Table S1-S3**). The proportion of SWS amounts is shifting from medium episode length (2.5-10 mins long) to longer episode duration (10-40 mins long, **Fig. 6A2, B2, C2 & Suppl. Tables S1-S3**). At the same time, the number of wake episodes in various wakefulness bout lengths [CNO: F(12.55, 71.72) = 1.821, P = 0.0580; DCZ: F(13.11, 74.90) = 1.544, P = 0.1212; C21: F(12.36, 70.64) = 1.378, P = 0.1949] and the proportion of wakefulness in various wakefulness bout lengths [CNO: F(24.26, 138.6) = 1.087, P = 0.3656; DCZ: F(27.21, 155.5) = 0.7883, P = 0.7627; C21: F(22.63, 129.3) = 1.096, P = 0.3587] were not affected by the 6-month treatment (**Suppl. Tables S4-S6**). The proportion of REMS in various REMS bout lengths [CNO: F(15.16, 84.45 = 0.8872, P = 0.5813; DCZ: F(13.15, 73.26) = 0.2256, P = 0.9977; C21: F(18.07, 100.7) = 0.5983, P = 0.8937] was not affected by the 6-month treatment, while the number of REMS episodes in various REMS bout lengths was [CNO: F(21.63, 123.6) = 3.235, P < 0.0001; DCZ: F(13.13, 75.02) = 2.118, P = 0.0220; C21: F(19.27, 110.1) = 3.244, P < 0.0001; **Suppl. Tables S7-S9]**. Interestingly, only the number of very short REMS episodes (0.1-0.5 mins long) was significantly decreased during the 3-hr period (ZT3-6) following chemogenetic agonist administration (**Suppl. Table S7-S9**). During the ZT6-12 period following chemogenetic agonist administration, the number of REMS episodes in various REMS bout lengths was still affected by the 6-month treatment [CNO: F(16.43, 93.88) = 2.028, P = 0.0179; DCZ: F(17.57, 100.4) = 1.954, P = 0.0202; C21: F(15.32, 87.57) = 3.669, P < 0.0001; **Suppl. Tables S10-S12]**. However, the proportion of REMS in various bout lengths was not affected during the ZT6-12 period following chemogenetic agonist administration [CNO: F(11.46, 65.46) = 0.9332, P = 0.5173; DCZ: F(10.74, 61.36) = 1.152, P = 0.3394; C21: F(9.673, 55.27) = 1.660, P = 0.1162; **Suppl. Tables S10-S12]**. The decrease in REMS amount (**Fig. 4A3, B3, C3**) was due to a generalized decrease in the number of REMS episodes (**Suppl. Table S10-S12**). During the seven washout days following the 6-month dosing, the number of SWS episodes and the proportion of the SWS in various SWS episode lengths returned to pre-dosing baseline values (**Fig. 6D & Suppl. Table S13**). Similar effects were seen in wakefulness and REMS episode number and proportion (**Suppl. Table S14-S15**).

**Figure 6:**
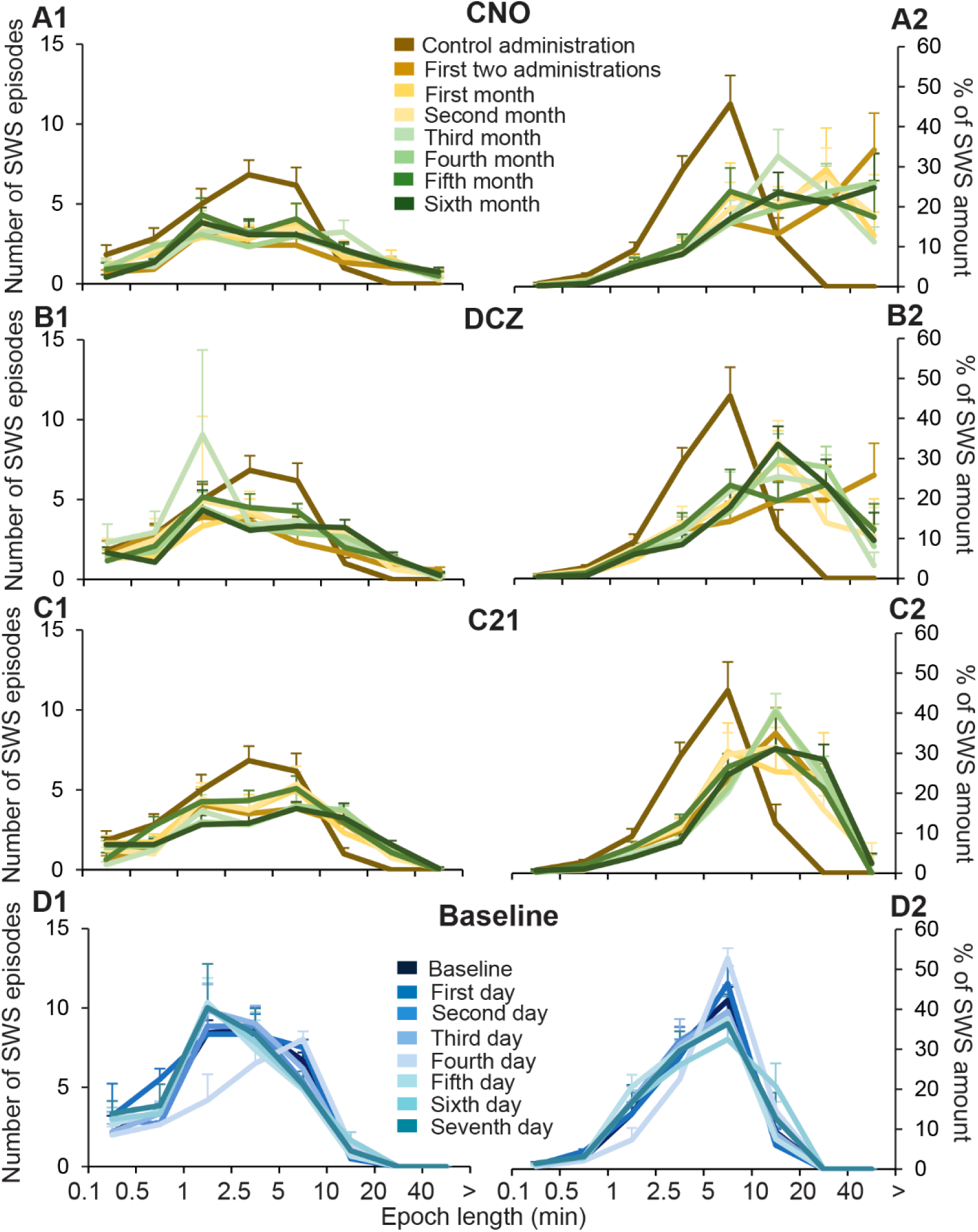
Number of episodes (± SEM) of SWS in various SWS episode lengths and percentage (±SEM) of SWS amount in various SWS episode lengths to the total amount of SWS during the ZT3-6 periods in PZ^Vgat-hM3Dq^ mice following CNO (**A**), DCZ (**B**) and C21 (**C**) administration as compared with control administration. (**D**) Number of episodes (± SEM) of SWS in various SWS episode lengths and percentage (±SEM) of SWS amount in various SWS episode lengths to the total amount of SWS during the first baseline day before the 6-month dosing and during the seven washout days following the 6-month dosing, during the ZT3-6 periods in PZ^Vgat-hM3Dq^ mice. N = 6 mice. For significance, see tables 1-3 & 13.

In the control group, the 6-month treatment significantly affected the percentage of SWS in various SWS bout lengths [CNO: F(49, 237) = 2.258, P =< 0.0001; DCZ: F(49, 237) = 3.000, P < 0.0001; C21: F(49, 237) = 3.636, P < 0.0001] during the 3-hr (ZT3-6) following administration of the agonists. As opposed to the lengthening of the SWS episodes seen in PZ^Vgat-hM3Dq^ mice, in control mice, the proportion of SWS episodes 10-20-min long were decreased following CNO and DCZ administration, every month, as compared with control administration (**Suppl. Table S16-S17**), a phenotype that is seen during aging. The opposite trend was seen following C21 administration (**Suppl. Table S18**), suggesting non-specific action of C21 on SWS maintenance. At the same time, the number of SWS episodes in various SWS bout lengths was significantly affected by CNO and DCZ administrations [CNO: F(49, 237) = 1.611, P = 0.0106; DCZ: F(49, 237) = 2.025, P = 0.0003] but not by C21 administration [C21: F(49, 237) = 1.045, P = 0.4022]. However, individual values remained mostly unaffected (**Suppl. Table S16-S18**). The 6-month treatment did not significantly affect the percentage of wakefulness in various wakefulness bout lengths [CNO: F(49, 237) = 1.301, P = 0.1031; DCZ: F(49, 237) = 0.8693, P = 0.7161; C21: F(49, 312) = 0.8060, P = 0.8191] during the 3-hr (ZT3-6) following the administration of the agonists. The individual values remained mostly unaffected (**Suppl. Table S19-S21**). However, the number of wake episodes in various wake bout lengths was significantly affected by CNO [F(49, 237) = 2.232, P < 0.0001] and C21 [F(49, 237) = 1.516, P = 0.0226] but not by DCZ [F(49, 237) = 1.095, P = 0.3225] 6-month administration. The number of wake episodes in short bout lengths showed some decreases (**Suppl. Table S19-S21**). The percentage of REMS in various bout lengths was not affected by CNO [F(49, 237) = 1.088, P = 0.3331] and DCZ [F(49, 237) = 1.284, P = 0.1144], but by C21 [F(49, 237) = 1.837, P = 0.0015] 6-month administration. The individual values remained mostly unaffected (**Suppl. Table S22-S24**). However, the number of REMS episodes in various bout lengths was affected by CNO [F(49, 237) = 1.987, P = 0.0004], DCZ [F(49, 237) = 1.983, P = 0.0004] and C21 [F(49, 237) = 1.998, P = 0.0003] 6-month administration. Following CNO administration, the number of very short (0.1-0.5-min) REMS episodes was significantly increased and the medium length (0.5-1-min) REMS episode number was decreased as compared with control administration (**Suppl. Table S25**). Following DCZ and C21 treatment, the number of REMS episodes in medium bout length (0.5-1-min) and long bout length (1-2.5-min) were decreased as compared with control administration (**Suppl. Table S26-27**).

### Power spectral distribution

In the PZ^Vgat-hM3Dq^ mouse group, the increase in SWS amount is associated with a significant increase in SWA, and the hallmark of PZ-mediated SWS enhancement is a particularly prominent increase in SWA beyond what is seen during normal NREMS (Anaclet et al., 2014). The 6-month daily drug dosing significantly effected SWS power band distribution during the hour following each chemogenetic agonist administration: CNO [F(28, 140) = 5.535, P < 0.0001, **Fig. 7A**], DCZ [F(28, 140) = 5.271, P < 0.0001, **Fig. 7B**] and C21 [F(28, 140) = 9.393, P < 0.0001, **Fig. 7C**]. Multiple comparisons showed that the proportion of delta (0.5-4.5 Hz) frequency band was significantly increased at the same level throughout the 6-month dosing period (**Fig. 7A-C**). This increase was associated with a significant decrease in the proportion of the theta (4.5-10 Hz) frequency band. Other frequency bands also appear to trend toward a decrease but do not reach significance, except in the case of the last 4 months of C21 treatment, when the proportion of the sigma (10-15 Hz) frequency band is significantly decreased. Interestingly, during the 7-day washout period following the end of the daily drug administration, the proportion of the delta frequency band during SWS remains significantly increased as compared with the 1^st^ baseline day preceding the 6-month treatment (same circadian period: ZT3-6; **Fig. 7D**). Therefore, we hypothesized that SWA could be enhanced during the entire 24-hr sleep-wake cycle. To test this hypothesis, power spectral analysis was performed on day 7 of the washout period and compared with the baseline day preceding the 6-month treatment period (**Fig. 8**). During SWS, the proportion of the delta band was significantly increased during most of the light period (ZT0-9; **Fig. 8B1**). At the same time and beyond, the proportion of the theta band was significantly decreased (ZT0-15; **Fig. 8B2**). The proportion of the sigma band was significantly decreased only during the first circadian period (ZT0-3; **Fig. 8B3**). The proportion of the beta and gamma frequency bands were not affected (**Fig. 8B4-5**). During wakefulness, the proportion of the frequency bands were not significantly affected (**Fig. 8A1-5**). During REMS, the proportion of the delta band was significantly increased during the first circadian period (ZT0-3; **Fig. 8C1**), when the other frequency bands do not show any significant changes (**Fig. 8C2-5**). These results indicate an increased SWS depth/quality during the time of high sleep pressure (ZT0-9) that lasts for at least a week after the end of the chronic SWA enhancement. In contrast to this phenotype, the 24-hr distribution of the three vigilance stages as well as the number and episode duration returned to pre-treatment baseline level on the first day of the 7-day washout period (**Suppl. Fig. S1, Suppl. Table S13-S15**).

**Figure 7:**
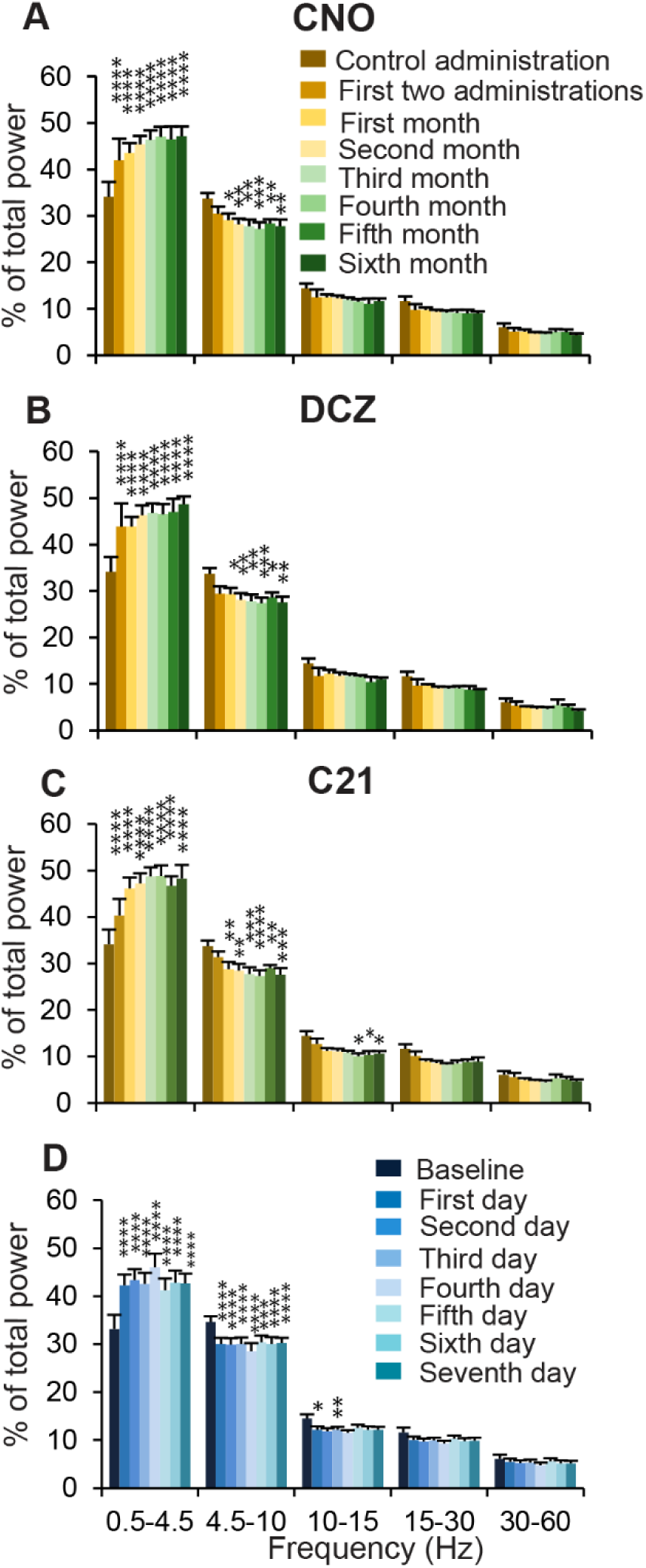
Cortical power bands in PZ^Vgat-hM3Dq^ mice. Percentage (± SEM) of total (0.5-60 Hz) power from the first hour of SWS following administration of CNO (**A**), DCZ (**B**) or C21 (**C**) and from the two first administrations of each month. N = 6 mice. Significant difference as compared with control administration, * p < 0.05, ** p < 0.01, *** p < 0.001, **** p < 0.0001, two-way ANOVA followed by Bonferroni test. (**D**) Percentage (± SEM) of total (0.5-120 Hz) power during SWS from ZT3-6, from the first baseline recording before the 6-month administration period and from the eight days following the end of the 6-month administration period. Significant difference as compared with the first baseline recording, * p < 0.05, ** p < 0.01, *** p < 0.001, **** p < 0.0001, two-way ANOVA followed by Bonferroni test.

**Figure 8:**
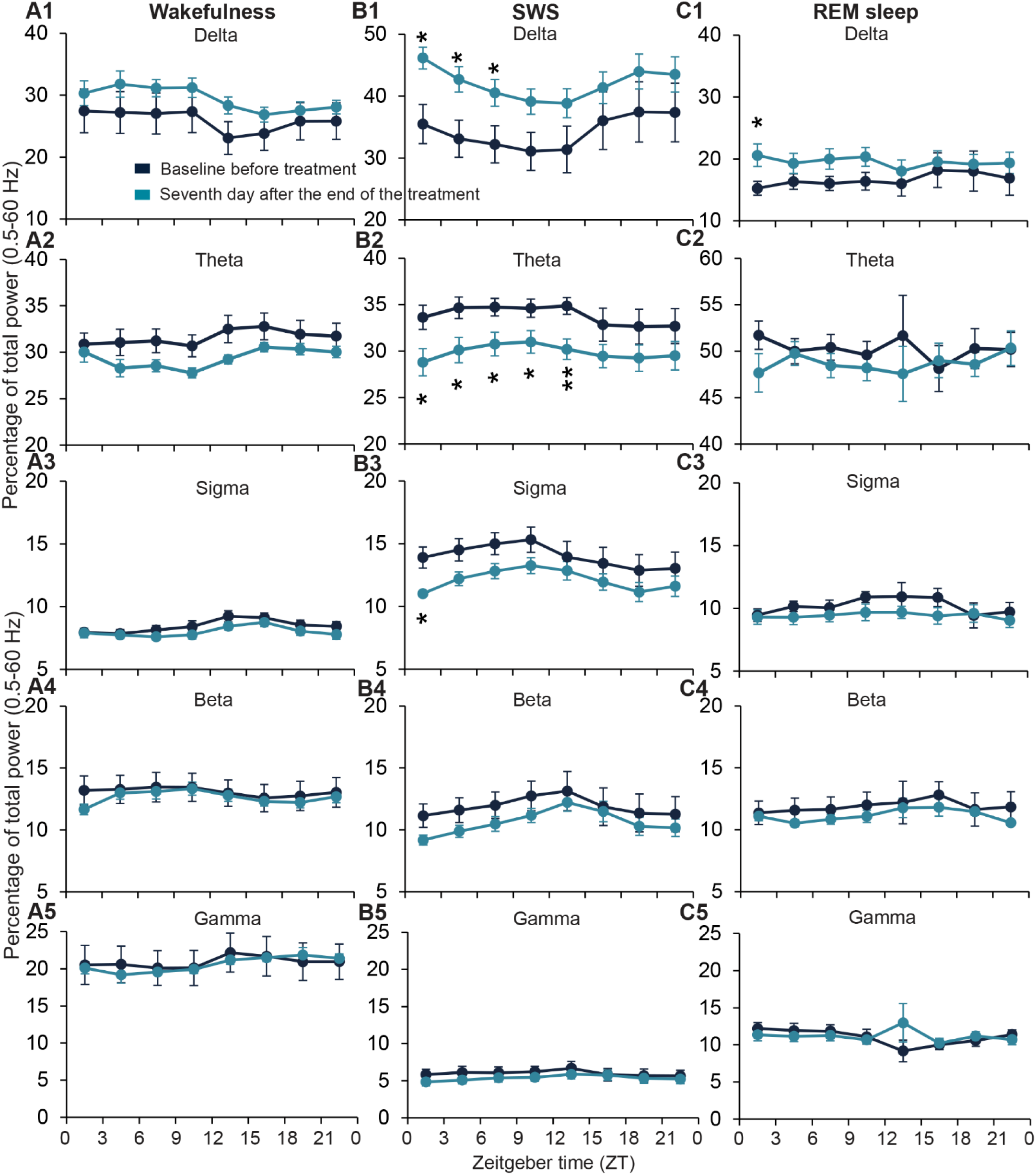
24-hr distribution of the cortical EEG power bands in PZ^Vgat-hM3Dq^ mice. Frequency bands: delta (0.5-4.5, A1-C1), theta (4.5-10 Hz, A2-C2), sigma (10-15 Hz, A3-C3), beta (15-30 Hz, A4-C4) and gamma (30-60 Hz, A5-C5), expressed in percentage (± SEM) of total (0.5-60 Hz) power during wakefulness (A), SWS (B) and REMS (C) across eight circadian periods from the first baseline recording before the 6-month administration period and from the seventh day following the end of the 6-month administration period. N = 6 mice. Significant difference as compared with the first baseline recording, * p < 0.05, ** p < 0.01, mixed-effect analysis for multiple comparison followed by two-stage linear step-up procedure of Benjamini, Krieger and Yekutieli.

In the control group, the 6-month daily drug dosing had no significant effect on SWS power spectral distribution during the 3-hr (ZT3-6) following each chemogenetic agonist administration: CNO [F(2.388, 11.09) = 3.780, P = 0.0503, **Fig. 9A**], DCZ [F(10.11, 68.58) = 1.207, P =0.3018, **Fig. 9B**] and C21 [F(1.302, 6.047) = 3.478, P = 0.1072, **Fig. 9C**]. The proportion of each frequency bands was not affected by the administration of CNO (**Fig. 9A**), DCZ (**Fig. 9B**) or C21 (**Fig. 9C**).

**Figure 9:**
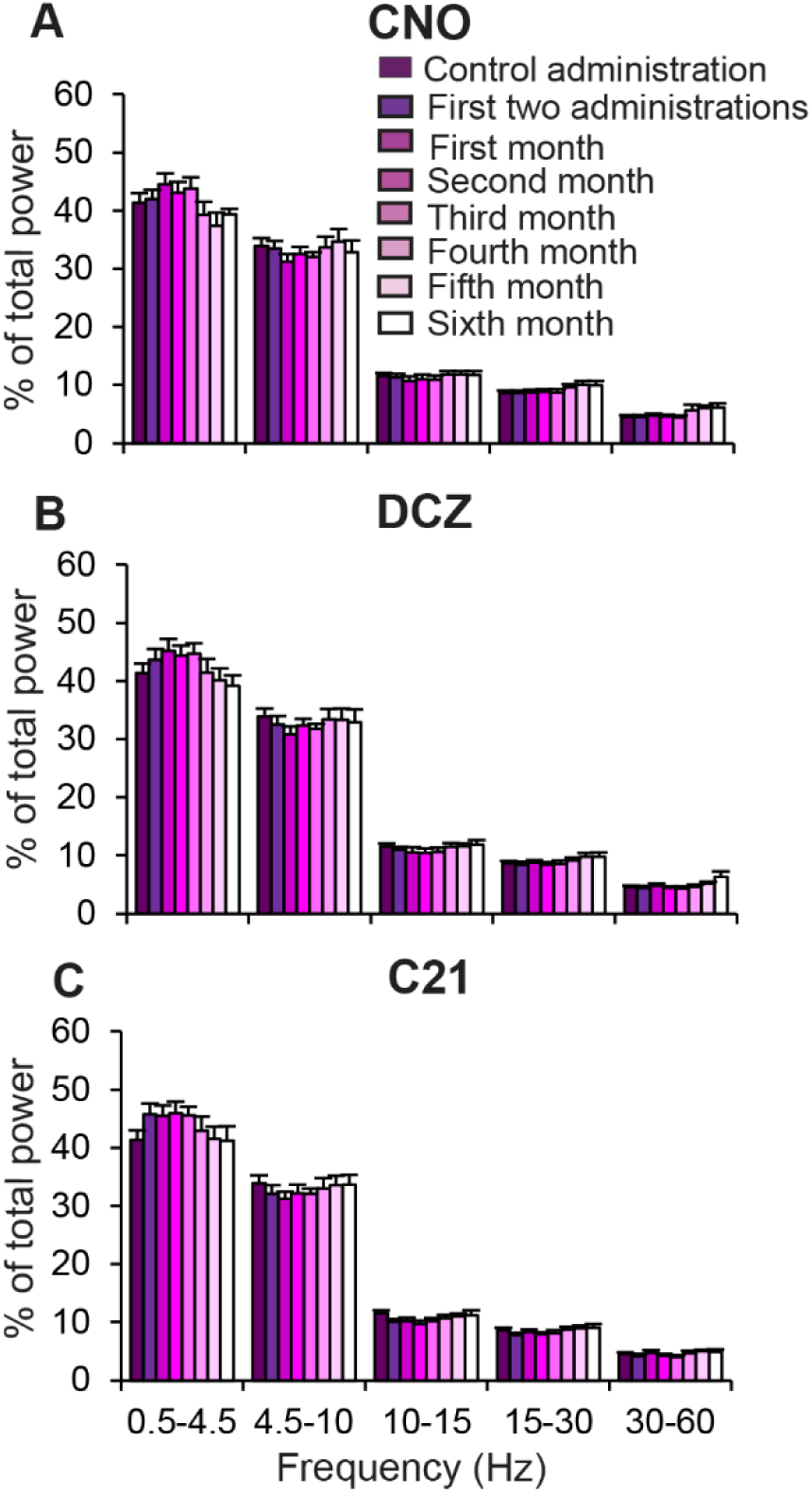
Cortical power bands in control mice. Percentage (± SEM) of total (0.5-60 Hz) power from the first hour of SWS following administration of CNO (**A**), DCZ (**B**) or C21 (**C**) and from the two first administrations of each month. BL to 5^th^ month: N = 6 mice; 6^th^ month: N = 4 mice. Significant difference as compared with control administration, * p < 0.05, ** p < 0.01, mixed-effect analysis for multiple comparison followed by Bonferroni test.

## Discussion

This study validates a method of chronic SWS enhancement in mice. Our results show that daily administration of chemogenetic ligands in mice expressing the chemogenetic receptor hM3Dq in PZ^GABA^ increases SWS amount and enhances SWA at a similar magnitude throughout a 6-month dosing period. These results validate an approach to specifically promoting a sleep stage over long durations of murine lifespans and are therefore critical to supporting further research into the effects of long-term sleep promotion.

### Technical considerations

Chronic administration of drugs associated with sleep-wake recording are accompanied with challenges, such as the stress of daily administration, the wake-promoting effect of the weekly cage change and the constraint of the recording cable. To minimize these effects, we used the voluntary oral administration technique (Jelly) (Ferrari et al., 2022) to administer the chemogenetic ligands. This is particularly important given the potential morbidity (skin breakdown, infection, adhesions, perforated viscus) that can occur with multiple intraperitoneal injections. Daily, the jelly was placed on the floor of the cage, therefore the mouse was not touched by the experimenter, reducing the stress associated with the administration. The weekly cage changes were performed between ZT11-12, when the mice usually show more wakefulness in anticipation of the upcoming dark period. Finally, the mice were connected to the cable and recorded for 2 weeks, followed by 2 weeks free from the cable. Cable connection/disconnection, were also performed between ZT11-12 to minimize the sleep deprivation due to the wake-promoting effect of the mouse handling.

### Limited non-specific effects

We showed that 6-month daily administration of the chemogenetic ligands did not affect the amount of the three vigilance stages, the sleep latencies or the power spectral distribution during SWS in control mice, indicating that the SWS enhancement phenotype in PZ^Vgat-hM3Dq^ mice is the result of specific activation of PZ^GABA^. We purposely used three previously validated chemogenetic agonists: CNO, DCZ and C21 (Ferrari et al., 2022), given sequentially, to limit potential build-up of metabolites and non-specific actions. The known metabolite of CNO is clozapine which binds multiple endogenous receptors (Morrison et al., 2025), including the muscarinic receptor from which hM3Dq is derived. Clozapine is further metabolized into N-desmethylclozapine (NDMC), which also binds numerous endogenous receptors (Raper et al., 2017). Our previous studies showed that acute administration of CNO did not have non-specific effects on the sleep-wake cycle in control mice that do not express hM3Dq in PZ^GABA^ (Anaclet et al., 2014). However, we cannot exclude non-specific effects on other physiological functions. Specifically, higher doses of CNO (1-10 mg/kg) induce non-specific sleep-wake effects in mice acutely (Traut et al., 2023). DCZ metabolites are DCZ N-oxide and C21 which seem to be pharmacologically inert on endogenous receptors (Nagai et al., 2020). Finally, the metabolites of C21 are unknown (Thompson et al., 2018), however, C21 could directly affect nigral neuronal activity (Goutaudier et al., 2020). According to this study, three-week chronic administration of CNO or C21 (1 mg/kg) in wild-type mice showed a significant but limited beneficial effect on anxiety and increased exploration in a novel environment (Tran et al., 2020). However, this effect was abolished after a 14-week treatment. These results are consistent with the antipsychotic action of clozapine (Baldessarini & Frankenburg, 1991) and suggest that C21 displays off target effects even at a lower dose. Specifically, according to a previous report (Traut et al., 2023), we found that REMS amounts were significantly decreased during the 3 hours following administration of C21 in control mice, due to the decrease in the number of medium (1.5-1-min) and long (1.-2.5-min) episodes. These findings confirm that C21 has off-target effects and its use should be carefully controlled. In our experiments, chemogenetic activation of PZ^GABA^ results in a significant reduction of REMS amount lasting about 8-9 hrs (Anaclet et al., 2014; Ferrari et al., 2022), while the SWS enhancement lasts only 3-hr on average. In the present study, REMS amounts are also reduced during the 9-hr period following the administration of the three chemogenetic ligands in PZ^Vgat-hM3Dq^ mice, throughout the 6-month dosing period. Therefore, in PZ^Vgat-hM3Dq^ mice, the REMS inhibitory effect of C21 is masked by the REMS inhibitory effect of PZ^GABA^ activation and does not seem to add an additional non-specific effect.

### Chronic SWS enhancement model

Administrations were performed at ZT3, a time when the mice are usually mostly asleep. Nevertheless, the time in SWS was significantly increased during the 3-hr post administration as compared with control administration. Importantly, SWA was significantly enhanced, indicating a deeper stage of sleep. Therefore, SWS can be promoted chronically during the habitual sleeping period. As previously shown in the acute SWS enhancement model (Anaclet et al., 2014), chronic administrations did not significantly increase the total 24-hr SWS amounts. Similar results are obtained when PZ^GABA^ are activated during the dark period, a time of high waking drive, when the SWS enhancement period is followed by a significant wake rebound (Anaclet et al., 2014), indicating that the sleep needs were fulfilled and suggesting a potential homeostatic regulation of wakefulness. Even though the daily amount of SWS is unchanged when SWS is enhanced during the normal sleep period, the mice experience deeper, more consolidated sleep, similar to a sleep rebound following sleep deprivation. This extended and intensified restorative activity may lead to therapeutic benefits (Greene & Frank, 2010; Lee et al., 2020). Interestingly, it was observed that when the experimenter opened the sleep chamber, daily at ZT3, all the PZ^Vgat-hM3Dq^ mice were awake and waiting for the Jelly delivery, while the control mice were mostly asleep. In this particular laboratory environment, the sleep chamber is located behind a door from where the jelly is prepared in a small positive air-pressure room, and is itself additionally light- and sound-proof, with a separate ventilation system. Cues to the administration time are thus possible, but unlikely. This anticipatory behavior is reflected by the increase in wake time during the ZT0-3 period in PZ^Vgat-hM3Dq^ mice (**Fig. 3A1, B1, C1**) but not in control mice (**Fig. 5A1, B1, C1**). These visual observations are reinforced by the short latency to consume the jelly. In fact, PZ^Vgat-hM3Dq^ mice immediately approached the Jelly and began eating it within a few seconds after the jelly was placed in the cage while the control mice seemed to be less excited about the jelly, thereby making it unlikely that this was food anticipatory activity (Fuller et al., 2008). It is quite likely that PZ^Vgat-hM3Dq^ mice exhibit reward-seeking anticipatory behavior and that therefore the deep and consolidated sleep following jelly administration is itself hedonic, particularly since the jelly, which is sweet and contains metabolites of the atypical antipsychotic clozapine, did not elicit such behaviors in control animals. The hedonic aspects of sleeping has been discussed previously (Rial et al., 2018), however, this is to our knowledge the first indication of sleep being a primary reinforcer. The relationship between the reward system and the sleep-wake system is well known. As highlighted in the Rial et al. (2018) review, SWA seems to be of particular importance. However, the mechanism by which sleep activates the reward system remains poorly understood due to the lack of tools permitting the study of the role of sleep using gain-of-sleep experiments. Our mouse model of SWS enhancement is ideal to study the mechanisms underlying the rewarding role of deep sleep and high SWA (see below).

The increase in SWS amount is associated with an increase in the proportion of SWA, as previously shown (Anaclet et al., 2014). The increase in SWA is associated with a decrease of the proportion of theta rhythm, indicating a deeper stage of SWS. This phenotype remains of the same magnitude throughout the 6-month dosing period, indicating no compensatory mechanisms, receptor desensitization or neuronal circuitry fatigue. In fact, SWA tends to increase over the 6-month dosing period, as does the amount and duration of PZ-mediated SWS. Interestingly, SWA remains significantly increased even after the end of the 6-month treatment period when the mice are no longer receiving chemogenetic agonist-infused jelly. This may indicate a plastic effect on the neuronal circuitry regulating SWA during SWS. We hypothesize that high amplitude SWA prompts the circuitry involved in regulating SWA and as a result, deep consolidated sleep will promote further deep consolidated sleep while poor sleep quality will do the opposite. There might be similarities to human sleep hygiene and cognitive behavioral therapy coaching for patients with insomnia, whereby small initial changes build upon each other and result in long-lasting beneficial sleep pattern changes. The mechanisms underlying these observations could also be studied using this model of chronic SWS enhancement.

SWS enhancement is accompanied by a reduction of REMS time. Therefore, our mouse model could open the door to specifically studying the role of SWS in various phenomena including memory, neurodegeneration, and other neurologic diseases. The reduction of REMS following the administration lasts longer than the SWS enhancement period and is not compensated by a rebound, indicating that the reduction in REMS did not induce a dramatic lack requiring homeostatic rebound. In fact, following sleep deprivation, the REMS rebound never fully compensates for the lack of REMS (Achermann & Borbely, 2011), indicating that REMS duration is not strictly regulated. Whereas REMS is necessary for various neurological functions (Mukai & Yamanaka, 2023), excessive REMS is associated with mood disorders and depression (Yasugaki et al., 2025). REMS deprivation elicits positive effects on mood and is used as a treatment in patients suffering from severe depression (Hemmeter et al., 2010). We also observed a trend to decreased REMS daily amounts during the course of the 6-month period, suggesting a negative entrainment of the neuronal circuitry regulating REMS. Even if the daily amount of REMS seems to return to normal at the end of the 6-month administration period, it is possible that chronic REMS inhibition entrains the circuitry leading to a reduced daily amount. On the contrary, increased amounts of REMS would be expected to promote the REM-on neuronal circuitry. Because REMS amounts are increased in patients suffering from depression (Yasugaki et al., 2025), and REMS deprivation positively impacts the mood (Hemmeter et al., 2010), this mechanism might be involved in mood regulation. This hypothesis could be studied using this mouse model of chronic SWS enhancement.

In conclusion, we showed that our mouse model permits long-term SWS enhancement and could be an appropriate tool to study the beneficial role of SWS in physiological functions, such as cognition and reward, as well as in chronic conditions such as aging, Alzheimer’s disease and epilepsy.

## Supplementary Figure and Table

**Figure S1:**
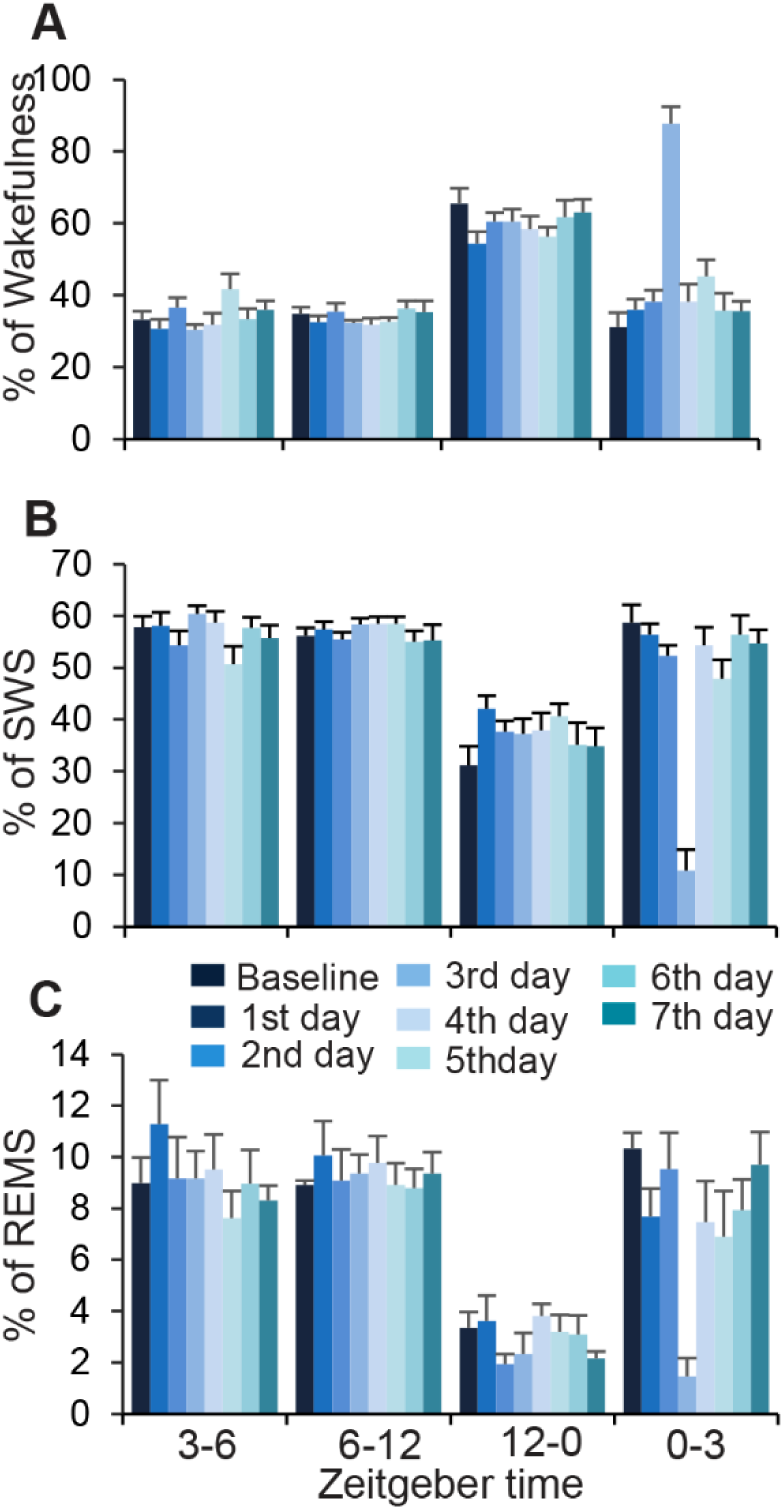
Percentage of sleep-wake states during the washout period following the 6-month dosing in PZ^Vgat-hM3Dq^ mice. (**A-C**) Percentage (± SEM) of wakefulness (**A**), SWS (**B**) and REMS (**C**) during the 7-day following discontinuation of the chemogenetic agonist dosing, between ZT3-6, ZT6-12, ZT12-0 and ZT0-3. N = 6 mice. No significant differences with the pre-dosing baseline period, two-way ANOVA followed by Bonferroni test.

**Supplementary Table S1:** Percentage (±SEM) of **SWS** amount in various SWS episode lengths to the total amount of SWS and number of episodes (± SEM) of SWS in various SWS episode lengths during the **ZT3-6** periods in PZ^Vgat-hM3Dq^ mice following **CNO** administration as compared with control administration. N = 6 mice. α: p < 0.05, β: p < 0.01, two-way ANOVA followed by a post hoc Bonferroni test.

| Percentage (±SEM) of SWS amount in various SWS episode lengths to the total amount of SWS during the ZT3-6 period following CNO administration as compared with control administration |  |  |  |  |  |  |  |  |  |  |  |  |  |  |  |  |  |  |  |  |  |  |  |  |
| --- | --- | --- | --- | --- | --- | --- | --- | --- | --- | --- | --- | --- | --- | --- | --- | --- | --- | --- | --- | --- | --- | --- | --- | --- |
| Episode length (min) | Control |  |  | 1<sup>st</sup> 2 administrations |  |  | 1<sup>st</sup> month |  |  | 2<sup>nd</sup> month |  |  | 3<sup>rd</sup> month |  |  | 4<sup>th</sup> month |  |  | 5<sup>th</sup> month |  |  | 6<sup>th</sup> month |  |  |
|  | Avg | ± | SEM | Avg | ± | SEM | Avg | ± | SEM | Avg | ± | SEM | Avg | ± | SEM | Avg | ± | SEM | Avg | ± | SEM | Avg | ± | SEM |
| 0.1-0.5 | 0.8 | ± | 0.2 | 0.3 | ± | 0.1 | 0.5 | ± | 0.2 | 0.3 | ± | 0.1 | 0.4 | ± | 0.2 | 0.3 | ± | 0.1 | 0.2 | ± | 0.1 | 0.1 | ± | 0.1 |
| 0.5-1 | 2.8 | ± | 0.6 | 0.9 | ± | 0.5 | 1.5 | ± | 0.6 | 1.5 | ± | 0.4 | 0.9 | ± | 0.5 | 1.4 | ± | 0.3 | 0.8 | ± | 0.2 | 0.8 | ± | 0.1 |
| 1-2.5 | 9.2 | ± | 1.9 | 5.7 | ± | 2.4 | 4.8 | ± | 1.9 | 4.8 | ± | 1.9 | 4.9 | ± | 2.6 | 5.1 | ± | 2.4 | 5.9 | ± | 1.3 | 5.0 | ± | 1.2 |
| 2.5-5 | 29.2 | ± | 3.5 | 9.2 | ± | 4.0 α | 9.1 | ± | 3.0 α | 8.2 | ± | 3.2 α | 9.9 | ± | 2.5 α | 8.0 | ± | 4.4 α | 10.1 | ± | 3.0 α | 8.1 | ± | 2.4 β |
| 5-10 | 45.6 | ± | 7.1 | 15.9 | ± | 3.4 α | 22.9 | ± | 8.1 | 19.6 | ± | 6.4 | 16.8 | ± | 4.8 | 15.9 | ± | 3.2 α | 23.8 | ± | 5.9 | 17.0 | ± | 2.5 |
| 10-20 | 12.5 | ± | 4.7 | 13.2 | ± | 6.9 | 19.1 | ± | 2.6 | 20.5 | ± | 4.5 | 32.5 | ± | 6.8 | 19.7 | ± | 3.9 | 19.8 | ± | 4.3 | 23.4 | ± | 5.2 |
| 20-40 | 0.0 | ± | 0.0 | 20.6 | ± | 6.5 | 29.4 | ± | 10.3 | 27.4 | ± | 7.3 | 23.4 | ± | 6.7 | 23.7 | ± | 7.1 | 22.0 | ± | 5.3 | 21.0 | ± | 3.0 β |
| 40-∞ | 0.0 | ± | 0.0 | 34.2 | ± | 9.2 | 12.7 | ± | 5.9 | 17.7 | ± | 10.3 | 11.2 | ± | 3.7 | 25.9 | ± | 7.5 | 17.4 | ± | 9.1 | 24.7 | ± | 8.6 |
| Number of episodes (± SEM) of SWS in various SWS episode lengths during the ZT3-6 period following CNO administration as compared with control administration |  |  |  |  |  |  |  |  |  |  |  |  |  |  |  |  |  |  |  |  |  |  |  |  |
| Episode length (min) | Control |  |  | 1<sup>st</sup> 2 administrations |  |  | 1<sup>st</sup> month |  |  | 2<sup>nd</sup> month |  |  | 3<sup>rd</sup> month |  |  | 4<sup>th</sup> month |  |  | 5<sup>th</sup> month |  |  | 6<sup>th</sup> month |  |  |
|  | Avg | ± | SEM | Avg | ± | SEM | Avg | ± | SEM | Avg | ± | SEM | Avg | ± | SEM | Avg | ± | SEM | Avg | ± | SEM | Avg | ± | SEM |
| 0.1-0.5 | 1.8 | ± | 0.6 | 0.8 | ± | 0.1 | 1.2 | ± | 0.4 | 0.9 | ± | 0.4 | 1.2 | ± | 0.5 | 1.0 | ± | 0.4 | 0.9 | ± | 0.4 | 0.4 | ± | 0.2 |
| 0.5-1 | 2.8 | ± | 0.7 | 0.9 | ± | 0.4 | 1.9 | ± | 0.6 | 2.1 | ± | 0.5 | 1.1 | ± | 0.6 | 2.3 | ± | 0.5 | 1.3 | ± | 0.2 | 1.3 | ± | 0.2 |
| 1-2.5 | 5.0 | ± | 1.0 | 3.2 | ± | 1.0 | 2.9 | ± | 1.0 | 3.4 | ± | 1.2 | 3.3 | ± | 1.3 | 3.1 | ± | 1.0 | 4.3 | ± | 1.0 | 3.8 | ± | 0.9 |
| 2.5-5 | 6.8 | ± | 0.9 | 2.4 | ± | 1.1 | 2.8 | ± | 0.8 | 3.0 | ± | 1.1 | 3.3 | ± | 0.7 α | 2.3 | ± | 0.9 α | 3.2 | ± | 0.8 | 3.1 | ± | 1.0 |
| 5-10 | 6.2 | ± | 1.1 | 2.4 | ± | 0.4 | 3.8 | ± | 1.2 | 3.5 | ± | 1.0 | 3.0 | ± | 0.6 | 3.0 | ± | 0.4 | 4.1 | ± | 1.0 | 3.1 | ± | 0.3 |
| 10-20 | 1.0 | ± | 0.4 | 1.3 | ± | 0.7 | 1.7 | ± | 0.2 | 2.0 | ± | 0.4 | 3.3 | ± | 0.7 | 2.1 | ± | 0.4 | 2.1 | ± | 0.4 | 2.2 | ± | 0.4 |
| 20-40 | 0.0 | ± | 0.0 | 1.1 | ± | 0.4 | 1.6 | ± | 0.5 | 1.3 | ± | 0.3 | 1.3 | ± | 0.4 | 1.4 | ± | 0.4 | 1.3 | ± | 0.3 α | 1.3 | ± | 0.2 α |
| 40-∞ | 0.0 | ± | 0.0 | 0.7 | ± | 0.2 | 0.3 | ± | 0.1 | 0.5 | ± | 0.3 | 0.3 | ± | 0.1 | 0.7 | ± | 0.2 | 0.5 | ± | 0.3 | 0.8 | ± | 0.3 |

**Supplementary Table S2:**
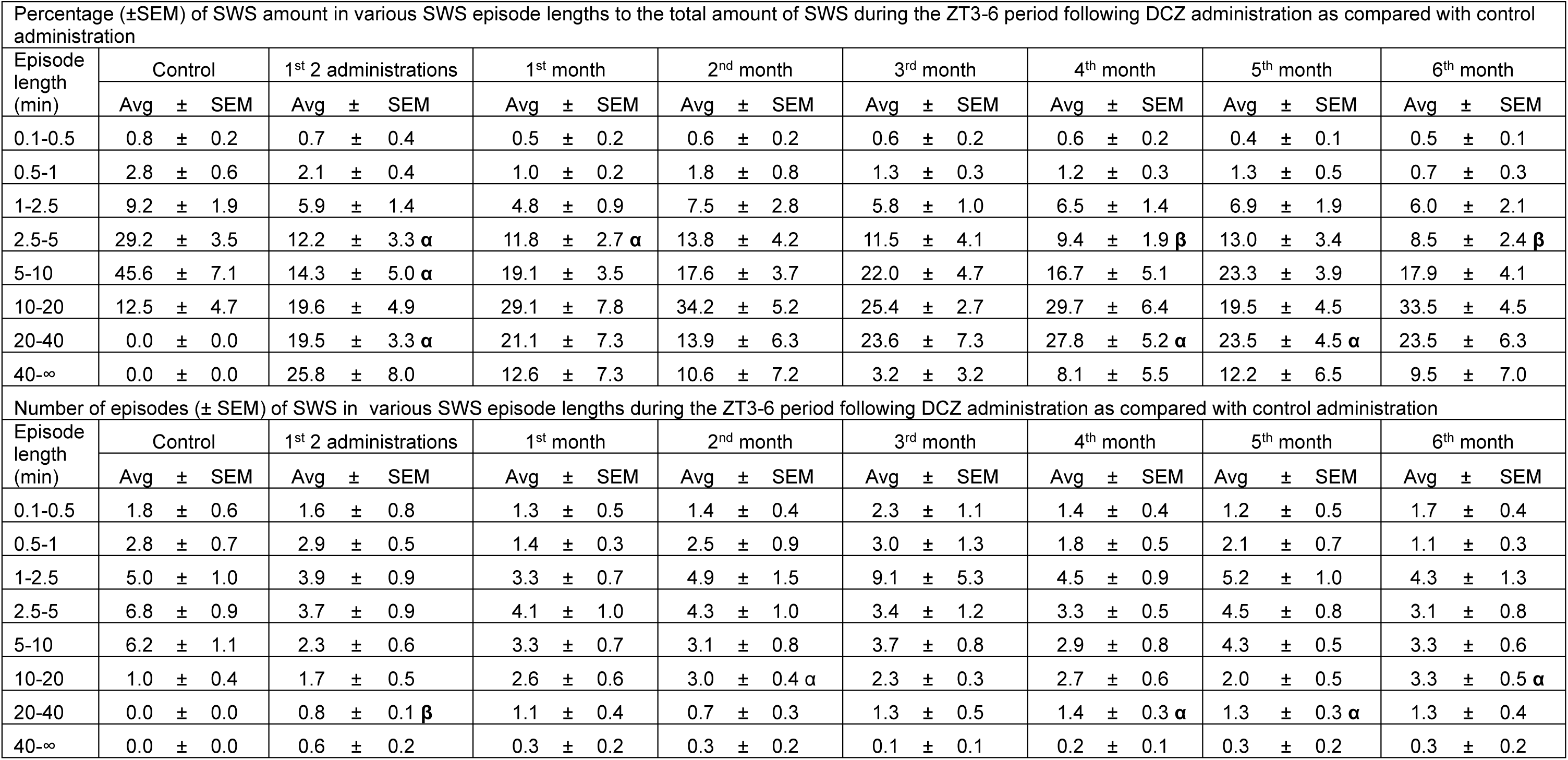
Percentage (±SEM) of **SWS** amount in various SWS episode lengths to the total amount of SWS and number of episodes (± SEM) of SWS in various SWS episode lengths during the **ZT3-6** periods in PZ^Vgat-hM3Dq^ mice following **DCZ** administration as compared with control administration. N = 6 mice. α: p < 0.05, β: p < 0.01, two-way ANOVA followed by a post hoc Bonferroni test.

**Supplementary Table S3:** Percentage (±SEM) of **SWS** amount in various SWS episode lengths to the total amount of SWS and number of episodes (± SEM) of SWS in various SWS episode lengths during the **ZT3-6** periods in PZ^Vgat-hM3Dq^ mice following **C21** administration as compared with control administration. N = 6 mice. α: p < 0.05, β: p < 0.01, two-way ANOVA followed by a post hoc Bonferroni test.

| Percentage (±SEM) of SWS amount in various SWS episode lengths to the total amount of SWS during the ZT3-6 period following C21 administration as compared with control administration | | | | | | | | |
| --- | --- | --- | --- | --- | --- | --- | --- | --- |
| Episode length (min) | Control | 1<sup>st</sup> 2 administrations | 1<sup>st</sup> month | 2<sup>nd</sup> month | 3<sup>rd</sup> month | 4<sup>th</sup> month | 5<sup>th</sup> month | 6<sup>th</sup> month |
| | Avg ± SEM | Avg ± SEM | Avg ± SEM | Avg ± SEM | Avg ± SEM | Avg ± SEM | Avg ± SEM | Avg ± SEM |
| 0.1-0.5 | 0.8 ± 0.2 | 0.2 ± 0.1 | 0.6 ± 0.2 | 0.4 ± 0.2 | 0.2 ± 0.1 | 0.2 ± 0.1 | 0.2 ± 0.2 | 0.7 ± 0.3 |
| 0.5-1 | 2.8 ± 0.6 | 1.3 ± 0.5 | 1.2 ± 0.3 | 0.7 ± 0.3 | 0.9 ± 0.4 | 0.9 ± 0.4 | 2.0 ± 0.5 | 1.0 ± 0.2 |
| 1-2.5 | 9.2 ± 1.9 | 6.0 ± 2.0 | 6.4 ± 1.6 | 6.7 ± 1.3 | 4.9 ± 1.7 | 5.0 ± 1.6 | 6.4 ± 1.4 | 3.9 ± 0.9 |
| 2.5-5 | 29.2 ± 3.5 | 10.5 ± 3.1 $\alpha$ | 11.3 ± 2.7 $\alpha$ | 11.5 ± 2.0 $\alpha$ | 8.4 ± 2.2 $\beta$ | 8.3 ± 2.3 $\beta$ | 12.7 ± 2.0 $\alpha$ | 7.8 ± 2.0 $\beta$ |
| 5-10 | 45.6 ± 7.1 | 23.6 ± 2.6 | 30.3 ± 7.2 | 29.6 ± 5.5 | 19.4 ± 5.8 | 20.2 ± 5.7 | 26.5 ± 3.4 | 24.6 ± 5.2 |
| 10-20 | 12.5 ± 4.7 | 35.0 ± 6.4 | 25.4 ± 8.6 | 31.4 ± 5.0 | 39.2 ± 4.1 $\alpha$ | 41.0 ± 3.9 | 31.1 ± 2.9 | 31.2 ± 8.0 |
| 20-40 | 0.0 ± 0.0 | 20.9 ± 6.5 | 24.8 ± 10.3 | 15.8 ± 3.6 $\alpha$ | 24.7 ± 6.9 | 22.2 ± 5.5 | 21.1 ± 4.2 $\alpha$ | 28.4 ± 3.8 $\beta$ |
| 40-∞ | 0.0 ± 0.0 | 2.4 ± 2.4 | 0.0 ± 0.0 | 3.8 ± 3.8 | 2.3 ± 2.3 | 2.3 ± 2.3 | 0.0 ± 0.0 | 2.5 ± 2.5 |
| Number of episodes (± SEM) of SWS in various SWS episode lengths during the ZT3-6 period following C21 administration as compared with control administration | | | | | | | | |
| Episode length (min) | Control | 1<sup>st</sup> 2 administrations | 1<sup>st</sup> month | 2<sup>nd</sup> month | 3<sup>rd</sup> month | 4<sup>th</sup> month | 5<sup>th</sup> month | 6<sup>th</sup> month |
| | Avg ± SEM | Avg ± SEM | Avg ± SEM | Avg ± SEM | Avg ± SEM | Avg ± SEM | Avg ± SEM | Avg ± SEM |
| 0.1-0.5 | 1.8 ± 0.6 | 0.6 ± 0.3 | 1.7 ± 0.5 | 1.3 ± 0.5 | 0.5 ± 0.2 | 0.3 ± 0.2 | 0.7 ± 0.4 | 1.6 ± 0.4 |
| 0.5-1 | 2.8 ± 0.7 | 1.6 ± 0.5 | 1.8 ± 0.5 | 1.0 ± 0.5 | 1.3 ± 0.3 | 1.3 ± 0.3 | 2.8 ± 0.6 | 1.6 ± 0.4 |
| 1-2.5 | 5.0 ± 1.0 | 4.1 ± 1.1 | 4.3 ± 0.9 | 4.4 ± 0.6 | 3.6 ± 1.1 | 3.7 ± 1.0 | 4.3 ± 0.8 | 2.8 ± 0.6 |
| 2.5-5 | 6.8 ± 0.9 | 3.5 ± 1.0 | 3.8 ± 1.0 | 3.8 ± 0.6 | 2.9 ± 0.7 $\alpha$ | 2.8 ± 0.8 $\alpha$ | 4.3 ± 0.6 | 2.9 ± 0.7 |
| 5-10 | 6.2 ± 1.1 | 3.8 ± 0.4 | 5.0 ± 1.1 | 5.3 ± 0.8 | 3.6 ± 1.2 | 3.8 ± 1.1 | 5.1 ± 0.8 | 3.8 ± 0.4 |
| 10-20 | 1.0 ± 0.4 | 2.9 ± 0.6 | 2.3 ± 0.9 | 3.1 ± 0.6 | 3.5 ± 0.5 $\beta$ | 3.8 ± 0.4 $\beta$ | 3.0 ± 0.3 $\alpha$ | 3.3 ± 0.9 |
| 20-40 | 0.0 ± 0.0 | 1.0 ± 0.3 | 1.1 ± 0.4 | 0.8 ± 0.2 $\alpha$ | 1.4 ± 0.4 $\alpha$ | 1.3 ± 0.3 $\alpha$ | 1.1 ± 0.2 $\alpha$ | 1.6 ± 0.2 $\beta$ |
| 40-∞ | 0.0 ± 0.0 | 0.1 ± 0.1 | 0.0 ± 0.0 | 0.1 ± 0.1 | 0.1 ± 0.1 | 0.1 ± 0.1 | 0.0 ± 0.0 | 0.1 ± 0.1 |

**Supplementary Table S4:**
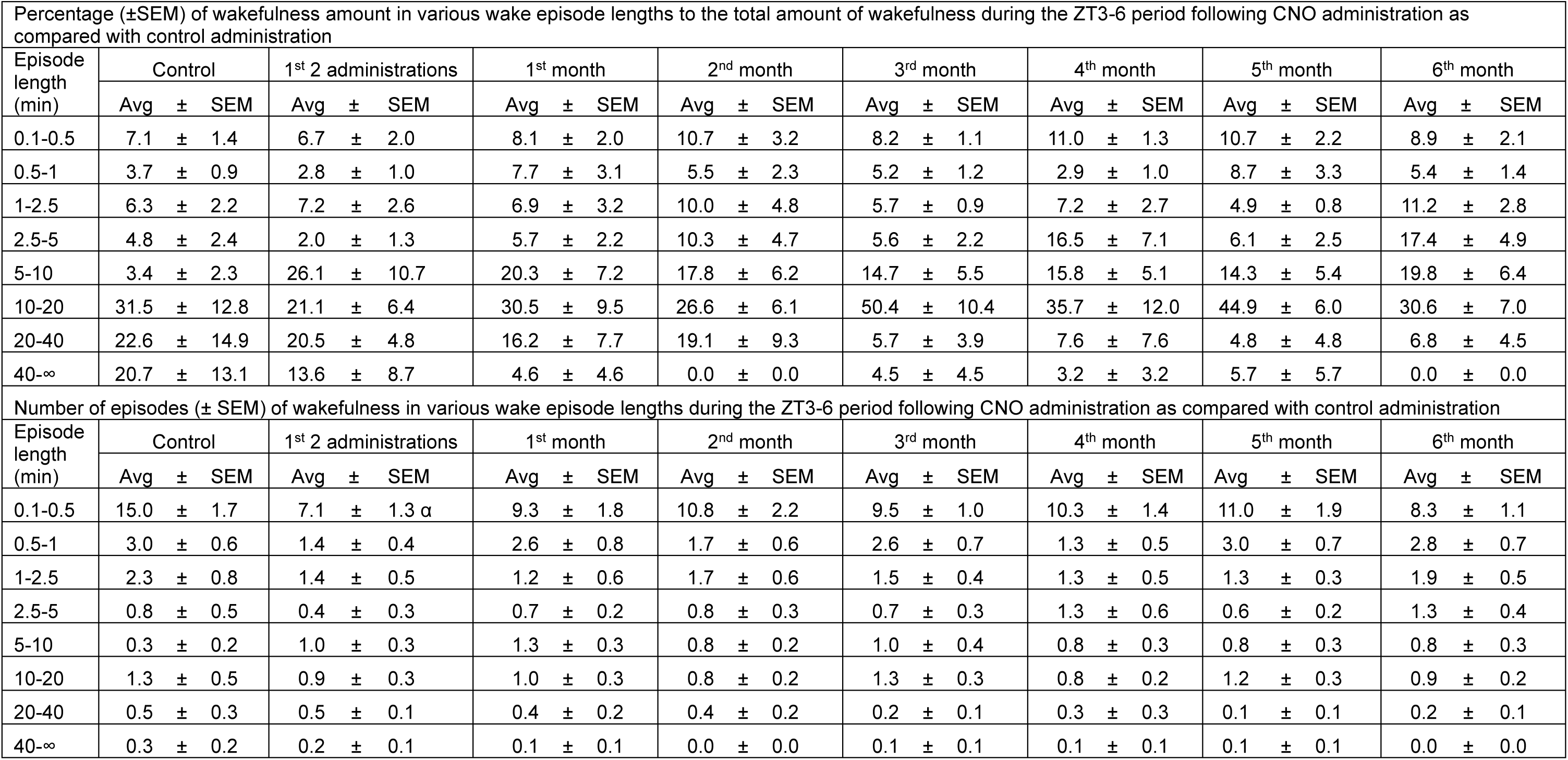
Percentage (±SEM) of **wakefulness** amount in various wake episode lengths to the total amount of wakefulness and number of episodes (± SEM) of wakefulness in various wake episode lengths during the **ZT3-6** periods in PZ^Vgat-hM3Dq^ mice following **CNO** administration as compared with control administration. N = 6 mice. No significance, two-way ANOVA followed by a post hoc Bonferroni test.

**Supplementary Table S5:** Percentage (±SEM) of **wakefulness** amount in various wake episode lengths to the total amount of wakefulness and number of episodes (± SEM) of wakefulness in various wake episode lengths during the **ZT3-6** periods in PZ^Vgat-hM3Dq^ mice following **DCZ** administration as compared with control administration. N = 6 mice. α: p < 0.05, two-way ANOVA followed by a post hoc Bonferroni test.

| Percentage (±SEM) of wakefulness amount in various wake episode lengths to the total amount of wakefulness during the ZT3-6 period following DCZ administration as compared with control administration |  |  |  |  |  |  |  |  |  |  |  |  |  |  |  |  |  |  |  |  |  |  |  |  |
| --- | --- | --- | --- | --- | --- | --- | --- | --- | --- | --- | --- | --- | --- | --- | --- | --- | --- | --- | --- | --- | --- | --- | --- | --- |
| Episode length (min) | Control |  |  | 1<sup>st</sup> 2 administrations |  |  | 1<sup>st</sup> month |  |  | 2<sup>nd</sup> month |  |  | 3<sup>rd</sup> month |  |  | 4<sup>th</sup> month |  |  | 5<sup>th</sup> month |  |  | 6<sup>th</sup> month |  |  |
|  | Avg | ± | SEM | Avg | ± | SEM | Avg | ± | SEM | Avg | ± | SEM | Avg | ± | SEM | Avg | ± | SEM | Avg | ± | SEM | Avg | ± | SEM |
| 0.1-0.5 | 7.1 | ± | 1.4 | 10.7 | ± | 3.7 | 7.2 | ± | 1.3 | 7.0 | ± | 1.5 | 9.9 | ± | 4.1 | 9.8 | ± | 1.5 | 13.4 | ± | 4.7 | 10.7 | ± | 3.1 |
| 0.5-1 | 3.7 | ± | 0.9 | 6.6 | ± | 1.7 | 2.8 | ± | 0.8 | 6.2 | ± | 1.4 | 3.4 | ± | 1.1 | 6.2 | ± | 1.5 | 5.9 | ± | 1.0 | 5.0 | ± | 0.8 |
| 1-2.5 | 6.3 | ± | 2.2 | 15.2 | ± | 4.9 | 7.5 | ± | 2.7 | 11.9 | ± | 3.6 | 6.6 | ± | 2.6 | 6.4 | ± | 1.6 | 13.6 | ± | 2.4 | 10.7 | ± | 2.1 |
| 2.5-5 | 4.8 | ± | 2.4 | 18.9 | ± | 4.1 | 8.9 | ± | 4.5 | 11.9 | ± | 7.2 | 14.8 | ± | 6.3 | 10.9 | ± | 5.1 | 14.9 | ± | 4.5 | 20.2 | ± | 6.2 |
| 5-10 | 3.4 | ± | 2.3 | 12.5 | ± | 4.5 | 32.2 | ± | 10.7 | 24.9 | ± | 5.0 α | 12.6 | ± | 3.0 | 25.1 | ± | 5.4 | 9.8 | ± | 4.5 | 22.0 | ± | 8.0 |
| 10-20 | 31.5 | ± | 12.8 | 15.8 | ± | 6.1 | 26.8 | ± | 7.5 | 17.0 | ± | 6.2 | 23.7 | ± | 10.9 | 23.1 | ± | 8.7 | 22.0 | ± | 9.1 | 17.9 | ± | 6.1 |
| 20-40 | 22.6 | ± | 14.9 | 14.2 | ± | 7.6 | 14.6 | ± | 9.5 | 11.1 | ± | 5.8 | 9.1 | ± | 6.0 | 14.5 | ± | 7.2 | 13.2 | ± | 6.5 | 13.3 | ± | 10.3 |
| 40-∞ | 20.7 | ± | 13.1 | 6.0 | ± | 6.0 | 0.0 | ± | 0.0 | 10.1 | ± | 10.1 | 13.4 | ± | 8.5 | 4.0 | ± | 4.0 | 7.0 | ± | 7.0 | 0.0 | ± | 0.0 |
| Number of episodes (± SEM) of wakefulness in various wake episode lengths during the ZT3-6 period following DCZ administration as compared with control administration |  |  |  |  |  |  |  |  |  |  |  |  |  |  |  |  |  |  |  |  |  |  |  |  |
| Episode length (min) | Control |  |  | 1<sup>st</sup> 2 administrations |  |  | 1<sup>st</sup> month |  |  | 2<sup>nd</sup> month |  |  | 3<sup>rd</sup> month |  |  | 4<sup>th</sup> month |  |  | 5<sup>th</sup> month |  |  | 6<sup>th</sup> month |  |  |
|  | Avg | ± | SEM | Avg | ± | SEM | Avg | ± | SEM | Avg | ± | SEM | Avg | ± | SEM | Avg | ± | SEM | Avg | ± | SEM | Avg | ± | SEM |
| 0.1-0.5 | 15.0 | ± | 1.7 | 8.6 | ± | 1.4 | 10.2 | ± | 1.7 | 10.5 | ± | 2.1 | 11.0 | ± | 1.5 | 10.6 | ± | 1.0 | 12.3 | ± | 1.4 | 10.0 | ± | 0.9 |
| 0.5-1 | 3.0 | ± | 0.6 | 2.6 | ± | 0.5 | 1.5 | ± | 0.4 | 3.3 | ± | 0.7 | 1.8 | ± | 0.5 | 2.9 | ± | 0.7 | 2.5 | ± | 0.4 | 2.2 | ± | 0.5 |
| 1-2.5 | 2.3 | ± | 0.8 | 2.3 | ± | 0.4 | 2.1 | ± | 0.7 | 2.8 | ± | 0.5 | 1.4 | ± | 0.2 | 1.6 | ± | 0.4 | 2.2 | ± | 0.3 | 2.2 | ± | 0.4 |
| 2.5-5 | 0.8 | ± | 0.5 | 2.0 | ± | 0.4 | 0.9 | ± | 0.4 | 1.0 | ± | 0.3 | 1.4 | ± | 0.4 | 0.9 | ± | 0.4 | 1.6 | ± | 0.4 | 1.8 | ± | 0.6 |
| 5-10 | 0.3 | ± | 0.2 | 1.2 | ± | 0.5 | 2.0 | ± | 0.5 | 1.7 | ± | 0.4 | 1.2 | ± | 0.3 | 1.3 | ± | 0.2 α | 0.8 | ± | 0.3 | 1.3 | ± | 0.5 |
| 10-20 | 1.3 | ± | 0.5 | 0.6 | ± | 0.2 | 0.8 | ± | 0.2 | 0.8 | ± | 0.4 | 1.1 | ± | 0.4 | 0.8 | ± | 0.3 | 0.7 | ± | 0.3 | 0.6 | ± | 0.2 |
| 20-40 | 0.5 | ± | 0.3 | 0.4 | ± | 0.2 | 0.3 | ± | 0.2 | 0.3 | ± | 0.2 | 0.3 | ± | 0.2 | 0.3 | ± | 0.2 | 0.3 | ± | 0.2 | 0.3 | ± | 0.2 |
| 40-∞ | 0.3 | ± | 0.2 | 0.1 | ± | 0.1 | 0.0 | ± | 0.0 | 0.2 | ± | 0.2 | 0.2 | ± | 0.1 | 0.1 | ± | 0.1 | 0.1 | ± | 0.1 | 0.0 | ± | 0.0 |

**Supplementary Table S6:** Percentage (±SEM) of **wakefulness** amount in various wake episode lengths to the total amount of wakefulness and number of episodes (± SEM) of wakefulness in various wake episode lengths during the **ZT3-6** periods in PZ^Vgat-hM3Dq^ mice following **C21** administration as compared with control administration. N = 6 mice. α: p < 0.05, two-way ANOVA followed by a post hoc Bonferroni test.

| Percentage (±SEM) of wakefulness amount in various wake episode lengths to the total amount of wakefulness during the ZT3-6 period following C21 administration as compared with control administration |  |  |  |  |  |  |  |  |  |  |  |  |  |  |  |  |  |  |  |  |  |  |  |  |
| --- | --- | --- | --- | --- | --- | --- | --- | --- | --- | --- | --- | --- | --- | --- | --- | --- | --- | --- | --- | --- | --- | --- | --- | --- |
| Episode length (min) | Control |  |  | 1<sup>st</sup> 2 administrations |  |  | 1<sup>st</sup> month |  |  | 2<sup>nd</sup> month |  |  | 3<sup>rd</sup> month |  |  | 4<sup>th</sup> month |  |  | 5<sup>th</sup> month |  |  | 6<sup>th</sup> month |  |  |
|  | Avg | ± | SEM | Avg | ± | SEM | Avg | ± | SEM | Avg | ± | SEM | Avg | ± | SEM | Avg | ± | SEM | Avg | ± | SEM | Avg | ± | SEM |
| 0.1-0.5 | 15.0 | ± | 1.7 | 3.3 | ± | 1.2 | 11.8 | ± | 2.5 | 12.3 | ± | 1.3 | 10.3 | ± | 1.7 | 14.0 | ± | 4.4 | 12.8 | ± | 1.2 | 10.7 | ± | 0.9 |
| 0.5-1 | 3.0 | ± | 0.6 | 12.2 | ± | 8.0 | 2.8 | ± | 0.9 | 2.6 | ± | 0.6 | 2.0 | ± | 0.4 | 6.1 | ± | 1.9 | 3.2 | ± | 0.5 | 2.7 | ± | 0.4 |
| 1-2.5 | 2.3 | ± | 0.8 | 65.6 | ± | 16.2 | 1.0 | ± | 0.2 | 2.1 | ± | 0.6 | 1.3 | ± | 0.4 | 6.0 | ± | 1.7 | 2.6 | ± | 0.4 | 1.7 | ± | 0.3 |
| 2.5-5 | 0.8 | ± | 0.5 | 18.8 | ± | 9.2 | 1.3 | ± | 0.4 | 0.5 | ± | 0.0 | 0.9 | ± | 0.2 | 7.9 | ± | 3.4 | 0.9 | ± | 0.2 | 1.0 | ± | 0.3 |
| 5-10 | 0.3 | ± | 0.2 | 0.0 | ± | 0.0 | 1.8 | ± | 0.5 | 1.3 | ± | 0.3 | 1.4 | ± | 0.5 α | 24.4 | ± | 4.6 | 1.3 | ± | 0.5 | 0.8 | ± | 0.4 |
| 10-20 | 1.3 | ± | 0.5 | 0.0 | ± | 0.0 | 0.8 | ± | 0.3 | 1.1 | ± | 0.3 | 0.7 | ± | 0.2 | 26.1 | ± | 9.8 | 0.3 | ± | 0.2 | 0.8 | ± | 0.3 |
| 20-40 | 0.5 | ± | 0.3 | 0.0 | ± | 0.0 | 0.8 | ± | 0.3 | 0.3 | ± | 0.2 | 0.3 | ± | 0.2 | 15.6 | ± | 7.2 | 0.1 | ± | 0.1 | 0.4 | ± | 0.2 |
| 40-∞ | 0.3 | ± | 0.2 | 0.0 | ± | 0.0 | 0.0 | ± | 0.0 | 0.1 | ± | 0.1 | 0.0 | ± | 0.0 | 0.0 | ± | 0.0 | 0.3 | ± | 0.2 | 0.2 | ± | 0.1 |
| Number of episodes (± SEM) of wakefulness in various wake episode lengths during the ZT3-6 period following C21 administration as compared with control administration |  |  |  |  |  |  |  |  |  |  |  |  |  |  |  |  |  |  |  |  |  |  |  |  |
| Episode length (min) | Control |  |  | 1<sup>st</sup> 2 administrations |  |  | 1<sup>st</sup> month |  |  | 2<sup>nd</sup> month |  |  | 3<sup>rd</sup> month |  |  | 4<sup>th</sup> month |  |  | 5<sup>th</sup> month |  |  | 6<sup>th</sup> month |  |  |
|  | Avg | ± | SEM | Avg | ± | SEM | Avg | ± | SEM | Avg | ± | SEM | Avg | ± | SEM | Avg | ± | SEM | Avg | ± | SEM | Avg | ± | SEM |
| 0.1-0.5 | 7.1 | ± | 1.4 | 0.4 | ± | 0.2 | 7.1 | ± | 1.8 | 9.5 | ± | 2.0 | 12.1 | ± | 4.6 | 10.8 | ± | 1.6 | 16.4 | ± | 3.7 | 8.4 | ± | 2.0 |
| 0.5-1 | 3.7 | ± | 0.9 | 0.3 | ± | 0.2 | 4.2 | ± | 1.2 | 4.6 | ± | 0.6 | 5.6 | ± | 2.0 | 2.0 | ± | 0.4 | 9.8 | ± | 3.6 | 6.6 | ± | 2.3 |
| 1-2.5 | 6.3 | ± | 2.2 | 1.5 | ± | 0.5 | 4.0 | ± | 1.3 | 8.6 | ± | 2.8 | 6.0 | ± | 1.7 | 1.3 | ± | 0.4 | 15.1 | ± | 2.7 | 8.2 | ± | 3.4 |
| 2.5-5 | 4.8 | ± | 2.4 | 0.4 | ± | 0.2 | 9.3 | ± | 2.6 | 5.4 | ± | 1.1 | 9.4 | ± | 3.4 | 0.8 | ± | 0.2 | 10.6 | ± | 2.9 | 10.9 | ± | 3.8 |
| 5-10 | 3.4 | ± | 2.3 | 0.0 | ± | 0.0 | 24.8 | ± | 7.5 | 23.7 | ± | 6.0 | 22.4 | ± | 3.9 | 1.3 | ± | 0.5 | 25.8 | ± | 7.3 | 12.4 | ± | 6.1 |
| 10-20 | 31.5 | ± | 12.8 | 0.0 | ± | 0.0 | 19.6 | ± | 5.1 | 34.0 | ± | 8.8 | 28.9 | ± | 8.7 | 0.6 | ± | 0.2 | 7.2 | ± | 5.8 | 22.4 | ± | 8.8 |
| 20-40 | 22.6 | ± | 14.9 | 0.0 | ± | 0.0 | 31.1 | ± | 12.0 | 9.8 | ± | 6.8 | 15.6 | ± | 7.2 | 0.3 | ± | 0.2 | 2.2 | ± | 2.2 | 19.6 | ± | 10.0 |
| 40-∞ | 20.7 | ± | 13.1 | 0.0 | ± | 0.0 | 0.0 | ± | 0.0 | 4.6 | ± | 4.6 | 0.0 | ± | 0.0 | 0.0 | ± | 0.0 | 12.9 | ± | 8.5 | 11.5 | ± | 7.5 |

**Supplementary Table S7:**
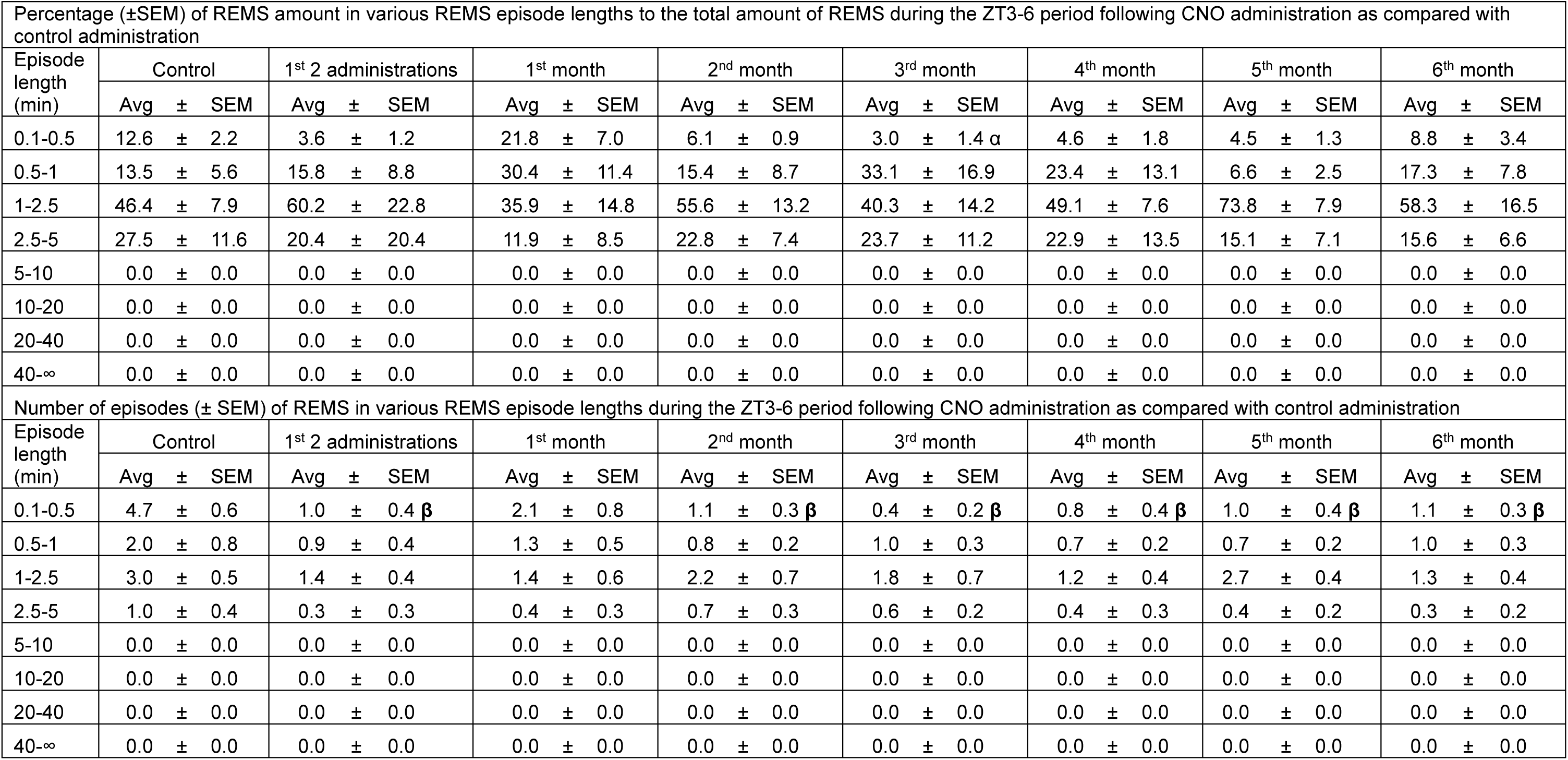
Percentage (±SEM) of **REMS** amount in various REMS episode lengths to the total amount of REMS and number of episodes (± SEM) of REMS in various REMS episode lengths during the **ZT3-6** periods in PZ^Vgat-hM3Dq^ mice following **CNO** administration as compared with control administration. N = 6 mice. β: p < 0.01, two-way ANOVA followed by a post hoc Bonferroni test.

**Supplementary Table S8:**
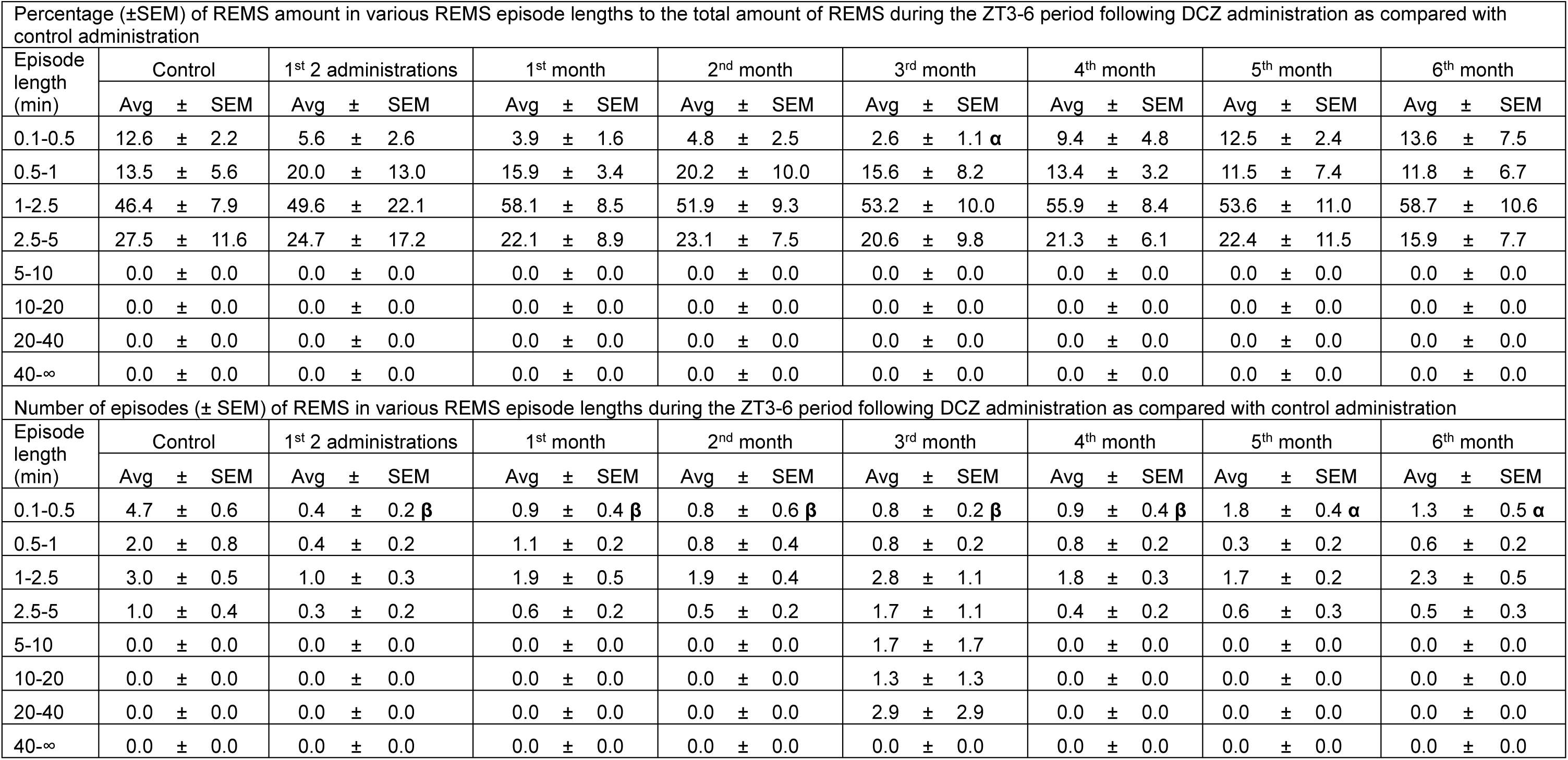
Percentage (±SEM) of **REMS** amount in various REMS episode lengths to the total amount of REMS and number of episodes (± SEM) of REMS in various REMS episode lengths during the **ZT3-6** periods in PZ^Vgat-hM3Dq^ mice following **DCZ** administration as compared with control administration. N = 6 mice. α: p < 0.05, β: p < 0.01, two-way ANOVA followed by a post hoc Bonferroni test.

**Supplementary Table S9:**
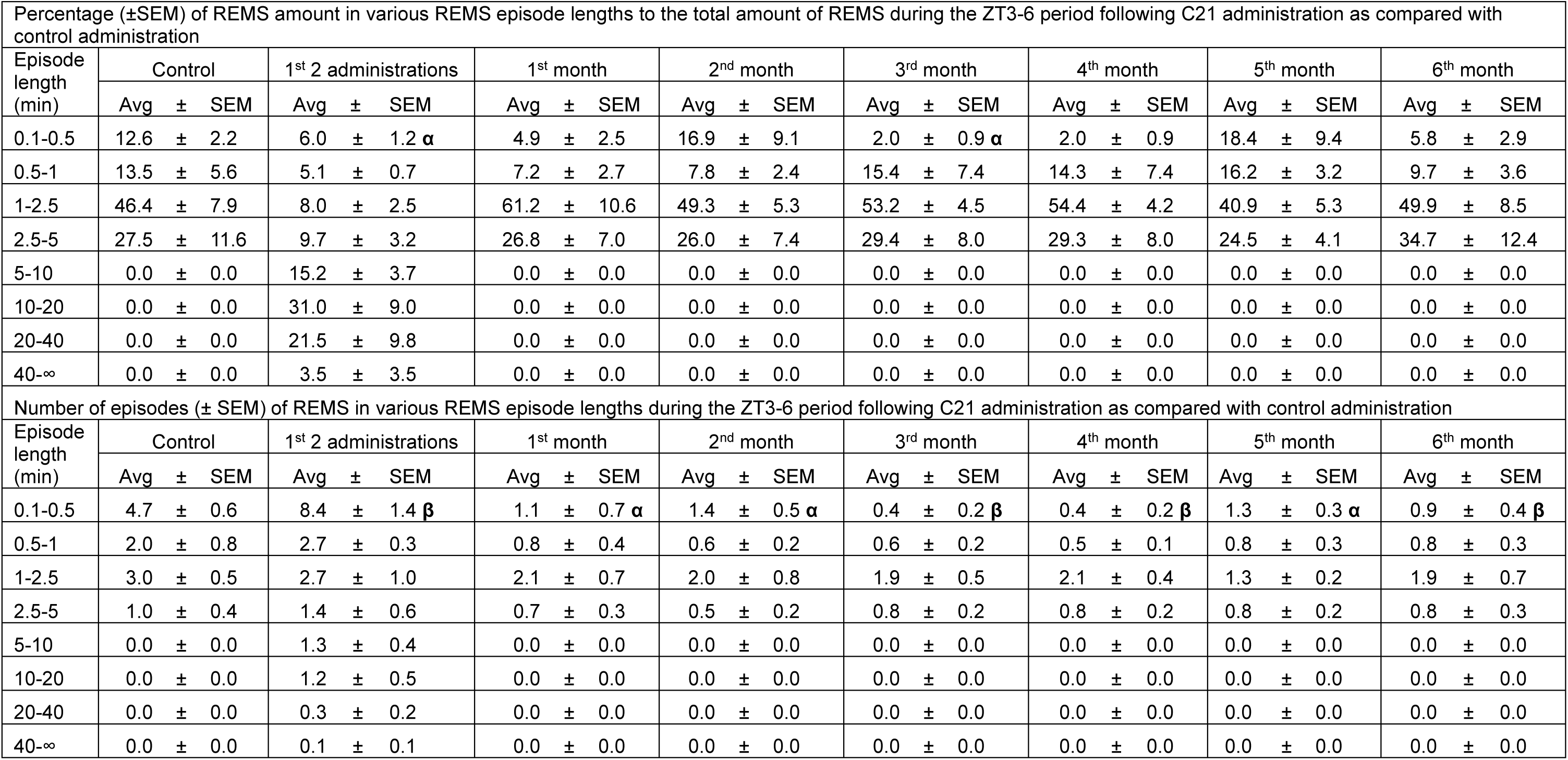
Percentage (±SEM) of **REMS** amount in various REMS episode lengths to the total amount of REMS and number of episodes (± SEM) of REMS in various REMS episode lengths during the **ZT3-6** periods in PZ^Vgat-hM3Dq^ mice following **C21** administration as compared with control administration. N = 6 mice. α: p < 0.05, β: p < 0.01, two-way ANOVA followed by a post hoc Bonferroni test.

**Supplementary Table S10:**
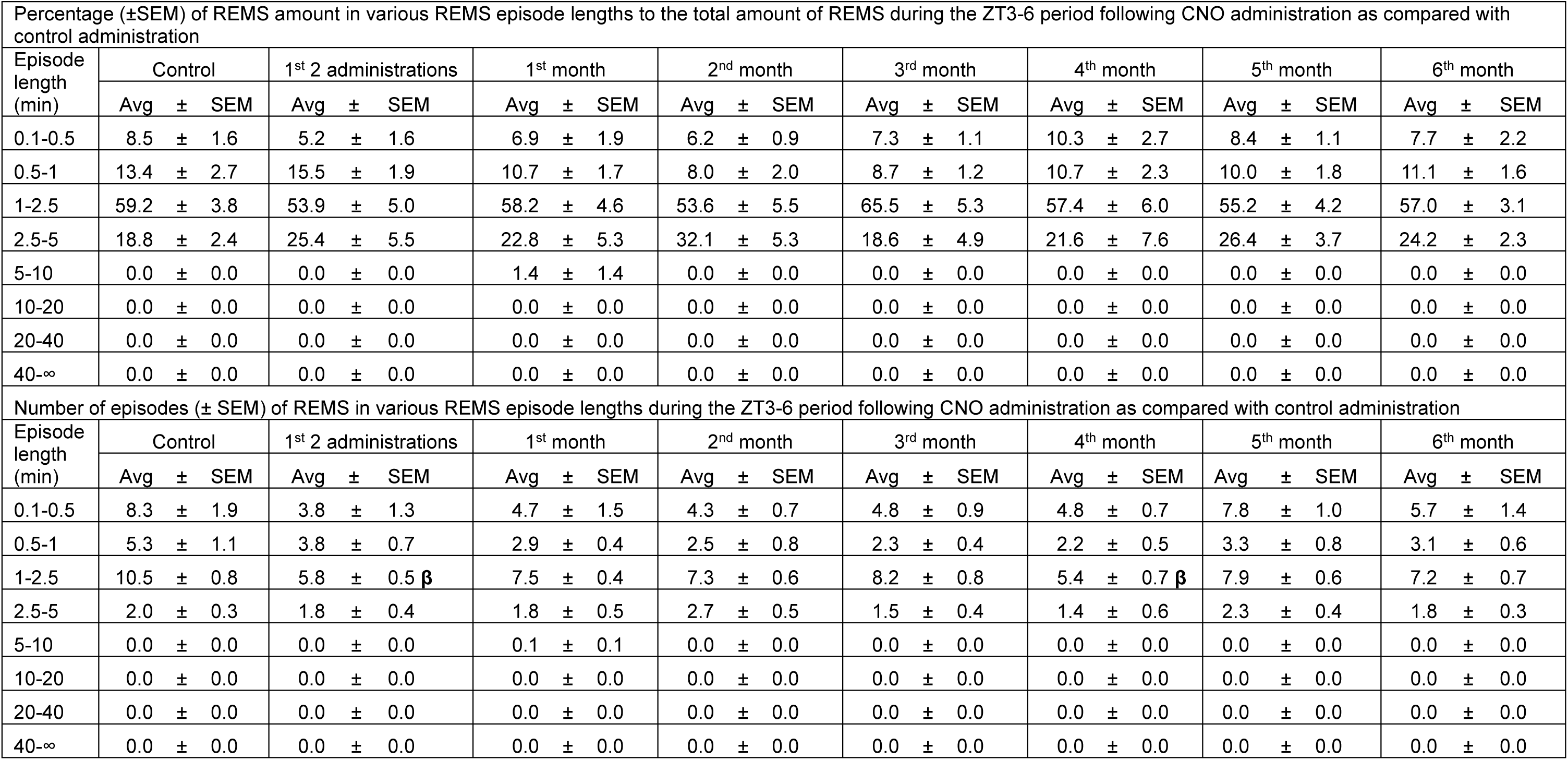
Percentage (±SEM) of **REMS** amount in various REMS episode lengths to the total amount of REMS and number of episodes (± SEM) of REMS in various REMS episode lengths during the **ZT6-12** periods in PZ^Vgat-hM3Dq^ mice following **CNO** administration as compared with control administration. N = 6 mice. β: p < 0.01, two-way ANOVA followed by a post hoc Bonferroni test.

**Supplementary Table S11:**
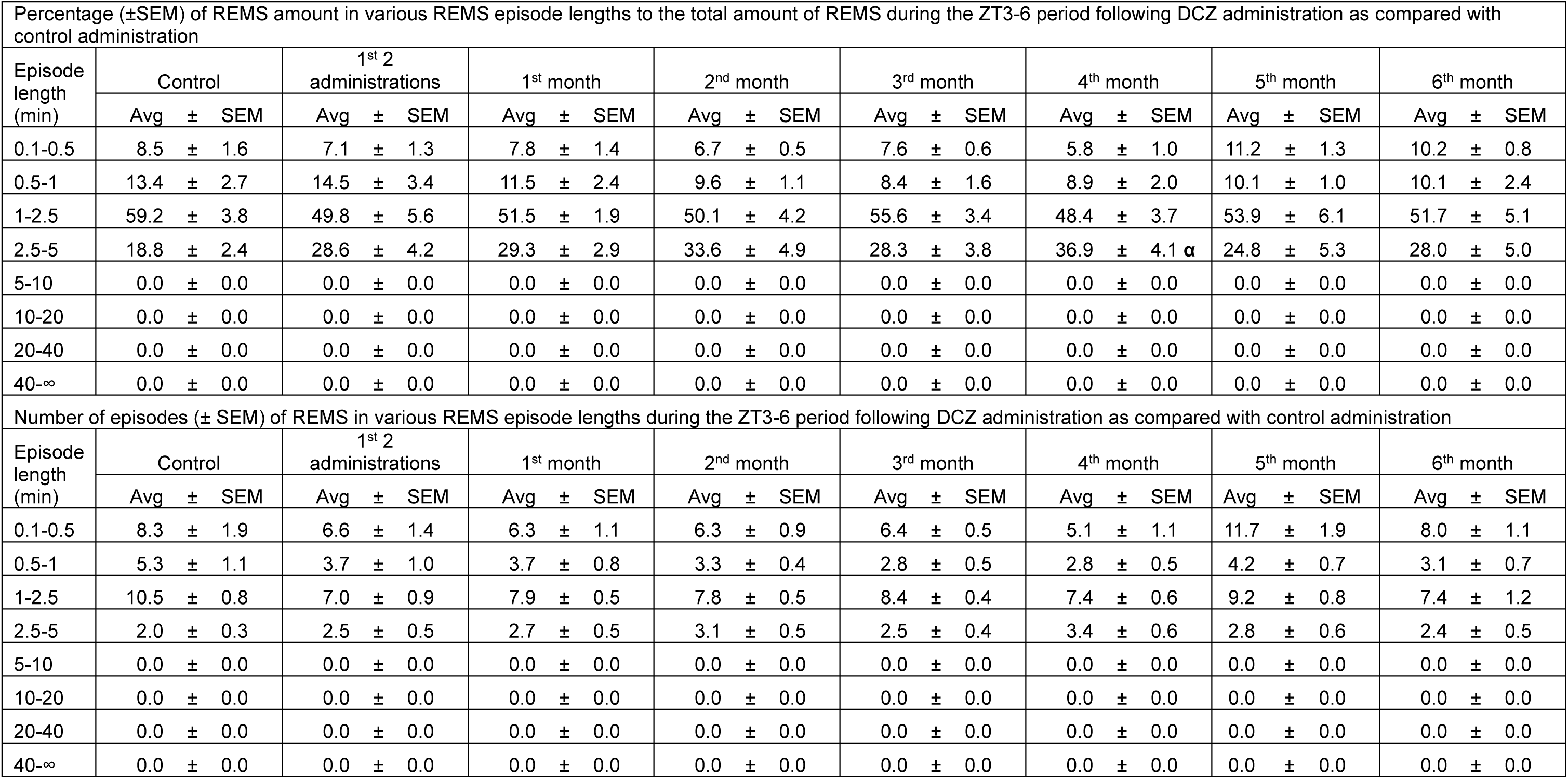
Percentage (±SEM) of **REMS** amount in various REMS episode lengths to the total amount of REMS and number of episodes (± SEM) of REMS in various REMS episode lengths during the **ZT6-12** periods in PZ^Vgat-hM3Dq^ mice following **DCZ** administration as compared with control administration. N = 6 mice. α: p < 0.05, two-way ANOVA followed by a post hoc Bonferroni test.

**Supplementary Table S12:**
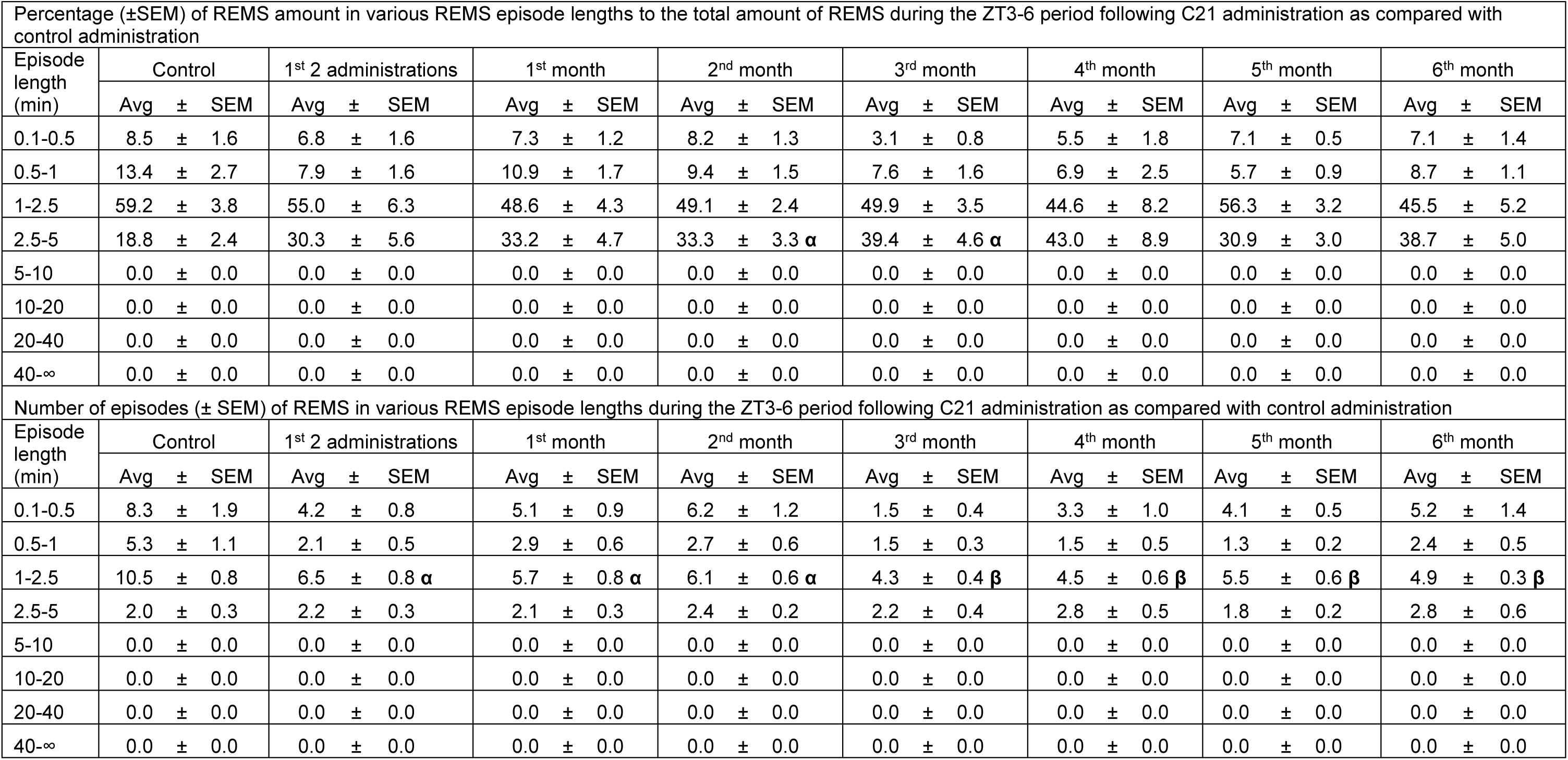
Percentage (±SEM) of **REMS** amount in various REMS episode lengths to the total amount of REMS and number of episodes (± SEM) of REMS in various REMS episode lengths during the **ZT6-12** periods in PZ^Vgat-hM3Dq^ mice following **C21** administration as compared with control administration. N = 6 mice. α: p < 0.05, β: p < 0.01, two-way ANOVA followed by a post hoc Bonferroni test.

**Supplementary Table S13:**
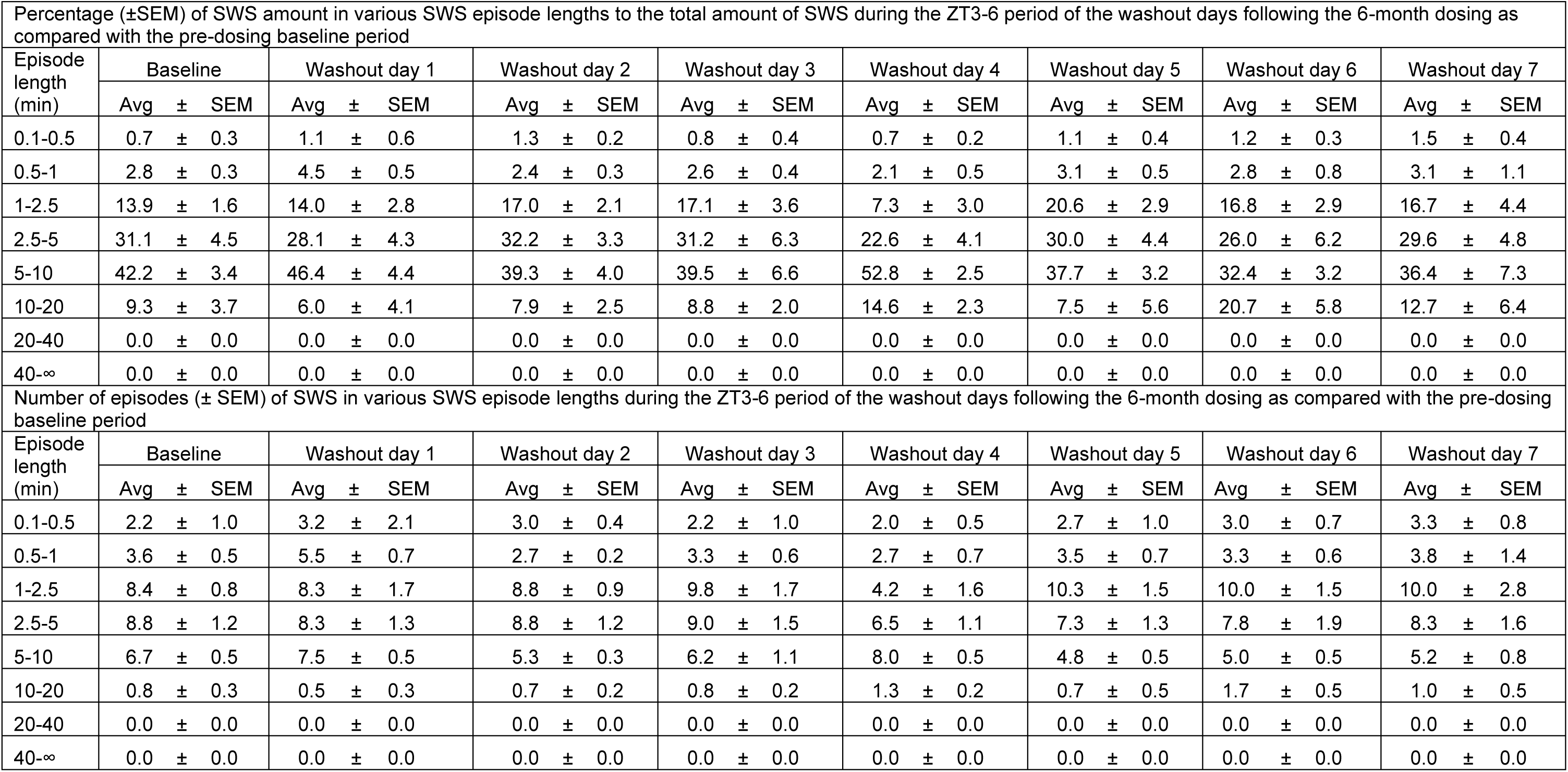
Percentage (±SEM) of **SWS** amount in various SWS episode lengths to the total amount of SWS and number of episodes (± SEM) of SWS in various SWS episode lengths during **the ZT3-6 period of the washout days following the 6-month dosing** in PZ^Vgat-hM3Dq^ mice as compared with the pre-dosing baseline period. N = 6 mice. No significance, two-way ANOVA followed by a post hoc Bonferroni test.

**Supplementary Table S14:** Percentage (±SEM) of **wakefulness** amount in various wake episode lengths to the total amount of wakefulness and number of episodes (± SEM) of wakefulness in various wake episode lengths during **the ZT3-6 period of the washout days following the 6-month dosing** in PZ^Vgat-hM3Dq^ mice as compared with the pre-dosing baseline period. N = 6 mice. α: p < 0.05, β: p < 0.01, two-way ANOVA followed by a post hoc Bonferroni test.

| Percentage (±SEM) of wakefulness amount in various wake episode lengths to the total amount of wakefulness during the ZT3-6 period of the washout days following the 6-month dosing as compared with the pre-dosing baseline period |  |  |  |  |  |  |  |  |  |  |  |  |  |  |  |  |  |  |  |  |  |  |  |  |
| --- | --- | --- | --- | --- | --- | --- | --- | --- | --- | --- | --- | --- | --- | --- | --- | --- | --- | --- | --- | --- | --- | --- | --- | --- |
| Episode length (min) | Baseline |  |  | Washout day 1 |  |  | Washout day 2 |  |  | Washout day 3 |  |  | Washout day 4 |  |  | Washout day 5 |  |  | Washout day 6 |  |  | Washout day 7 |  |  |
|  | Avg | ± | SEM | Avg | ± | SEM | Avg | ± | SEM | Avg | ± | SEM | Avg | ± | SEM | Avg | ± | SEM | Avg | ± | SEM | Avg | ± | SEM |
| 0.1-0.5 | 10.6 | ± | 1.2 | 12.8 | ± | 2.0 | 10.3 | ± | 1.7 | 12.1 | ± | 0.7 | 11.9 | ± | 2.1 | 8.3 | ± | 1.5 | 11.4 | ± | 2.5 | 11.5 | ± | 2.4 |
| 0.5-1 | 5.8 | ± | 0.7 | 7.4 | ± | 1.6 | 5.6 | ± | 2.3 | 6.9 | ± | 1.5 | 2.5 | ± | 0.5 α | 4.8 | ± | 1.8 | 4.8 | ± | 1.4 | 6.2 | ± | 2.7 |
| 1-2.5 | 3.8 | ± | 1.2 | 3.6 | ± | 1.0 | 8.2 | ± | 2.1 | 8.3 | ± | 1.7 | 1.8 | ± | 1.4 | 4.9 | ± | 1.3 | 7.6 | ± | 3.1 | 3.9 | ± | 2.0 |
| 2.5-5 | 5.6 | ± | 1.7 | 3.2 | ± | 1.4 | 1.6 | ± | 1.6 | 5.2 | ± | 2.7 | 9.7 | ± | 5.1 | 2.3 | ± | 1.4 | 3.2 | ± | 2.1 | 7.7 | ± | 2.5 |
| 5-10 | 18.3 | ± | 2.6 | 26.1 | ± | 7.1 | 21.7 | ± | 8.7 | 16.0 | ± | 4.2 | 15.1 | ± | 7.6 | 10.0 | ± | 6.1 | 7.3 | ± | 5.9 | 7.3 | ± | 5.0 |
| 10-20 | 16.3 | ± | 5.9 | 19.6 | ± | 6.6 | 21.7 | ± | 10.0 | 16.8 | ± | 8.1 | 0.0 | ± | 0.0 | 0.0 | ± | 0.0 | 7.8 | ± | 5.1 | 9.9 | ± | 4.6 |
| 20-40 | 28.7 | ± | 10.8 | 27.3 | ± | 13.5 | 22.1 | ± | 10.1 | 34.8 | ± | 11.2 | 32.8 | ± | 14.8 | 30.4 | ± | 15.3 | 36.5 | ± | 17.4 | 18.9 | ± | 12.8 |
| 40-∞ | 11.0 | ± | 7.2 | 0.0 | ± | 0.0 | 8.8 | ± | 8.8 | 0.0 | ± | 0.0 | 26.2 | ± | 16.7 | 39.4 | ± | 17.9 | 21.4 | ± | 13.7 | 34.6 | ± | 16.0 |
| Number of episodes (± SEM) of wakefulness in various wake episode lengths during the ZT3-6 period of the washout days following the 6-month dosing as compared with the pre-dosing baseline period |  |  |  |  |  |  |  |  |  |  |  |  |  |  |  |  |  |  |  |  |  |  |  |  |
| Episode length (min) | Baseline |  |  | Washout day 1 |  |  | Washout day 2 |  |  | Washout day 3 |  |  | Washout day 4 |  |  | Washout day 5 |  |  | Washout day 6 |  |  | Washout day 7 |  |  |
|  | Avg | ± | SEM | Avg | ± | SEM | Avg | ± | SEM | Avg | ± | SEM | Avg | ± | SEM | Avg | ± | SEM | Avg | ± | SEM | Avg | ± | SEM |
| 0.1-0.5 | 20.7 | ± | 2.7 | 22.8 | ± | 3.9 | 19.7 | ± | 1.2 | 20.7 | ± | 1.7 | 19.0 | ± | 2.6 | 20.3 | ± | 3.0 | 22.2 | ± | 1.9 | 22.2 | ± | 2.6 |
| 0.5-1 | 4.3 | ± | 0.4 | 5.2 | ± | 0.9 | 3.8 | ± | 1.2 | 4.7 | ± | 1.3 | 1.7 | ± | 0.3 β | 4.8 | ± | 1.7 | 3.7 | ± | 1.3 | 4.5 | ± | 1.8 |
| 1-2.5 | 1.4 | ± | 0.4 | 1.3 | ± | 0.3 | 2.8 | ± | 0.5 | 2.3 | ± | 0.4 | 0.5 | ± | 0.3 | 2.0 | ± | 0.4 | 2.5 | ± | 0.7 | 1.5 | ± | 0.8 |
| 2.5-5 | 0.9 | ± | 0.3 | 0.5 | ± | 0.2 | 0.3 | ± | 0.3 | 0.7 | ± | 0.3 | 1.0 | ± | 0.5 | 0.5 | ± | 0.3 | 0.3 | ± | 0.2 | 1.3 | ± | 0.4 |
| 5-10 | 1.4 | ± | 0.2 | 1.8 | ± | 0.5 | 1.7 | ± | 0.5 | 1.3 | ± | 0.3 | 0.8 | ± | 0.3 | 0.8 | ± | 0.5 | 0.5 | ± | 0.3 | 0.7 | ± | 0.4 |
| 10-20 | 0.8 | ± | 0.3 | 0.8 | ± | 0.3 | 0.8 | ± | 0.3 | 0.5 | ± | 0.2 | 0.0 | ± | 0.0 | 0.0 | ± | 0.0 | 0.3 | ± | 0.2 | 0.5 | ± | 0.2 |
| 20-40 | 0.7 | ± | 0.2 | 0.7 | ± | 0.3 | 0.5 | ± | 0.2 | 0.7 | ± | 0.2 | 0.5 | ± | 0.2 | 0.7 | ± | 0.3 | 0.8 | ± | 0.4 | 0.5 | ± | 0.3 |
| 40-∞ | 0.2 | ± | 0.1 | 0.0 | ± | 0.0 | 0.2 | ± | 0.2 | 0.0 | ± | 0.0 | 0.3 | ± | 0.2 | 0.7 | ± | 0.3 | 0.3 | ± | 0.2 | 0.5 | ± | 0.2 |

**Supplementary Table S15:**
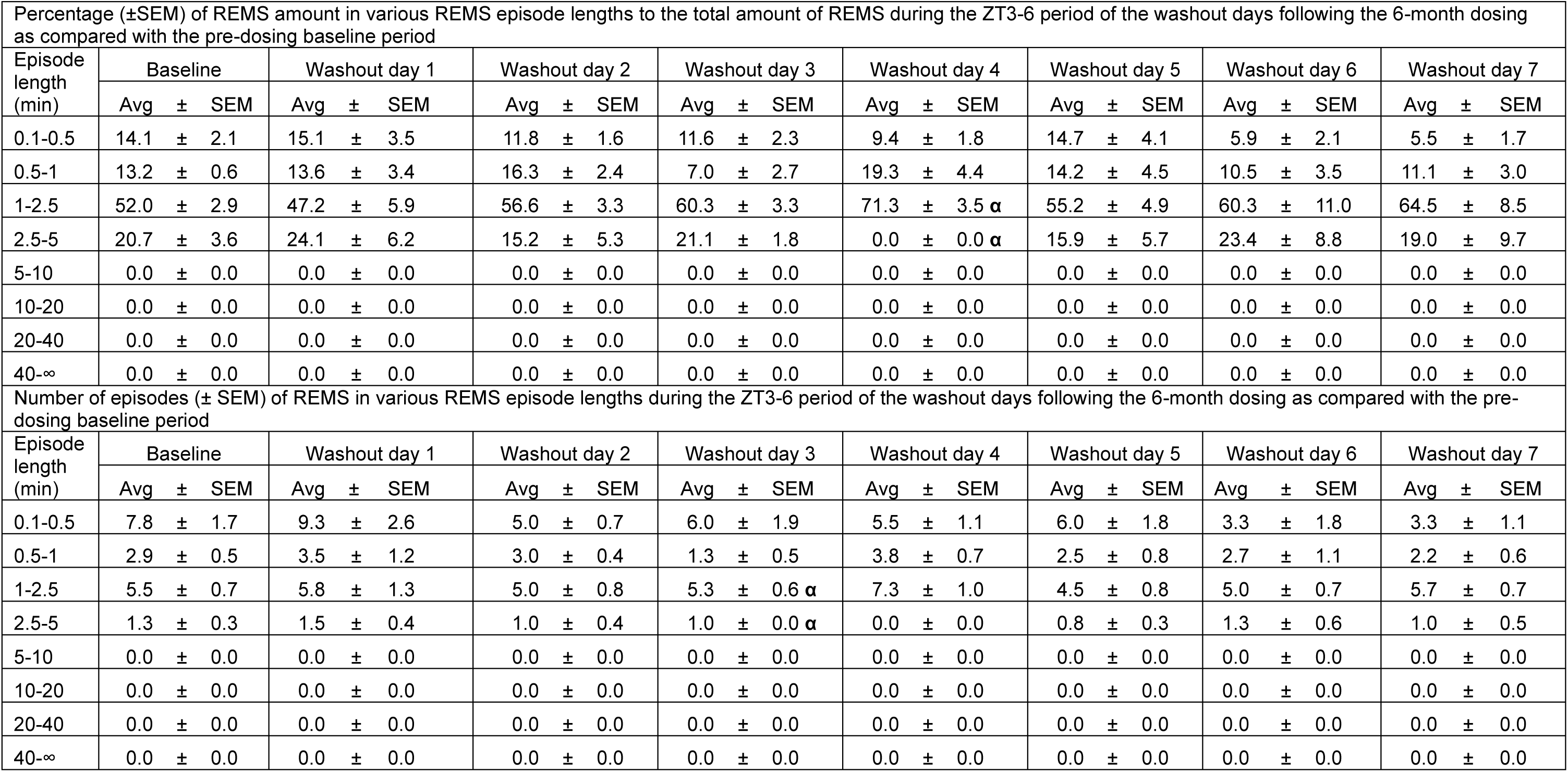
Percentage (±SEM) of **REMS** amount in various REMS episode lengths to the total amount of REMS and number of episodes (± SEM) of REMS in various REMS episode lengths during **the ZT3-6 period of the washout days following the 6-month dosing** in PZ^Vgat-hM3Dq^ mice as compared with the pre-dosing baseline period. N = 6 mice. α: p < 0.05, two-way ANOVA followed by a post hoc Bonferroni test.

**Supplementary Table S16:**
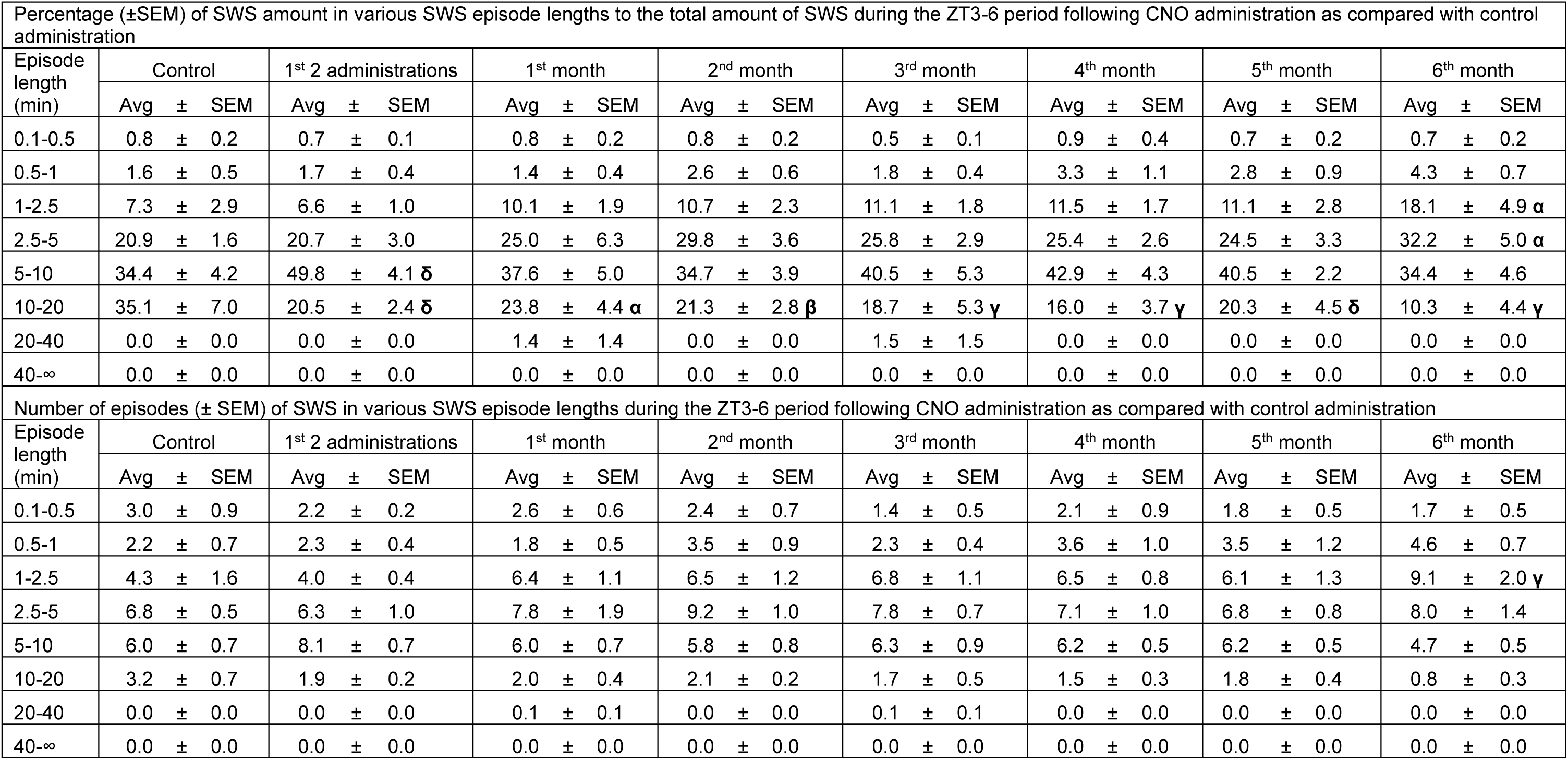
Percentage (±SEM) of **SWS** amount in various SWS episode lengths to the total amount of SWS and number of episodes (± SEM) of SWS in various SWS episode lengths during the **ZT3-6** periods in control mice following **CNO** administration as compared with control administration. N = 6 mice. α: p < 0.05, β: p < 0.01, δ: p < 0.001, γ: p < 0.0001, two-way ANOVA followed by a post hoc Bonferroni test.

**Supplementary Table S17:** Percentage (±SEM) of **SWS** amount in various SWS episode lengths to the total amount of SWS and number of episodes (± SEM) of SWS in various SWS episode lengths during the **ZT3-6** periods in control mice following **DCZ** administration as compared with control administration. N = 6 mice. α: p < 0.05, β: p < 0.01, δ: p < 0.001, γ: p < 0.0001, two-way ANOVA followed by a post hoc Bonferroni test.

| Percentage (±SEM) of SWS amount in various SWS episode lengths to the total amount of SWS during the ZT3-6 period following DCZ administration as compared with control administration | | | | | | | | |
| --- | --- | --- | --- | --- | --- | --- | --- | --- |
| Episode length (min) | Control | 1<sup>st</sup> 2 administrations | 1<sup>st</sup> month | 2<sup>nd</sup> month | 3<sup>rd</sup> month | 4<sup>th</sup> month | 5<sup>th</sup> month | 6<sup>th</sup> month |
| | Avg ± SEM | Avg ± SEM | Avg ± SEM | Avg ± SEM | Avg ± SEM | Avg ± SEM | Avg ± SEM | Avg ± SEM |
| 0.1-0.5 | 0.8 ± 0.2 | 0.7 ± 0.2 | 0.6 ± 0.1 | 0.4 ± 0.1 | 0.6 ± 0.1 | 0.9 ± 0.4 | 0.5 ± 0.2 | 0.9 ± 0.2 |
| 0.5-1 | 1.6 ± 0.5 | 1.0 ± 0.2 | 1.1 ± 0.2 | 1.7 ± 0.5 | 2.5 ± 0.7 | 3.2 ± 0.7 | 2.2 ± 0.3 | 2.8 ± 1.0 |
| 1-2.5 | 7.3 ± 2.9 | 6.1 ± 1.3 | 9.5 ± 1.4 | 8.7 ± 2.0 | 13.1 ± 3.1 | 8.9 ± 1.3 | 12.6 ± 3.2 | 15.0 ± 3.4 |
| 2.5-5 | 20.9 ± 1.6 | 14.1 ± 3.3 | 19.5 ± 2.4 | 21.8 ± 1.3 | 22.7 ± 3.4 | 22.3 ± 2.9 | 27.1 ± 3.6 | 30.6 ± 5.9 |
| 5-10 | 34.4 ± 4.2 | 38.3 ± 2.9 | 49.7 ± 3.5 $\delta$ | 44.8 ± 4.2 $\alpha$ | 41.1 ± 2.9 | 37.4 ± 4.1 | 40.0 ± 5.3 | 35.8 ± 6.9 |
| 10-20 | 35.1 ± 7.0 | 38.4 ± 3.4 | 19.6 ± 4.1 $\delta$ | 22.6 ± 4.8 $\beta$ | 16.4 ± 3.7 $\gamma$ | 20.1 ± 4.3 $\delta$ | 16.1 ± 3.8 $\gamma$ | 14.9 ± 7.0 $\gamma$ |
| 20-40 | 0.0 ± 0.0 | 1.4 ± 1.4 | 0.0 ± 0.0 | 0.0 ± 0.0 | 3.7 ± 3.7 | 3.2 ± 2.1 | 1.5 ± 1.5 | 0.0 ± 0.0 |
| 40-∞ | 0.0 ± 0.0 | 0.0 ± 0.0 | 0.0 ± 0.0 | 0.0 ± 0.0 | 0.0 ± 0.0 | 4.0 ± 4.0 | 0.0 ± 0.0 | 0.0 ± 0.0 |
| Number of episodes (± SEM) of SWS in various SWS episode lengths during the ZT3-6 period following DCZ administration as compared with control administration | | | | | | | | |
| Episode length (min) | Control | 1<sup>st</sup> 2 administrations | 1<sup>st</sup> month | 2<sup>nd</sup> month | 3<sup>rd</sup> month | 4<sup>th</sup> month | 5<sup>th</sup> month | 6<sup>th</sup> month |
| | Avg ± SEM | Avg ± SEM | Avg ± SEM | Avg ± SEM | Avg ± SEM | Avg ± SEM | Avg ± SEM | Avg ± SEM |
| 0.1-0.5 | 3.0 ± 0.9 | 2.2 ± 0.5 | 2.1 ± 0.5 | 1.3 ± 0.4 | 1.7 ± 0.4 | 1.9 ± 0.6 | 1.3 ± 0.5 | 2.2 ± 0.5 |
| 0.5-1 | 2.2 ± 0.7 | 1.4 ± 0.3 | 1.7 ± 0.3 | 2.3 ± 0.6 | 3.1 ± 0.7 | 3.6 ± 0.7 | 2.8 ± 0.4 | 3.4 ± 1.0 |
| 1-2.5 | 4.3 ± 1.6 | 4.0 ± 1.0 | 6.1 ± 0.9 | 5.3 ± 1.2 | 7.3 ± 1.6 $\alpha$ | 4.8 ± 0.7 | 7.5 ± 1.8 $\beta$ | 8.6 ± 1.7 $\gamma$ |
| 2.5-5 | 6.8 ± 0.5 | 4.3 ± 1.1 $\alpha$ | 6.4 ± 0.8 | 7.0 ± 0.3 | 6.6 ± 1.2 | 6.1 ± 0.8 | 8.2 ± 1.0 | 8.5 ± 1.3 |
| 5-10 | 6.0 ± 0.7 | 6.1 ± 0.4 | 7.9 ± 0.5 | 7.8 ± 0.9 | 6.3 ± 0.5 | 5.7 ± 0.9 | 6.3 ± 0.9 | 5.5 ± 1.0 |
| 10-20 | 3.2 ± 0.7 | 3.5 ± 0.3 | 1.9 ± 0.5 | 2.2 ± 0.4 | 1.4 ± 0.3 | 1.8 ± 0.5 | 1.3 ± 0.3 | 1.2 ± 0.6 |
| 20-40 | 0.0 ± 0.0 | 0.1 ± 0.1 | 0.0 ± 0.0 | 0.0 ± 0.0 | 0.1 ± 0.1 | 0.2 ± 0.1 | 0.1 ± 0.1 | 0.0 ± 0.0 |
| 40-∞ | 0.0 ± 0.0 | 0.0 ± 0.0 | 0.0 ± 0.0 | 0.0 ± 0.0 | 0.0 ± 0.0 | 0.1 ± 0.1 | 0.0 ± 0.0 | 0.0 ± 0.0 |

**Supplementary Table S18:** Percentage (±SEM) of **SWS** amount in various SWS episode lengths to the total amount of SWS and number of episodes (± SEM) of SWS in various SWS episode lengths during the **ZT3-6** periods in control mice following **C21** administration as compared with control administration. N = 6 mice. α: p < 0.05, β: p < 0.01, δ: p < 0.001, γ: p < 0.0001, two-way ANOVA followed by a post hoc Bonferroni test.

| Percentage (±SEM) of SWS amount in various SWS episode lengths to the total amount of SWS during the ZT3-6 period following C21 administration as compared with control administration | | | | | | | | | |
| --- | --- | --- | --- | --- | --- | --- | --- | --- | --- |
| Episode length (min) | Control | 1<sup>st</sup> 2 administrations | 1<sup>st</sup> month | 2<sup>nd</sup> month | 3<sup>rd</sup> month | 4<sup>th</sup> month | 5<sup>th</sup> month | 6<sup>th</sup> month |  |
| | Avg ± SEM | Avg ± SEM | Avg ± SEM | Avg ± SEM | Avg ± SEM | Avg ± SEM | Avg ± SEM | Avg ± SEM | Avg ± SEM |
| 0.1-0.5 | 0.8 ± 0.2 | 0.4 ± 0.1 | 0.4 ± 0.1 | 0.6 ± 0.2 | 0.3 ± 0.1 | 0.2 ± 0.0 | 0.5 ± 0.2 | 0.4 ± 0.2 | |
| 0.5-1 | 1.6 ± 0.5 | 0.7 ± 0.2 | 1.0 ± 0.3 | 1.1 ± 0.2 | 0.8 ± 0.3 | 0.9 ± 0.3 | 1.7 ± 0.6 | 2.1 ± 0.9 | |
| 1-2.5 | 7.3 ± 2.9 | 3.5 ± 0.6 | 6.2 ± 2.3 | 4.6 ± 0.7 | 5.2 ± 1.0 | 6.1 ± 1.9 | 7.9 ± 2.5 | 8.9 ± 4.4 | |
| 2.5-5 | 20.9 ± 1.6 | 11.2 ± 0.8 | 14.9 ± 3.9 | 12.2 ± 2.4 | 15.4 ± 3.9 $\alpha$ | 16.4 ± 2.3 | 18.7 ± 5.9 | 25.4 ± 7.1 | |
| 5-10 | 34.4 ± 4.2 | 30.7 ± 3.9 | 28.8 ± 1.6 | 35.4 ± 6.5 | 32.8 ± 3.2 | 31.9 ± 3.9 | 31.9 ± 2.9 | 32.4 ± 6.0 | |
| 10-20 | 35.1 ± 7.0 | 48.8 ± 3.0 $\alpha$ | 34.7 ± 4.0 | 33.1 ± 6.2 | 37.7 ± 4.8 $\gamma$ | 31.0 ± 4.3 | 36.6 ± 9.8 | 27.5 ± 7.5 | |
| 20-40 | 0.0 ± 0.0 | 4.8 ± 2.2 | 14.0 ± 3.7 $\beta$ | 13.0 ± 5.4 $\alpha$ | 7.9 ± 4.3 | 10.5 ± 2.8 | 2.7 ± 1.7 | 3.4 ± 2.1 | |
| 40-∞ | 0.0 ± 0.0 | 0.0 ± 0.0 | 0.0 ± 0.0 | 0.0 ± 0.0 | 0.0 ± 0.0 | 3.0 ± 3.0 | 0.0 ± 0.0 | 0.0 ± 0.0 | |
| Number of episodes (± SEM) of SWS in various SWS episode lengths during the ZT3-6 period following C21 administration as compared with control administration | | | | | | | | | |
| Episode length (min) | Control | 1<sup>st</sup> 2 administrations | 1<sup>st</sup> month | 2<sup>nd</sup> month | 3<sup>rd</sup> month | 4<sup>th</sup> month | 5<sup>th</sup> month | 6<sup>th</sup> month |  |
| | Avg ± SEM | Avg ± SEM | Avg ± SEM | Avg ± SEM | Avg ± SEM | Avg ± SEM | Avg ± SEM | Avg ± SEM | Avg ± SEM |
| 0.1-0.5 | 3.0 ± 0.9 | 1.2 ± 0.3 | 1.3 ± 0.2 | 1.8 ± 0.4 | 1.3 ± 0.3 | 0.8 ± 0.1 | 1.3 ± 0.3 | 1.2 ± 0.6 | |
| 0.5-1 | 2.2 ± 0.7 | 1.1 ± 0.3 | 1.4 ± 0.4 | 1.8 ± 0.4 | 1.4 ± 0.4 | 1.3 ± 0.4 | 2.2 ± 0.6 | 2.7 ± 0.9 | |
| 1-2.5 | 4.3 ± 1.6 | 2.5 ± 0.4 | 4.3 ± 1.5 | 3.3 ± 0.5 | 3.8 ± 0.5 | 3.6 ± 0.9 | 4.5 ± 1.1 | 5.5 ± 2.2 | |
| 2.5-5 | 6.8 ± 0.5 | 3.7 ± 0.5 $\beta$ | 5.0 ± 1.2 | 4.3 ± 0.8 $\alpha$ | 6.3 ± 1.1 | 5.1 ± 0.6 | 5.8 ± 1.8 | 8.0 ± 1.8 | |
| 5-10 | 6.0 ± 0.7 | 5.2 ± 0.5 | 5.0 ± 0.4 | 6.0 ± 1.3 | 5.3 ± 0.4 | 5.4 ± 0.9 | 5.1 ± 0.4 | 5.4 ± 1.0 | |
| 10-20 | 3.2 ± 0.7 | 4.3 ± 0.4 | 3.3 ± 0.4 | 3.3 ± 0.7 | 3.4 ± 0.5 | 2.8 ± 0.4 | 3.3 ± 0.8 | 2.6 ± 0.7 | |
| 20-40 | 0.0 ± 0.0 | 0.3 ± 0.1 | 0.8 ± 0.2 | 0.7 ± 0.3 | 0.2 ± 0.1 | 0.6 ± 0.2 | 0.2 ± 0.1 | 0.2 ± 0.1 | |
| 40-∞ | 0.0 ± 0.0 | 0.0 ± 0.0 | 0.0 ± 0.0 | 0.0 ± 0.0 | 0.0 ± 0.0 | 0.1 ± 0.1 | 0.0 ± 0.0 | 0.0 ± 0.0 | |

**Supplementary Table S19:** Percentage (±SEM) of **wakefulness** amount in various wake episode lengths to the total amount of wakefulness and number of episodes (± SEM) of wakefulness in various wake episode lengths during the **ZT3-6** periods in control mice following **CNO** administration as compared with control administration. N = 6 mice. α: p < 0.05, β: p < 0.01, δ: p < 0.001, γ: p < 0.0001, two-way ANOVA followed by a post hoc Bonferroni test.

| Percentage (±SEM) of wakefulness amount in various wake episode lengths to the total amount of wakefulness during the ZT3-6 period following CNO administration as compared with control administration |  |  |  |  |  |  |  |  |  |  |  |  |  |  |  |  |  |  |  |  |  |  |  |  |
| --- | --- | --- | --- | --- | --- | --- | --- | --- | --- | --- | --- | --- | --- | --- | --- | --- | --- | --- | --- | --- | --- | --- | --- | --- |
| Episode length (min) | Control |  |  | 1<sup>st</sup> 2 administrations |  |  | 1<sup>st</sup> month |  |  | 2<sup>nd</sup> month |  |  | 3<sup>rd</sup> month |  |  | 4<sup>th</sup> month |  |  | 5<sup>th</sup> month |  |  | 6<sup>th</sup> month |  |  |
|  | Avg | ± | SEM | Avg | ± | SEM | Avg | ± | SEM | Avg | ± | SEM | Avg | ± | SEM | Avg | ± | SEM | Avg | ± | SEM | Avg | ± | SEM |
| 0.1-0.5 | 9.6 | ± | 1.3 | 9.5 | ± | 0.6 | 10.6 | ± | 0.7 | 13.3 | ± | 1.1 | 10.8 | ± | 2.0 | 11.6 | ± | 1.7 | 8.4 | ± | 1.6 | 10.9 | ± | 1.9 |
| 0.5-1 | 2.1 | ± | 0.6 | 2.4 | ± | 0.5 | 3.9 | ± | 1.7 | 5.2 | ± | 1.3 | 4.0 | ± | 0.7 | 4.4 | ± | 1.4 | 6.9 | ± | 1.8 | 4.4 | ± | 1.3 |
| 1-2.5 | 3.1 | ± | 1.2 | 2.3 | ± | 0.7 | 3.6 | ± | 1.2 | 2.6 | ± | 0.8 | 2.4 | ± | 0.7 | 3.3 | ± | 1.3 | 7.8 | ± | 1.9 | 7.6 | ± | 2.4 |
| 2.5-5 | 3.4 | ± | 1.6 | 6.1 | ± | 2.8 | 9.4 | ± | 3.3 | 13.6 | ± | 3.6 | 6.6 | ± | 2.6 | 4.1 | ± | 2.2 | 11.9 | ± | 3.4 | 4.8 | ± | 2.2 |
| 5-10 | 23.1 | ± | 8.0 | 19.3 | ± | 8.4 | 15.1 | ± | 4.1 | 15.1 | ± | 4.2 | 16.3 | ± | 4.5 | 19.2 | ± | 7.3 | 19.9 | ± | 4.1 | 10.8 | ± | 3.0 |
| 10-20 | 28.9 | ± | 14.7 | 33.7 | ± | 6.8 | 28.7 | ± | 5.4 | 30.0 | ± | 3.3 | 23.4 | ± | 6.1 | 15.1 | ± | 8.4 | 14.7 | ± | 5.7 | 21.4 | ± | 7.2 |
| 20-40 | 29.8 | ± | 13.7 | 26.8 | ± | 12.2 | 24.1 | ± | 6.8 | 20.3 | ± | 6.8 | 36.5 | ± | 8.9 | 17.0 | ± | 7.8 | 30.4 | ± | 11.7 | 33.9 | ± | 11.1 |
| 40-∞ | 0.0 | ± | 0.0 | 0.0 | ± | 0.0 | 4.5 | ± | 4.5 | 0.0 | ± | 0.0 | 0.0 | ± | 0.0 | 25.3 | ± | 11.8<sup>β</sup> | 0.0 | ± | 0.0 | 6.3 | ± | 6.3 |
| Number of episodes (± SEM) of wakefulness in various wake episode lengths during the ZT3-6 period following CNO administration as compared with control administration |  |  |  |  |  |  |  |  |  |  |  |  |  |  |  |  |  |  |  |  |  |  |  |  |
| Episode length (min) | Control |  |  | 1<sup>st</sup> 2 administrations |  |  | 1<sup>st</sup> month |  |  | 2<sup>nd</sup> month |  |  | 3<sup>rd</sup> month |  |  | 4<sup>th</sup> month |  |  | 5<sup>th</sup> month |  |  | 6<sup>th</sup> month |  |  |
|  | Avg | ± | SEM | Avg | ± | SEM | Avg | ± | SEM | Avg | ± | SEM | Avg | ± | SEM | Avg | ± | SEM | Avg | ± | SEM | Avg | ± | SEM |
| 0.1-0.5 | 19.8 | ± | 1.6 | 19.5 | ± | 0.9 | 19.0 | ± | 1.4 | 22.3 | ± | 2.5 | 19.7 | ± | 1.6 | 20.1 | ± | 1.1 | 14.6 | ± | 1.7 <sup>γ</sup> | 20.3 | ± | 2.0 |
| 0.5-1 | 1.2 | ± | 0.3 | 1.5 | ± | 0.3 | 2.8 | ± | 1.4 | 2.8 | ± | 0.5 | 2.6 | ± | 0.5 | 2.8 | ± | 1.0 | 5.0 | ± | 1.6 <sup>γ</sup> | 2.9 | ± | 0.6 |
| 1-2.5 | 1.0 | ± | 0.4 | 0.6 | ± | 0.2 | 1.1 | ± | 0.4 | 0.9 | ± | 0.4 | 0.8 | ± | 0.2 | 1.3 | ± | 0.7 | 2.5 | ± | 0.6 | 2.7 | ± | 0.7 |
| 2.5-5 | 0.5 | ± | 0.2 | 0.6 | ± | 0.3 | 1.1 | ± | 0.4 | 1.3 | ± | 0.2 | 0.8 | ± | 0.4 | 0.4 | ± | 0.2 | 1.6 | ± | 0.4 | 0.8 | ± | 0.3 |
| 5-10 | 1.7 | ± | 0.5 | 1.3 | ± | 0.5 | 0.9 | ± | 0.2 | 1.1 | ± | 0.3 | 1.1 | ± | 0.2 | 1.3 | ± | 0.4 | 1.6 | ± | 0.3 | 0.9 | ± | 0.3 |
| 10-20 | 0.8 | ± | 0.4 | 0.9 | ± | 0.2 | 0.9 | ± | 0.2 | 1.0 | ± | 0.2 | 0.9 | ± | 0.3 | 0.5 | ± | 0.3 | 0.5 | ± | 0.2 | 0.9 | ± | 0.3 |
| 20-40 | 0.5 | ± | 0.2 | 0.5 | ± | 0.2 | 0.5 | ± | 0.1 | 0.4 | ± | 0.2 | 0.8 | ± | 0.2 | 0.3 | ± | 0.2 | 0.7 | ± | 0.2 | 0.9 | ± | 0.3 |
| 40-∞ | 0.0 | ± | 0.0 | 0.0 | ± | 0.0 | 0.1 | ± | 0.1 | 0.0 | ± | 0.0 | 0.0 | ± | 0.0 | 0.3 | ± | 0.2 | 0.0 | ± | 0.0 | 0.1 | ± | 0.1 |

**Supplementary Table S20:** Percentage (±SEM) of **wakefulness** amount in various wake episode lengths to the total amount of wakefulness and number of episodes (± SEM) of wakefulness in various wake episode lengths during the **ZT3-6** periods in control mice following **DCZ** administration as compared with control administration. N = 6 mice. α: p < 0.05, β: p < 0.01, δ: p < 0.001, γ: p < 0.0001, two-way ANOVA followed by a post hoc Bonferroni test.

| Percentage (±SEM) of wakefulness amount in various wake episode lengths to the total amount of wakefulness during the ZT3-6 period following DCZ administration as compared with control administration |  |  |  |  |  |  |  |  |  |  |  |  |  |  |  |  |  |  |  |  |  |  |  |  |
| --- | --- | --- | --- | --- | --- | --- | --- | --- | --- | --- | --- | --- | --- | --- | --- | --- | --- | --- | --- | --- | --- | --- | --- | --- |
| Episode length (min) | Control |  |  | 1<sup>st</sup> 2 administrations |  |  | 1<sup>st</sup> month |  |  | 2<sup>nd</sup> month |  |  | 3<sup>rd</sup> month |  |  | 4<sup>th</sup> month |  |  | 5<sup>th</sup> month |  |  | 6<sup>th</sup> month |  |  |
|  | Avg | ± | SEM | Avg | ± | SEM | Avg | ± | SEM | Avg | ± | SEM | Avg | ± | SEM | Avg | ± | SEM | Avg | ± | SEM | Avg | ± | SEM |
| 0.1-0.5 | 9.6 | ± | 1.3 | 8.5 | ± | 1.3 | 11.5 | ± | 1.6 | 10.3 | ± | 1.5 | 10.2 | ± | 2.3 | 10.3 | ± | 1.1 | 10.8 | ± | 1.5 | 11.4 | ± | 1.1 |
| 0.5-1 | 2.1 | ± | 0.6 | 2.7 | ± | 0.7 | 4.6 | ± | 1.0 | 4.5 | ± | 0.9 | 4.4 | ± | 1.0 | 4.2 | ± | 0.7 | 4.1 | ± | 0.8 | 6.6 | ± | 2.4 |
| 1-2.5 | 3.1 | ± | 1.2 | 3.5 | ± | 1.8 | 6.8 | ± | 1.8 | 2.4 | ± | 1.2 | 2.6 | ± | 1.1 | 4.2 | ± | 1.2 | 5.2 | ± | 2.4 | 6.0 | ± | 1.9 |
| 2.5-5 | 3.4 | ± | 1.6 | 0.5 | ± | 0.5 | 12.1 | ± | 4.0 | 8.3 | ± | 2.8 | 7.5 | ± | 2.9 | 12.8 | ± | 2.8 | 8.0 | ± | 1.3 | 6.0 | ± | 2.0 |
| 5-10 | 23.1 | ± | 8.0 | 15.7 | ± | 3.2 | 11.3 | ± | 2.3 | 8.4 | ± | 3.4 | 20.4 | ± | 4.5 | 8.4 | ± | 3.1 | 16.2 | ± | 7.4 | 10.5 | ± | 3.8 |
| 10-20 | 28.9 | ± | 14.7 | 39.6 | ± | 9.0 | 33.6 | ± | 6.2 | 43.0 | ± | 8.2 | 17.6 | ± | 7.4 | 31.8 | ± | 7.3 | 34.0 | ± | 8.1 | 25.6 | ± | 12.5 |
| 20-40 | 29.8 | ± | 13.7 | 29.5 | ± | 11.1 | 20.0 | ± | 9.1 | 23.1 | ± | 8.4 | 23.3 | ± | 7.9 | 28.2 | ± | 9.5 | 21.7 | ± | 10.3 | 27.7 | ± | 8.7 |
| 40-∞ | 0.0 | ± | 0.0 | 0.0 | ± | 0.0 | 0.0 | ± | 0.0 | 0.0 | ± | 0.0 | 14.0 | ± | 8.8 | 0.0 | ± | 0.0 | 0.0 | ± | 0.0 | 6.3 | ± | 6.3 |
| Number of episodes (± SEM) of wakefulness in various wake episode lengths during the ZT3-6 period following DCZ administration as compared with control administration |  |  |  |  |  |  |  |  |  |  |  |  |  |  |  |  |  |  |  |  |  |  |  |  |
| Episode length (min) | Control |  |  | 1<sup>st</sup> 2 administrations |  |  | 1<sup>st</sup> month |  |  | 2<sup>nd</sup> month |  |  | 3<sup>rd</sup> month |  |  | 4<sup>th</sup> month |  |  | 5<sup>th</sup> month |  |  | 6<sup>th</sup> month |  |  |
|  | Avg | ± | SEM | Avg | ± | SEM | Avg | ± | SEM | Avg | ± | SEM | Avg | ± | SEM | Avg | ± | SEM | Avg | ± | SEM | Avg | ± | SEM |
| 0.1-0.5 | 19.8 | ± | 1.6 | 16.2 | ± | 1.5 **δ** | 17.8 | ± | 2.1 | 19.2 | ± | 1.5 | 18.8 | ± | 2.3 | 16.7 | ± | 1.9 **β** | 19.4 | ± | 1.6 | 20.2 | ± | 1.4 |
| 0.5-1 | 1.2 | ± | 0.3 | 1.5 | ± | 0.3 | 2.7 | ± | 0.6 | 2.8 | ± | 0.4 | 3.2 | ± | 0.8 | 2.7 | ± | 0.5 | 2.8 | ± | 0.6 | 4.7 | ± | 1.9 **β** |
| 1-2.5 | 1.0 | ± | 0.4 | 1.0 | ± | 0.5 | 1.9 | ± | 0.5 | 0.8 | ± | 0.4 | 0.8 | ± | 0.4 | 1.3 | ± | 0.6 | 1.8 | ± | 0.9 | 2.0 | ± | 0.6 |
| 2.5-5 | 0.5 | ± | 0.2 | 0.1 | ± | 0.1 | 1.5 | ± | 0.5 | 1.0 | ± | 0.3 | 1.0 | ± | 0.5 | 1.4 | ± | 0.2 | 1.1 | ± | 0.2 | 0.8 | ± | 0.3 |
| 5-10 | 1.7 | ± | 0.5 | 1.0 | ± | 0.1 | 0.8 | ± | 0.2 | 0.6 | ± | 0.2 | 1.5 | ± | 0.3 | 0.6 | ± | 0.2 | 1.2 | ± | 0.5 | 0.9 | ± | 0.3 |
| 10-20 | 0.8 | ± | 0.4 | 1.2 | ± | 0.2 | 1.0 | ± | 0.2 | 1.5 | ± | 0.4 | 0.6 | ± | 0.2 | 1.3 | ± | 0.3 | 1.3 | ± | 0.3 | 0.8 | ± | 0.3 |
| 20-40 | 0.5 | ± | 0.2 | 0.8 | ± | 0.3 | 0.3 | ± | 0.2 | 0.5 | ± | 0.2 | 0.6 | ± | 0.2 | 0.6 | ± | 0.2 | 0.4 | ± | 0.2 | 0.6 | ± | 0.2 |
| 40-∞ | 0.0 | ± | 0.0 | 0.0 | ± | 0.0 | 0.0 | ± | 0.0 | 0.0 | ± | 0.0 | 0.2 | ± | 0.1 | 0.0 | ± | 0.0 | 0.0 | ± | 0.0 | 0.1 | ± | 0.1 |

**Supplementary Table S21:**
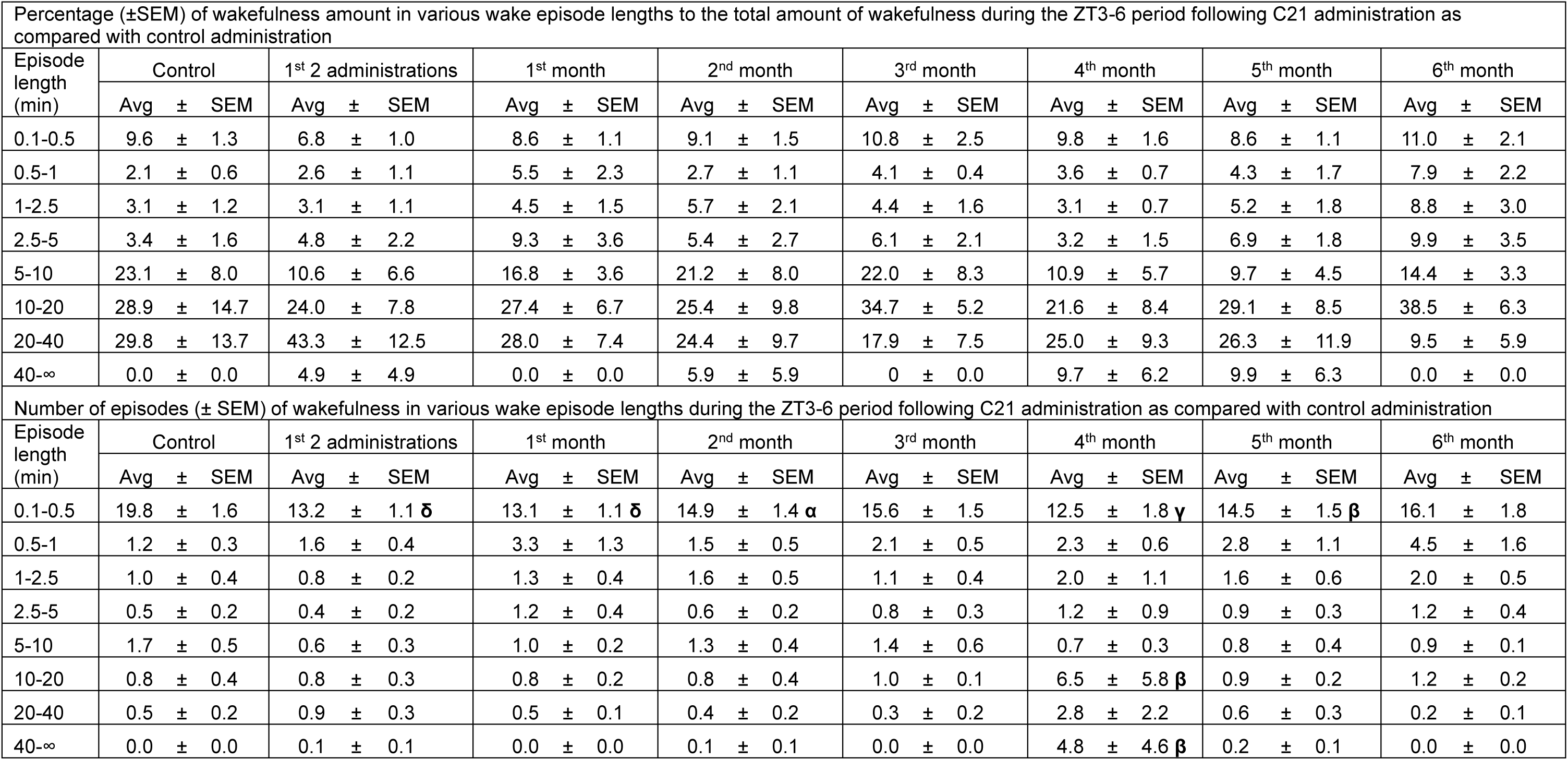
Percentage (±SEM) of **wakefulness** amount in various wake episode lengths to the total amount of wakefulness and number of episodes (± SEM) of wakefulness in various wake episode lengths during the **ZT3-6** periods in control mice following **C21** administration as compared with control administration. N = 6 mice. α: p < 0.05, β: p < 0.01, δ: p < 0.001, γ: p < 0.0001, two-way ANOVA followed by a post hoc Bonferroni test.

| Percentage (±SEM) of wakefulness amount in various wake episode lengths to the total amount of wakefulness during the ZT3-6 period following C21 administration as compared with control administration |  |  |  |  |  |  |  |  |  |  |  |  |  |  |  |  |  |  |  |  |  |  |  |  |
| --- | --- | --- | --- | --- | --- | --- | --- | --- | --- | --- | --- | --- | --- | --- | --- | --- | --- | --- | --- | --- | --- | --- | --- | --- |
| Episode length (min) | Control |  |  | 1<sup>st</sup> 2 administrations |  |  | 1<sup>st</sup> month |  |  | 2<sup>nd</sup> month |  |  | 3<sup>rd</sup> month |  |  | 4<sup>th</sup> month |  |  | 5<sup>th</sup> month |  |  | 6<sup>th</sup> month |  |  |
|  | Avg | ± | SEM | Avg | ± | SEM | Avg | ± | SEM | Avg | ± | SEM | Avg | ± | SEM | Avg | ± | SEM | Avg | ± | SEM | Avg | ± | SEM |
| 0.1-0.5 | 9.6 | ± | 1.3 | 6.8 | ± | 1.0 | 8.6 | ± | 1.1 | 9.1 | ± | 1.5 | 10.8 | ± | 2.5 | 9.8 | ± | 1.6 | 8.6 | ± | 1.1 | 11.0 | ± | 2.1 |
| 0.5-1 | 2.1 | ± | 0.6 | 2.6 | ± | 1.1 | 5.5 | ± | 2.3 | 2.7 | ± | 1.1 | 4.1 | ± | 0.4 | 3.6 | ± | 0.7 | 4.3 | ± | 1.7 | 7.9 | ± | 2.2 |
| 1-2.5 | 3.1 | ± | 1.2 | 3.1 | ± | 1.1 | 4.5 | ± | 1.5 | 5.7 | ± | 2.1 | 4.4 | ± | 1.6 | 3.1 | ± | 0.7 | 5.2 | ± | 1.8 | 8.8 | ± | 3.0 |
| 2.5-5 | 3.4 | ± | 1.6 | 4.8 | ± | 2.2 | 9.3 | ± | 3.6 | 5.4 | ± | 2.7 | 6.1 | ± | 2.1 | 3.2 | ± | 1.5 | 6.9 | ± | 1.8 | 9.9 | ± | 3.5 |
| 5-10 | 23.1 | ± | 8.0 | 10.6 | ± | 6.6 | 16.8 | ± | 3.6 | 21.2 | ± | 8.0 | 22.0 | ± | 8.3 | 10.9 | ± | 5.7 | 9.7 | ± | 4.5 | 14.4 | ± | 3.3 |
| 10-20 | 28.9 | ± | 14.7 | 24.0 | ± | 7.8 | 27.4 | ± | 6.7 | 25.4 | ± | 9.8 | 34.7 | ± | 5.2 | 21.6 | ± | 8.4 | 29.1 | ± | 8.5 | 38.5 | ± | 6.3 |
| 20-40 | 29.8 | ± | 13.7 | 43.3 | ± | 12.5 | 28.0 | ± | 7.4 | 24.4 | ± | 9.7 | 17.9 | ± | 7.5 | 25.0 | ± | 9.3 | 26.3 | ± | 11.9 | 9.5 | ± | 5.9 |
| 40-∞ | 0.0 | ± | 0.0 | 4.9 | ± | 4.9 | 0.0 | ± | 0.0 | 5.9 | ± | 5.9 | 0 | ± | 0.0 | 9.7 | ± | 6.2 | 9.9 | ± | 6.3 | 0.0 | ± | 0.0 |
| Number of episodes (± SEM) of wakefulness in various wake episode lengths during the ZT3-6 period following C21 administration as compared with control administration |  |  |  |  |  |  |  |  |  |  |  |  |  |  |  |  |  |  |  |  |  |  |  |  |
| Episode length (min) | Control |  |  | 1<sup>st</sup> 2 administrations |  |  | 1<sup>st</sup> month |  |  | 2<sup>nd</sup> month |  |  | 3<sup>rd</sup> month |  |  | 4<sup>th</sup> month |  |  | 5<sup>th</sup> month |  |  | 6<sup>th</sup> month |  |  |
|  | Avg | ± | SEM | Avg | ± | SEM | Avg | ± | SEM | Avg | ± | SEM | Avg | ± | SEM | Avg | ± | SEM | Avg | ± | SEM | Avg | ± | SEM |
| 0.1-0.5 | 19.8 | ± | 1.6 | 13.2 | ± | 1.1 δ | 13.1 | ± | 1.1 δ | 14.9 | ± | 1.4 α | 15.6 | ± | 1.5 | 12.5 | ± | 1.8 γ | 14.5 | ± | 1.5 β | 16.1 | ± | 1.8 |
| 0.5-1 | 1.2 | ± | 0.3 | 1.6 | ± | 0.4 | 3.3 | ± | 1.3 | 1.5 | ± | 0.5 | 2.1 | ± | 0.5 | 2.3 | ± | 0.6 | 2.8 | ± | 1.1 | 4.5 | ± | 1.6 |
| 1-2.5 | 1.0 | ± | 0.4 | 0.8 | ± | 0.2 | 1.3 | ± | 0.4 | 1.6 | ± | 0.5 | 1.1 | ± | 0.4 | 2.0 | ± | 1.1 | 1.6 | ± | 0.6 | 2.0 | ± | 0.5 |
| 2.5-5 | 0.5 | ± | 0.2 | 0.4 | ± | 0.2 | 1.2 | ± | 0.4 | 0.6 | ± | 0.2 | 0.8 | ± | 0.3 | 1.2 | ± | 0.9 | 0.9 | ± | 0.3 | 1.2 | ± | 0.4 |
| 5-10 | 1.7 | ± | 0.5 | 0.6 | ± | 0.3 | 1.0 | ± | 0.2 | 1.3 | ± | 0.4 | 1.4 | ± | 0.6 | 0.7 | ± | 0.3 | 0.8 | ± | 0.4 | 0.9 | ± | 0.1 |
| 10-20 | 0.8 | ± | 0.4 | 0.8 | ± | 0.3 | 0.8 | ± | 0.2 | 0.8 | ± | 0.4 | 1.0 | ± | 0.1 | 6.5 | ± | 5.8 β | 0.9 | ± | 0.2 | 1.2 | ± | 0.2 |
| 20-40 | 0.5 | ± | 0.2 | 0.9 | ± | 0.3 | 0.5 | ± | 0.1 | 0.4 | ± | 0.2 | 0.3 | ± | 0.2 | 2.8 | ± | 2.2 | 0.6 | ± | 0.3 | 0.2 | ± | 0.1 |
| 40-∞ | 0.0 | ± | 0.0 | 0.1 | ± | 0.1 | 0.0 | ± | 0.0 | 0.1 | ± | 0.1 | 0.0 | ± | 0.0 | 4.8 | ± | 4.6 β | 0.2 | ± | 0.1 | 0.0 | ± | 0.0 |

**Supplementary Table S22:**
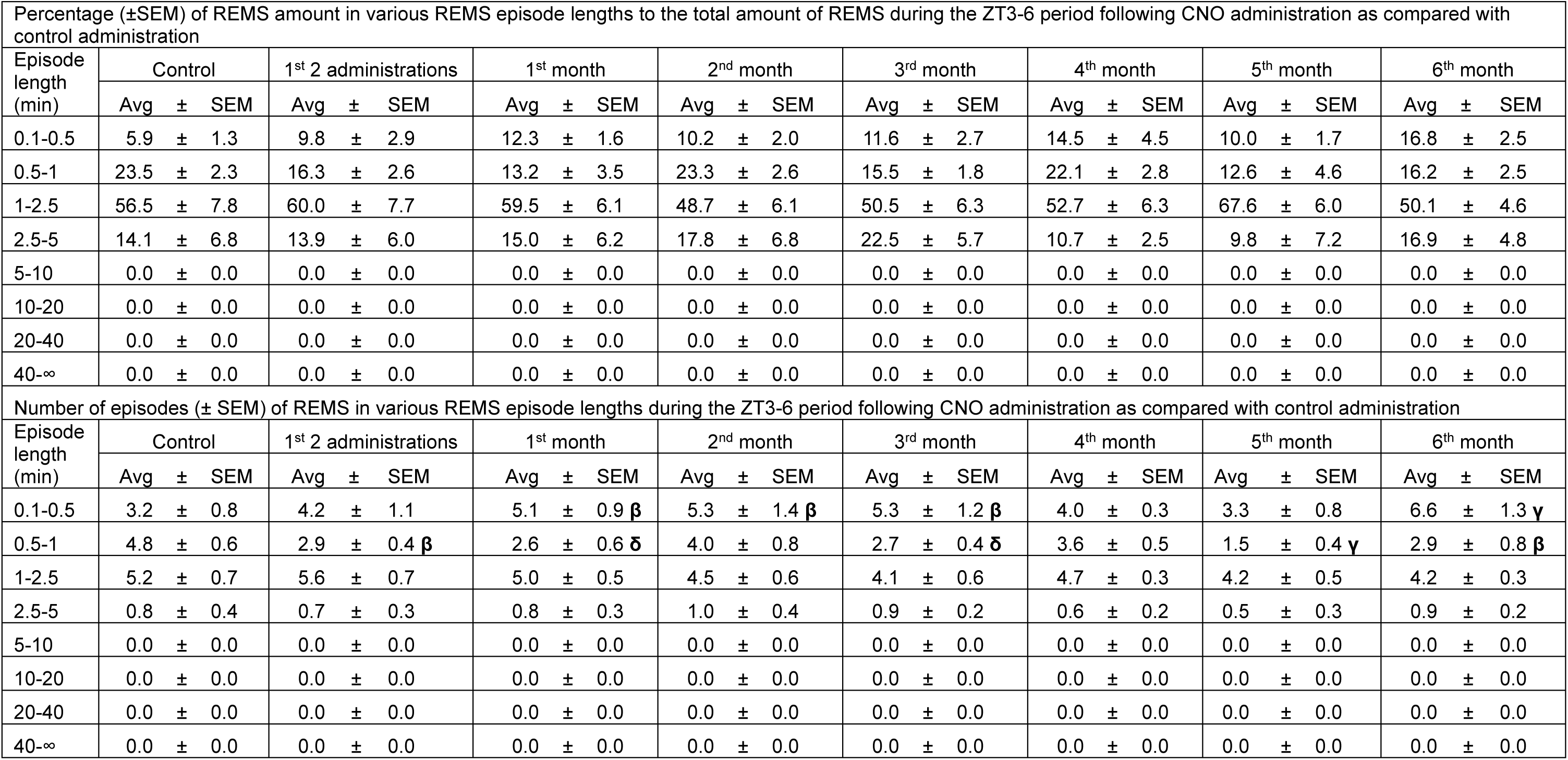
Percentage (±SEM) of **REMS** amount in various REMS episode lengths to the total amount of REMS and number of episodes (± SEM) of REMS in various REMS episode lengths during the **ZT3-6** periods in control mice following **CNO** administration as compared with control administration. N = 6 mice. α: p < 0.05, β: p < 0.01, δ: p < 0.001, γ: p < 0.0001, two-way ANOVA followed by a post hoc Bonferroni test.

**Supplementary Table S23:** Percentage (±SEM) of **REMS** amount in various REMS episode lengths to the total amount of REMS and number of episodes (± SEM) of REMS in various REMS episode lengths during the **ZT3-6** periods in control mice following **DCZ** administration as compared with control administration. N = 6 mice. α: p < 0.05, β: p < 0.01, δ: p < 0.001, γ: p < 0.0001, two-way ANOVA followed by a post hoc Bonferroni test.

| Percentage (±SEM) of REMS amount in various REMS episode lengths to the total amount of REMS during the ZT3-6 period following DCZ administration as compared with control administration |  |  |  |  |  |  |  |  |  |  |  |  |  |  |  |  |  |  |  |  |  |  |  |  |
| --- | --- | --- | --- | --- | --- | --- | --- | --- | --- | --- | --- | --- | --- | --- | --- | --- | --- | --- | --- | --- | --- | --- | --- | --- |
| Episode length (min) | Control |  |  | 1<sup>st</sup> 2 administrations |  |  | 1<sup>st</sup> month |  |  | 2<sup>nd</sup> month |  |  | 3<sup>rd</sup> month |  |  | 4<sup>th</sup> month |  |  | 5<sup>th</sup> month |  |  | 6<sup>th</sup> month |  |  |
|  | Avg | ± | SEM | Avg | ± | SEM | Avg | ± | SEM | Avg | ± | SEM | Avg | ± | SEM | Avg | ± | SEM | Avg | ± | SEM | Avg | ± | SEM |
| 0.1-0.5 | 5.9 | ± | 1.3 | 9.7 | ± | 2.4 | 12.0 | ± | 2.1 | 8.8 | ± | 2.8 | 11.7 | ± | 2.8 | 19.7 | ± | 8.2 α | 15.6 | ± | 2.4 | 11.1 | ± | 3.5 |
| 0.5-1 | 23.5 | ± | 2.3 | 18.0 | ± | 1.5 | 18.3 | ± | 4.2 | 15.7 | ± | 3.2 | 13.2 | ± | 4.8 | 13.9 | ± | 3.9 | 17.3 | ± | 4.7 | 18.5 | ± | 3.8 |
| 1-2.5 | 56.5 | ± | 7.8 | 60.3 | ± | 4.7 | 49.9 | ± | 5.9 | 62.4 | ± | 4.6 | 62.1 | ± | 4.8 | 50.3 | ± | 5.1 | 42.5 | ± | 4.4 α | 46.8 | ± | 9.9 |
| 2.5-5 | 14.1 | ± | 6.8 | 11.9 | ± | 4.6 | 19.8 | ± | 5.3 | 13.1 | ± | 5.9 | 13.0 | ± | 4.4 | 16.2 | ± | 5.0 | 24.6 | ± | 8.6 | 23.5 | ± | 7.8 |
| 5-10 | 0.0 | ± | 0.0 | 0.0 | ± | 0.0 | 0.0 | ± | 0.0 | 0.0 | ± | 0.0 | 0.0 | ± | 0.0 | 0.0 | ± | 0.0 | 0.0 | ± | 0.0 | 0.0 | ± | 0.0 |
| 10-20 | 0.0 | ± | 0.0 | 0.0 | ± | 0.0 | 0.0 | ± | 0.0 | 0.0 | ± | 0.0 | 0.0 | ± | 0.0 | 0.0 | ± | 0.0 | 0.0 | ± | 0.0 | 0.0 | ± | 0.0 |
| 20-40 | 0.0 | ± | 0.0 | 0.0 | ± | 0.0 | 0.0 | ± | 0.0 | 0.0 | ± | 0.0 | 0.0 | ± | 0.0 | 0.0 | ± | 0.0 | 0.0 | ± | 0.0 | 0.0 | ± | 0.0 |
| 40-∞ | 0.0 | ± | 0.0 | 0.0 | ± | 0.0 | 0.0 | ± | 0.0 | 0.0 | ± | 0.0 | 0.0 | ± | 0.0 | 0.0 | ± | 0.0 | 0.0 | ± | 0.0 | 0.0 | ± | 0.0 |
| Number of episodes (± SEM) of REMS in various REMS episode lengths during the ZT3-6 period following DCZ administration as compared with control administration |  |  |  |  |  |  |  |  |  |  |  |  |  |  |  |  |  |  |  |  |  |  |  |  |
| Episode length (min) | Control |  |  | 1<sup>st</sup> 2 administrations |  |  | 1<sup>st</sup> month |  |  | 2<sup>nd</sup> month |  |  | 3<sup>rd</sup> month |  |  | 4<sup>th</sup> month |  |  | 5<sup>th</sup> month |  |  | 6<sup>th</sup> month |  |  |
|  | Avg | ± | SEM | Avg | ± | SEM | Avg | ± | SEM | Avg | ± | SEM | Avg | ± | SEM | Avg | ± | SEM | Avg | ± | SEM | Avg | ± | SEM |
| 0.1-0.5 | 3.2 | ± | 0.8 | 3.8 | ± | 0.8 | 4.8 | ± | 1.1 α | 3.5 | ± | 0.8 | 4.1 | ± | 1.0 | 4.3 | ± | 0.9 | 4.7 | ± | 0.8 α | 3.8 | ± | 0.9 |
| 0.5-1 | 4.8 | ± | 0.6 | 3.2 | ± | 0.3 β | 3.1 | ± | 0.9 β | 2.5 | ± | 0.6 γ | 2.3 | ± | 0.8 γ | 2.4 | ± | 0.7 γ | 2.3 | ± | 0.6 γ | 2.8 | ± | 0.4 β |
| 1-2.5 | 5.2 | ± | 0.7 | 5.3 | ± | 0.4 | 3.8 | ± | 0.3 α | 5.1 | ± | 0.7 | 4.3 | ± | 0.4 | 3.6 | ± | 0.7 α | 2.5 | ± | 0.4 γ | 3.7 | ± | 0.8 α |
| 2.5-5 | 0.8 | ± | 0.4 | 0.6 | ± | 0.2 | 0.9 | ± | 0.2 | 0.6 | ± | 0.2 | 0.5 | ± | 0.2 | 0.8 | ± | 0.2 | 1.0 | ± | 0.4 | 1.0 | ± | 0.4 |
| 5-10 | 0.0 | ± | 0.0 | 0.0 | ± | 0.0 | 0.0 | ± | 0.0 | 0.0 | ± | 0.0 | 0.0 | ± | 0.0 | 0.0 | ± | 0.0 | 0.0 | ± | 0.0 | 0.0 | ± | 0.0 |
| 10-20 | 0.0 | ± | 0.0 | 0.0 | ± | 0.0 | 0.0 | ± | 0.0 | 0.0 | ± | 0.0 | 0.0 | ± | 0.0 | 0.0 | ± | 0.0 | 0.0 | ± | 0.0 | 0.0 | ± | 0.0 |
| 20-40 | 0.0 | ± | 0.0 | 0.0 | ± | 0.0 | 0.0 | ± | 0.0 | 0.0 | ± | 0.0 | 0.0 | ± | 0.0 | 0.0 | ± | 0.0 | 0.0 | ± | 0.0 | 0.0 | ± | 0.0 |
| 40-∞ | 0.0 | ± | 0.0 | 0.0 | ± | 0.0 | 0.0 | ± | 0.0 | 0.0 | ± | 0.0 | 0.0 | ± | 0.0 | 0.0 | ± | 0.0 | 0.0 | ± | 0.0 | 0.0 | ± | 0.0 |

**Supplementary Table S24:**
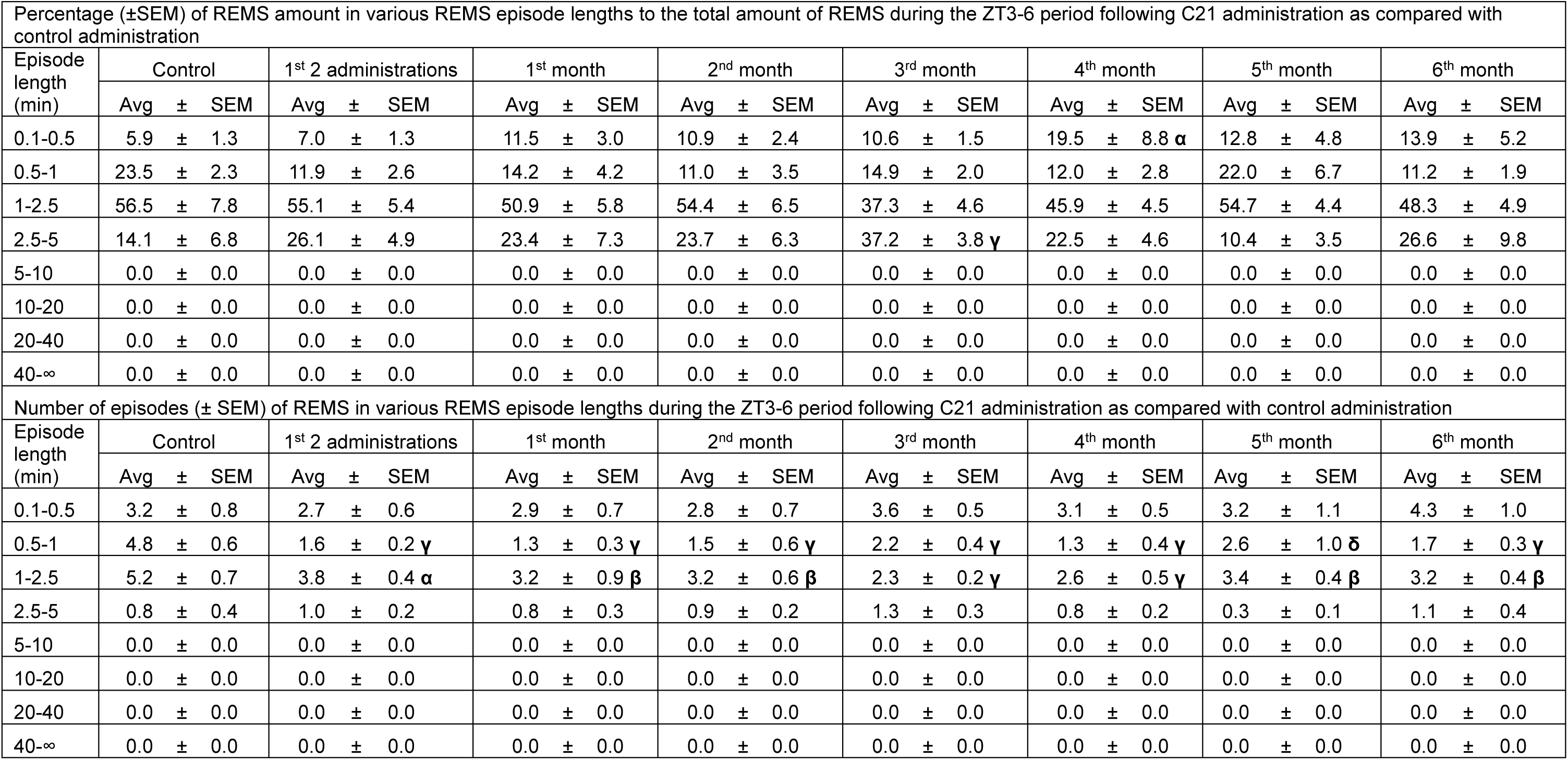
Percentage (±SEM) of **REMS** amount in various REMS episode lengths to the total amount of REMS and number of episodes (± SEM) of REMS in various REMS episode lengths during the **ZT3-6** periods in control mice following **C21** administration as compared with control administration. N = 6 mice. α: p < 0.05, β: p < 0.01, δ: p < 0.001, γ: p < 0.0001, two-way ANOVA followed by a post hoc Bonferroni test.

